# Molecular topography of an entire nervous system

**DOI:** 10.1101/2020.12.15.422897

**Authors:** Seth R Taylor, Gabriel Santpere, Alexis Weinreb, Alec Barrett, Molly B. Reilly, Chuan Xu, Erdem Varol, Panos Oikonomou, Lori Glenwinkel, Rebecca McWhirter, Abigail Poff, Manasa Basavaraju, Ibnul Rafi, Eviatar Yemini, Steven J Cook, Alexander Abrams, Berta Vidal, Cyril Cros, Saeed Tavazoie, Nenad Sestan, Marc Hammarlund, Oliver Hobert, David M. Miller

## Abstract

Nervous systems are constructed from a deep repertoire of neuron types but the underlying gene expression programs that specify individual neuron identities are poorly understood. To address this deficit, we have produced an expression profile of all 302 neurons of the *C. elegans* nervous system that matches the single cell resolution of its anatomy and wiring diagram. Our results suggest that individual neuron classes can be solely identified by combinatorial expression of specific gene families. For example, each neuron class expresses unique codes of ∼23 neuropeptide-encoding genes and ∼36 neuropeptide receptors thus pointing to an expansive “wireless” signaling network. To demonstrate the utility of this uniquely comprehensive gene expression catalog, we used computational approaches to (1) identify cis-regulatory elements for neuron-specific gene expression across the nervous system and (2) reveal adhesion proteins with potential roles in synaptic specificity and process placement. These data are available at cengen.org and can be interrogated at the web application CengenApp. We expect that this neuron-specific directory of gene expression will spur investigations of underlying mechanisms that define anatomy, connectivity and function throughout the *C. elegans* nervous system.

## INTRODUCTION

Neurons share the common functions of receiving, processing, propagating and transmitting information. Yet neurons also adopt a remarkable variety of distinct types, each with unique features that define its role in nervous system function. Because genetic programs likely specify individual neuron types, the goal of building a molecular model of the brain requires a gene expression map at single-cell resolution. Although new profiling methods have been used to catalog diverse neuron types in a variety of organisms (Adorjan et al., 2019; Poulin et al., 2016; Tasic et al., 2016; Zeisel et al., 2015; Zhu et al., 2018), their complexity has limited these approaches to specific subset of neurons. Moreover, incomplete knowledge of the lineage, anatomy and synaptic wiring of complex nervous systems has constrained the opportunity to correlate functional and anatomical properties with molecular signatures.

To understand the global relationship between gene expression and neuronal anatomy and function, we have produced single cell RNA-Seq (scRNA-Seq) profiles for all individual neurons in an entire nervous system, that of the *C. elegans* hermaphrodite. The complete anatomy and wiring diagram of the *C. elegans* nervous system have been reconstructed from electron micrographs of serial sections (Albertson and Thomson, 1976; Brittin et al., 2020; Cook et al., 2019; White et al., 1986; Witvliet et al., 2020). This approach identified 118 anatomically distinct classes among the 302 neurons in the mature hermaphrodite nervous system. We established the *C. elegans* Neuronal Gene Expression Map & Network (CeNGEN) consortium (Hammarlund et al., 2018) to generate transcriptional profiles of each neuron class, thereby bridging the gap between *C. elegans* neuroanatomy and the genetic blueprint that defines it. We used fluorescence activated cell sorting (FACS) to isolate neurons from L4 stage larvae for single cell RNA-sequencing. By the L4 stage, the entire nervous system has been generated and most neurons have terminally differentiated. Our approach generated profiles of 70,296 neurons. Unbiased clustering resolved these cells into distinct groups that included each of the 118 canonical neuron classes, confirming that gene expression is tightly linked to anatomy and function. To our knowledge, these data are the first to describe neuron-specific gene expression across an entire nervous system.

Additional analyses allowed us to probe the robustness of our data and to discover new biology. First, subtle differences in wiring and reporter gene expression within these anatomically defined neuron classes predicted a substantially larger array of neuron types each with a unique molecular profile (Hobert et al., 2016). We have substantiated this idea by detecting distinct single cell profiles that correspond to subtypes of neurons within at least 10 of the known neuron classes. Second, we used these data to map expression of key neuronal gene families for neurotransmitters, receptors, ion channels and transcription factors across the *C. elegans* nervous system thereby identifying unique aspects of expression in each family that may support the diverse functions of different neuron types. Among the striking findings or our approach is the discovery that every neuron class is defined by distinct combinations of neuropeptide-encoding genes and neuropeptide receptors which could be indicative of unique roles for each type of neuron in sending and receiving signals. Third, for 8 selected neuron types, we used bulk RNA-sequencing to confirm scRNA-Seq profiles and also to expand these neuron-specific fingerprints to include non-coding RNAs and whole transcript coverage that documents alternatively spliced isoforms. Fourth, we compared our results for *C. elegans* cell types to profiles from other organisms, and found striking concurrences in other species. Fifth, we utilized our global map of gene expression across the entire nervous system to identify an expansive catalog of DNA and RNA sequence motifs that are correlated with cohorts of co-regulated genes. Finally, we used computational approaches to identify cell adhesion molecules associated with neuron-specific synapses and bundling. Together, our results provide a uniquely comprehensive link between neuron-specific gene expression and the structure and function of an entire nervous system. We expect that these data sets and the tools that we have developed for interrogating them will power new investigations into the genetic basis of neuronal connectivity and function.

## RESULTS AND DISCUSSION

### Single cell RNA-Sequencing identifies all known neuron classes in the mature *C. elegans* nervous system

To profile the entire *C. elegans* nervous system, we isolated neurons from L4 larvae, a time point by which all neuron types have been generated (Sulston and Horvitz, 1977). In our initial experiment, we used FACS to isolate neurons from a pan-neural marker strain (*otIs355*) which expresses nuclear-localized TagRFP in all neuron types except CAN (Stefanakis et al., 2015) (Figure S1A, C). Although this approach yielded a variety of recognizable neuron types, many neuron classes were either underrepresented or absent (Figure S1C). To overcome this limitation, we isolated cells from a series of fluorescent marker strains that labeled distinct subsets of neurons, including CAN (Figure 1A, Table S2). With this approach, we produced a combined dataset of 100,955 single cell transcriptomes. Application of the Uniform Manifold Approximation and Projection (UMAP) (Becht et al., 2019; McInnes et al., 2018) dimensional reduction algorithm effectively segregated most of these cells into distinct groups in the UMAP space (Figure S2A). We clustered cells in the UMAP using the Louvain algorithm (Blondel et al., 2008). We detected a median of 928 UMIs/cell and 328 genes/cell across the entire dataset.

**Figure 1.**
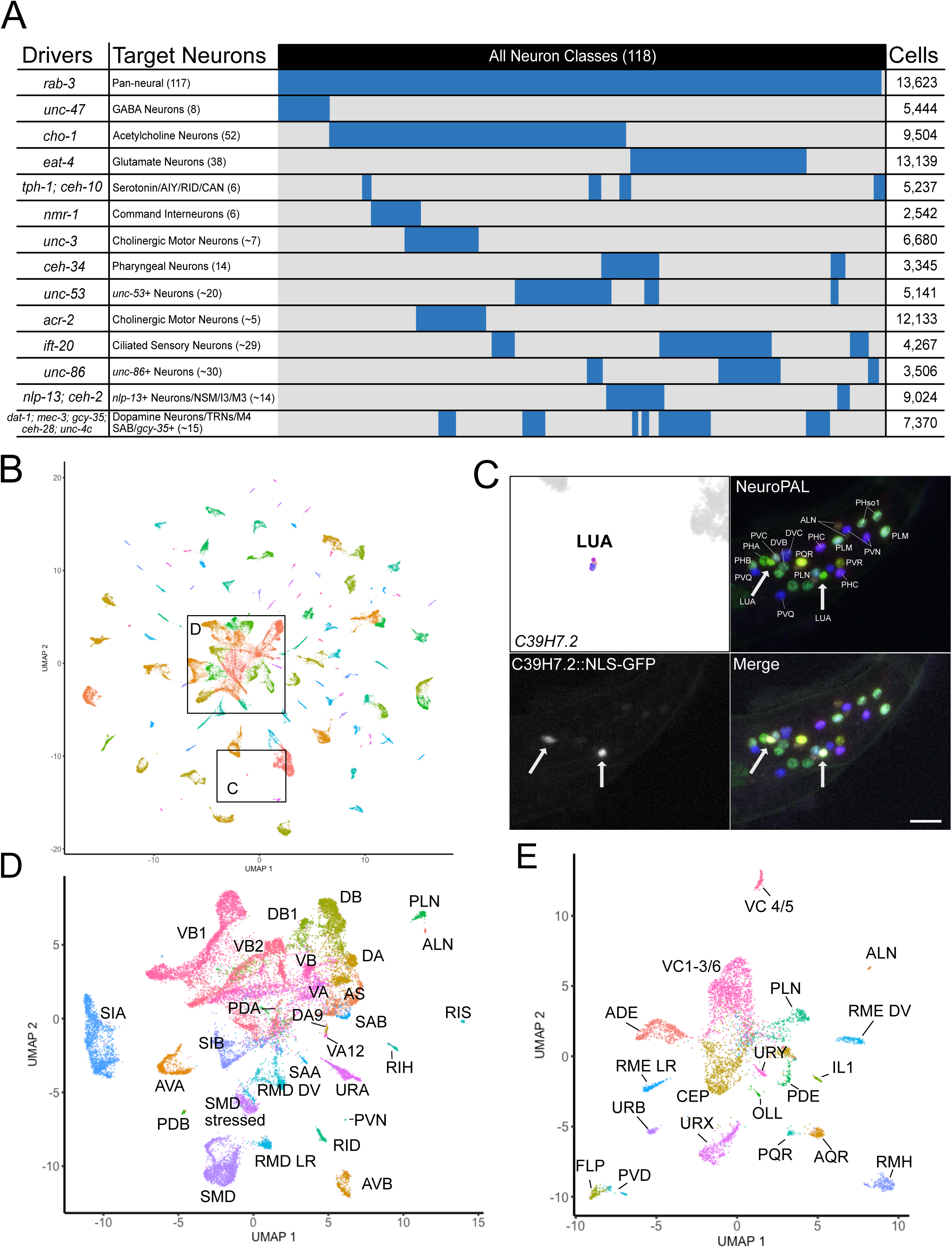
Targeted isolation of selected groups of neurons to capture all known neuron classes for single-cell profiling. A) Graphical representation of neurons targeted in individual experiments. Drivers are reporter genes (e.g., *rab-3, unc-47, cho-1*) used to mark Target Neurons (description + number of neuron types in each group) for each experiment. The column on the right lists the total number of cells obtained per experiment. Some reporter genes were used for multiple experiments. In these cases, the number of cells is the sum across those experiments. B) UMAP projection of all neuronal cells colored by cluster. Boxes outline cells featured in C and D. C) The LUA cluster exclusively expressed *C39H7.2*. Z-projection of confocal stack showing exclusive expression of transcriptional reporter C39H7.2::NLS-GFP in LUA neurons (LUAL and LUAR) (arrows) in tail region of NeuroPAL marker strain. Scale bar = 10 µm. D) Sub-UMAP of central group of cells in B. Clusters are annotated by cell types. E) Sub-UMAP of several neuron types that clustered together in B. This level of resolution clearly separates closely related neuron types (e.g., FLP vs PVD) into individual clusters. See also Figure S1, S2.

We annotated 70,296 cells (69.6% of all single cell profiles) as neurons and 27,427 cells (27.2%) as non-neuronal based on gene expression data from WormBase. We manually excluded non-neuronal cells from the global dataset to create a separate neuronal UMAP (Figure 1B). We detected a median of 1033 UMIs and 363 genes per individual profiled neuron. A substantial fraction of the UMAP clusters could be assigned to individual neuron classes on the basis of known marker genes which cover most of the nervous system, with an average of 40 markers per neuron (Hobert et al., 2016) (Figure 2A-B, Figure S3A-C). For cases in which a cluster could not be readily identified on the basis of existing markers, we generated GFP transcriptional reporters for genes enriched in the target clusters for direct examination *in vivo*. For example, *C39H7.2* was exclusively detected in a small cluster that expressed no known unique markers (Figure 1C). We used the multi-colored NeuroPAL marker strain (Yemini et al., 2019) to determine that a *C39H7.2::NLS-GFP* transcriptional reporter was exclusively expressed in the tail interneuron LUA (Figure 1C). In another example, the nuclear hormone receptor *nhr-236* was detected in three clusters, two of which correspond to the pharyngeal neurons M2 and MC. Based on other genes expressed in the third *nhr-236* cluster (Table S3), we suspected it might correspond to the ring/pharynx interneuron RIP. A *nhr-236::NLS-GFP* transcriptional reporter was consistently expressed in MC, M2 and RIP, confirming our assignment (Figure S3D-E). Overall, we annotated 95.9% of the cells in the entire dataset and identified clusters encompassing all of the 118 anatomically-defined neuron classes in the mature hermaphrodite nervous system (White et al., 1986). Ninety of the 118 neuronal types were detected in distinct, individual clusters in the pan-neuronal UMAP (Figure 2B). The remaining clusters in the pan-neuronal map contained multiple, closely related neuron classes (i.e., dopamine neurons, touch receptor neurons, oxygen-sensing neurons, ventral cord motor neurons). Individual UMAP projections of these clusters facilitated a more precise annotation that revealed an additional 38 identified neuron types. (Figure 1D-E, Figure S4). In one case, the DD and VD classes of ventral cord GABAergic motor neurons were co-mingled in a single cluster that did not separate in the neuronal sub-UMAP, despite known differences in gene expression (Melkman and Sengupta, 2005; Petersen et al., 2011; Shan et al., 2005). Overall, identified neuronal clusters included profiles of a median of 352 cells (Figure 2C).

**Figure 2.**
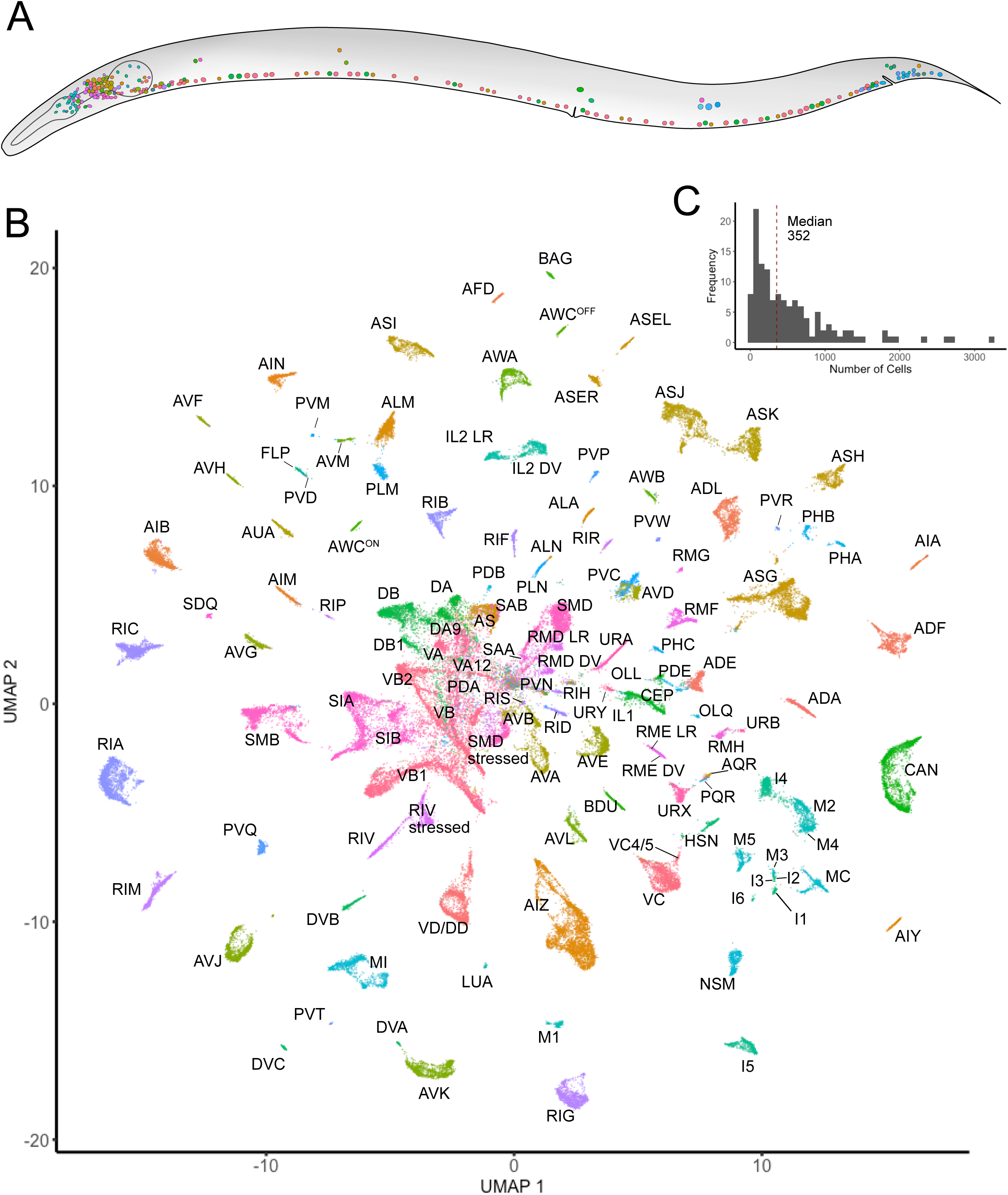
All known neuron types in the *C. elegans* nervous system are identified as individual clusters of scRNA-seq profiles. A) Top – depiction of all neuron types in the mature *C. elegans* hermaphrodite. B) UMAP projection of 70,296 neurons with all neuron types annotated. Sub-types of ten anatomically defined classes were also identified (e.g., ASER vs ASEL, see Figure 3). Colors match neuron depicted in Panel A. C) Histogram showing frequency of the number of single cells per neuron type, with a median of 352. See also Figure S3, S4.

Using 7,390 genes identified as highly variable based on the mean and variance of their expression across neurons, we generated a network that describes the relative molecular relationship of the 128 anatomically distinct neuron classes and subclasses (see below) identified in this work (Figure S4F). This network reveals a clustering of neurons that tracks with functional and anatomical features, including separate groupings of sensory neurons vs motor neurons as well as a distinct cluster of pharyngeal neurons. Interestingly, pre-motor interneurons cluster with motor neurons. Amphid/phasmid sensory neurons are clearly separated from non-amphid/phasmid sensory neuron types. Within amphid/phasmid neurons, some neurons cluster according to sensory modalities. Notably, the chemorepulsive neurons ADL, ASH and PHA/PHB form their own subcluster. Two neuron types, the CO_2_ sensitive BAG neuron and the CAN neuron, show the least similarity to other neuron types. Thus, a systematic comparison of neuron-specific profiles confirms that neurons with shared anatomical and functional characteristics are defined by similar patterns of gene expression.

### Identification of transcriptionally distinct neuronal sub-types

Reporter-based gene expression and connectivity data suggest that some of the 118 anatomically-defined neuron classes may be comprised of separate neuron subclasses (Hobert et al., 2016; White et al., 1986). Our results confirmed this prediction by revealing that 10 of the canonical neuron classes contained transcriptionally distinct subtypes. Consistent with earlier findings (Johnston et al., 2005; Lesch et al., 2009; Pierce-Shimomura et al., 2001; Troemel et al., 1999; Vidal et al., 2018; Yu et al., 1997), we detected individual clusters for the bilaterally asymmetric sensory neuron pairs ASE (ASER and ASEL) and AWC (AWC^ON^ and AWC^OFF^) (Figure 3A, Figure S5A). Previous embryonic and early larval single-cell RNA datasets also detected the ASE and AWC subtypes (Cao et al., 2017; Packer et al., 2019). We used differential expression analysis to identify expanded lists of subtype-specific transcripts for the ASE and AWC subclasses (Figure 3B, Figure S5B). For example, in addition to confirming the left/right asymmetric expression of receptor-type guanylyl cyclase (rGC) genes (Ortiz et al., 2006), we identified several neuropeptide genes specific to either ASER or ASEL (Figure 3A-B, Figure S5A). Other than for the AWC and ASE neuron pairs, we detected no other cases of molecularly separable left/right homologous cells within a neuron class.

**Figure 3.**
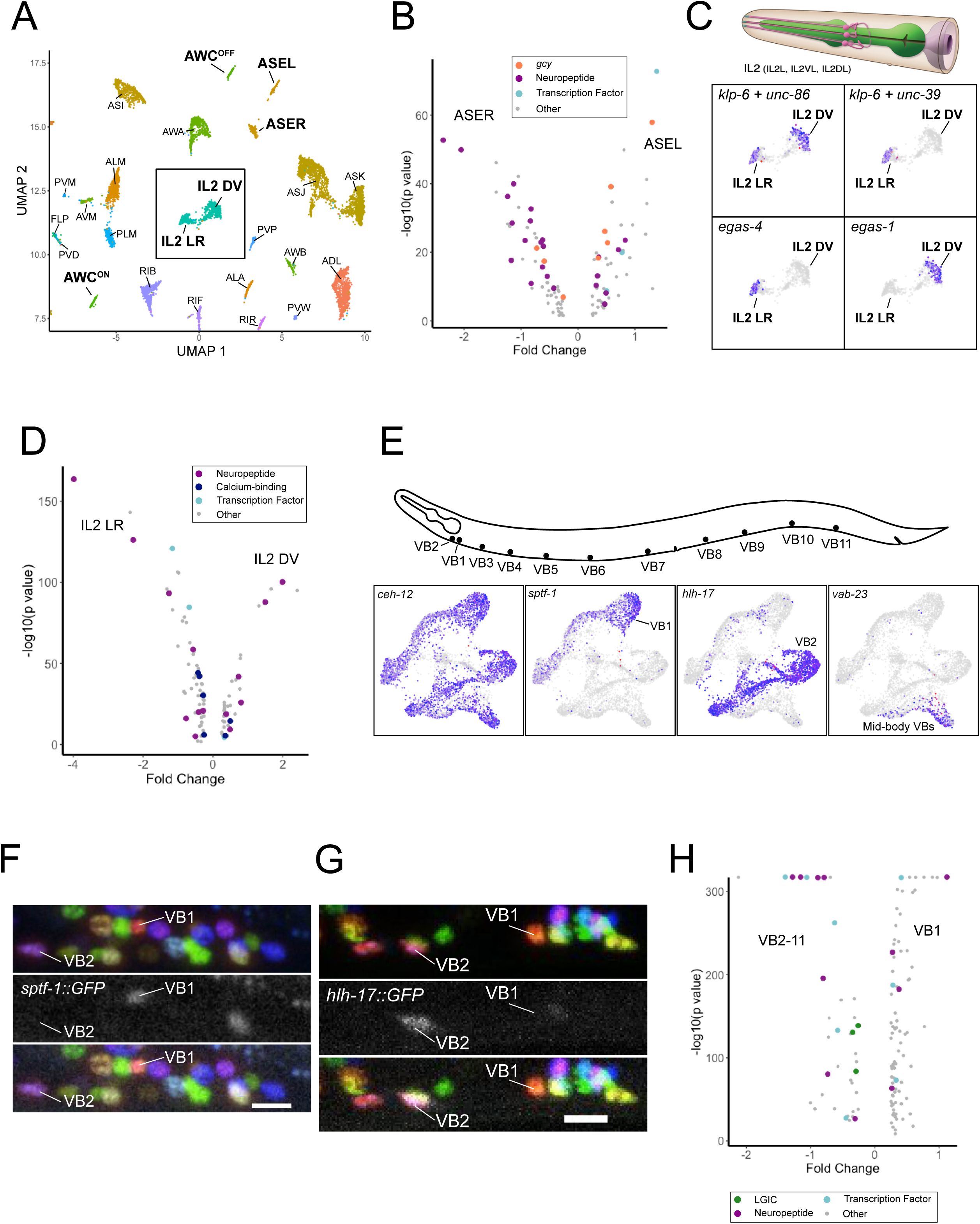
Identification of transcriptionally distinct neuron sub-types. A) Selected region from global neuronal UMAP in Figure 2A highlights examples of neuron classes with distinct sub-types (bold labels). Inset denotes IL2 DV vs IL2 LR sub-types. B) Volcano plot of differentially expressed genes (FDR < 0.05) between ASER and ASEL. Guanylyl cyclases (*gcy* genes), neuropeptides, and transcription factors are marked. C) Top – depiction of 3 pairs of IL2 sensory neurons (IL2L/R, IL2VL/R, IL2DL/R) from WormAtlas. Bottom – Expression of marker genes for all IL2 neurons (*klp-6, unc-86)*, for IL2 LR sub-type (*unc-39*, *egas-4*) and for IL2DV (*egas-1*). D) Volcano plot of differentially expressed genes (FDR < 0.05) between IL2 sub-types. Neuropeptides, calcium-binding genes and transcription factors are marked. E) Top – Locations of VB motor neuron soma in the ventral nerve cord region. Bottom – sub-UMAPs of VB neurons highlighting VB marker (*ceh-12*) and genes (*sptf-1, hlh-17, vab-23*) expressed in specific VB sub-clusters. F) Confocal image of *sptf-1::GFP* reporter in NeuroPAL strain detects *sptf-1* expression in VB1 but not VB2 whereas G) *hlh-17::GFP* is selectively expressed in VB2 but not VB1. Scale bars in F and G = 10 µm. Images are z-projections. H) Volcano plot of differentially expressed genes (neuropeptides, ligand-gated ion channels (LGICs) transcription factors) (FDR < 0.05) between VB1 and all other VB neurons. See also Figure S5.

The remaining eight neuron classes with transcriptionally distinct subtypes are either arranged in radially symmetric groups of 4 or 6 neurons or are distributed along the anterior/posterior axis. We detected distinct subclusters for two neuron classes that display six-fold symmetry at the nerve ring, the inner labial IL2 neurons (Figure 3A, C) and the RMD neurons (Figure 1D, Figure S5). In both cases, a separate cluster for the left/right pair of neurons (e.g., IL2L/R) segregates from a cluster for the dorsal/ventral pairs (IL2DL/R and IL2VL/R). Differentially expressed genes between the IL2 clusters encode neuropeptides, ion channels, calcium binding proteins and transcription factors and point to potentially unique functional roles for the IL2D/V vs IL2L/R subtypes (Figure 3C, D). For the GABAergic RME head motor neurons, we detected distinct dorsal/ventral (RMED/V) and left/right clusters (RMEL/R) (Figure 1E, Figure S5). We also identified multiple clusters for the DA, DB, VA, VB, and VC ventral nerve cord motor neuron classes. In each case, one subtype corresponded to either one or two individual members of these classes. For example, VC4 and VC5, which flank the vulva, clustered independently from the other four VC neurons (VC1-3, VC6) (Figure 1E, Figure S5). For A-class motor neurons (DA, VA), we detected unique clusters corresponding to the most posterior neurons located in the pre- anal ganglion, DA9 and VA12 (Figure 1D, Figure S5). Most of the differences in gene expression among A-class motor neuron subtypes (i.e., DA, VA) are found in genes expressed exclusively in the most posterior neurons. For example, > 90% of the differentially expressed genes between the VA and VA12 subtypes were enriched in VA12 (Figure S5), a finding suggestive of unique functional roles for the most posterior neurons of these classes that are not shared with other A-class motor neurons.

Both B-class motor neuron classes (DB and VB) contained multiple independent clusters (Figure 3E, Figure S5). In contrast to the A-class motor neurons, in which the posteriorly located DA9 and VA12 neurons clustered independently, the most anterior B-class motor neurons (DB1, VB1, VB2) segregated into separate clusters (Figure 3D-F). In these cases, we relied on GFP reporters for cluster-specific genes to identify subtypes because known markers were not available. With 6,576 cells, VB motor neurons were the most abundantly represented neuron class in our scRNA-Seq data set. The homeodomain transcription factor CEH-12 is selectively expressed in VBs (Von Stetina et al., 2007) and marks the VB clusters (Figure 3E). We identified VB1 based on expression of a GFP reporter gene for the subcluster-specific marker *sptf-1* (Figure 3E, F). The VB2 subcluster was similarly identified by the selective expression of *hlh-17::GFP* in VB2 *in vivo* (Figure 3E, G). Interestingly, all of the molecularly distinct subclasses we detected also have known differences in synaptic connectivity (Hobert et al., 2016; White et al., 1986).

We did not detect subtypes for additional classes with 3, 4, or 6-fold symmetry (e.g., SAB, CEP, OLQ, SAA, SIA, SIB, SMD, SMB, URA, URY, and IL1). In some cases, this finding may be due to the low number of cells (< 100 for OLQ, SAA, URY, IL1, see Table S4) assigned to these classes. Alternatively, potential molecular differences among subsets of these neuron types (Hobert et al., 2016) may be limited to a small number of genes that would be insufficient to drive separation in our analyses. Overall, we detected 128 transcriptionally-distinct neuron types.

### Defining gene expression across neuron types

A key consideration for scRNA-Seq data is relating read counts to true gene expression for an accurate determination of whether the detected signal (UMI) for a given gene is indicative of actual expression in a cell type (rather than noise). We reasoned that we could address this question quantitatively by using known high-confidence gene expression results from fluorescent reporter strains to set empirical thresholds for our scRNA-Seq data. Ideally, these reporters should meet two criteria: (1) They must accurately reflect endogenous gene expression, and (2) Expression data must be available for every individual neuron type. For our approach, we selected a ground truth data set of 160 genes with expression patterns for fosmid reporters or reporter-tagged endogenous loci (e.g., CRISPR-engineered genomic markers) that have been mapped with single cell resolution throughout the entire nervous system (Table S5) (Bhattacharya et al., 2019; Reilly et al., 2020; Stefanakis et al., 2015; Yemini et al., 2019).

Next, we aggregated expression of each gene in our scRNA-Seq data set across the single cells corresponding to each neuron type to produce a single expression value for each gene in each neuron type. We used that dataset to select dynamic thresholds for each gene that best reflect the known expression pattern of the ground truth reporters (see Methods). We selected 4 threshold levels (designated as 1-4) offering different compromises between the risk of false positives and false negatives. Threshold 1 captures most gene expression at the price of an increased risk of false positives. On the other hand, threshold 4 offers the most conservative estimate, displaying few false positives but failing to capture some of the expressed genes (Figure S6). Researchers wishing to use this data may select the thresholding level best suited to their particular application (available at cengen.org).

We used threshold 2 for subsequent analyses for profiling gene expression across all neuron types and across gene families. With this threshold, we estimate a true positive detection rate of 0.81 and a false discovery rate of 0.14 (see Methods). The number of detected genes per neuron type was positively correlated with the number of cells (range = 12 [M4] to 3189 [AIZ], median 352) sequenced per neuron type (Figure S6I, Spearman rank correlation = 0.783, p < 2.2e-16) and with the true positive rate (Figure S6J, Spearman rank correlation = 0.6776, p < 2.2e-16). At threshold 2, the number of genes detected per neuron type ranged from 1371 (ALN) to 7429 (ASJ), with a median of 5652. Neurons with fewer cells and fewer detected genes were concentrated in the anterior and pre-anal ganglia (Figure S6H). Nine neuron classes with the fewest detected genes and lowest true positive rates compared to ground truth are labeled in Figure S6J. These cell types are likely to have higher rates of false negatives than other cell types.

We examined the distribution of genes encoding ribosomal proteins as a test of whether our thresholding approach would preserve a predicted ubiquitous pattern of gene expression. Our results show that 65 of the 78 ribosomal genes (83%) are detected in ≥ 98% of neuron types with 53 (68%) expressed in all but one cell type (ALN, the cell type with the fewest genes, Figure 4A). Overall, these results indicate that applying this thresholding approach to our dataset reliably results in accurate expression calling across genes for most cell types in the *C. elegans* nervous system.

**Figure 4.**
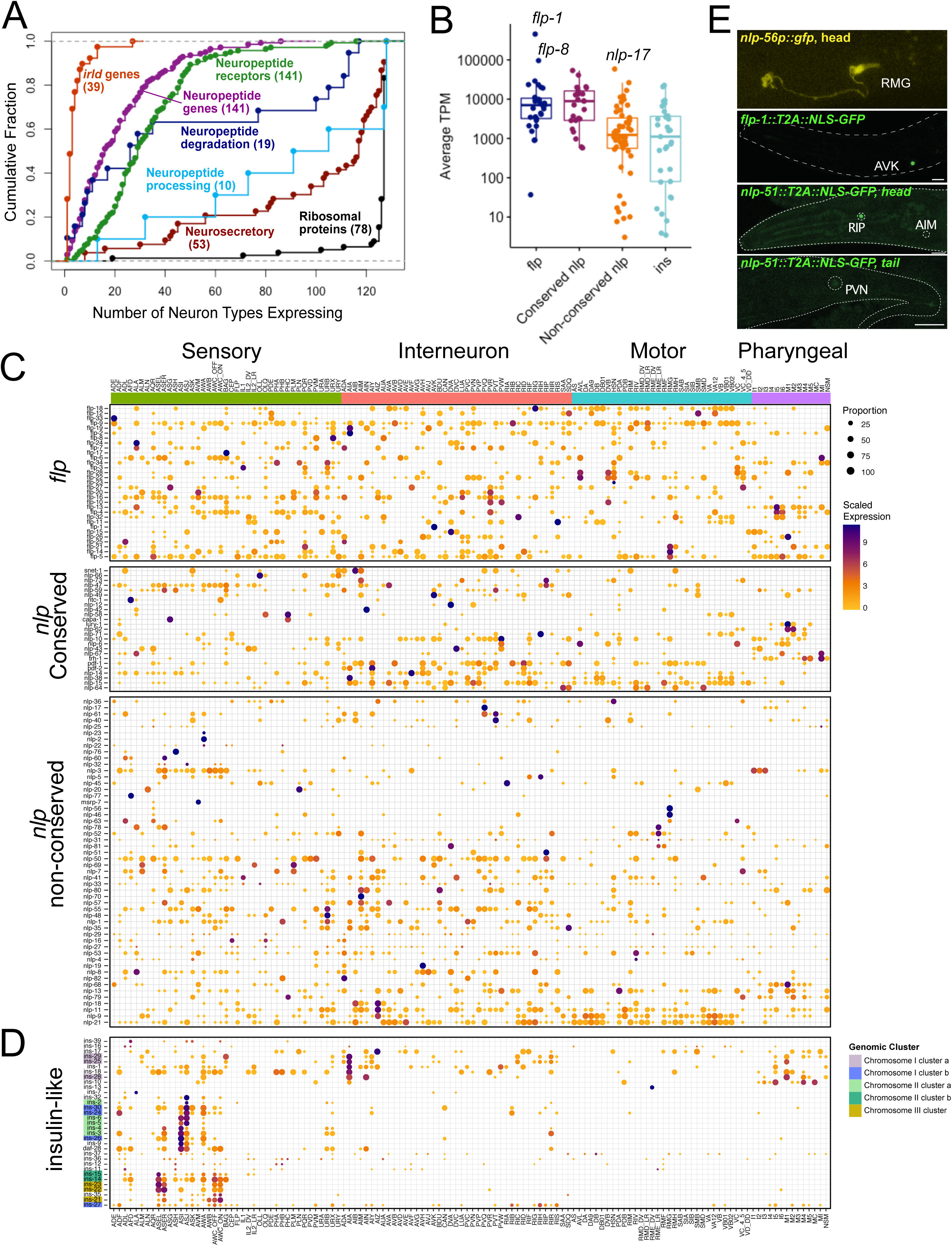
Expression of neuropeptide signaling genes. A) Cumulative distribution plot of the number of neuron types expressing different classes of neuropeptide signaling genes. Each dot is a gene. Numbers in parentheses denote the sum of genes in each category. B) Average expression (TPM) for neuropeptide subfamilies across neuron types. *flp-1*, *flp-8* and *nlp-17* denote highly expressed neuropeptide genes. Boxplot represents 25^th^ percentile, median and 75^th^ percentile. C) Heatmap showing expression of neuropeptide genes (rows) for *flp* (FMRFamide-related peptides) and *nlp* (neuropeptide-like proteins) subfamilies across 128 neuron types (columns) grouped by functional/anatomical modalities (i.e., Sensory, Interneuron, Motor, Pharyngeal). Conserved *nlp* genes are shown separately. Rows are clustered within each family. Circle diameter is correlated with the proportion of cells in each neuron-specific cluster that expresses a given gene. D) Heatmap showing expression of 33 insulin-like peptide (*ins*) genes. Neurons (columns) are grouped by modality and genomic clusters of *ins* genes denoted by colors. E) GFP reporters confirming selective expression of *nlp-56* (promoter fusion) in RMG, *flp-1* (CRISPR reporter) in AVK, and *nlp-51* (CRISPR reporter) in RIP, with weaker expression in PVN and AIM. See also Figure S8, S9.

### Neuron-specific codes of neuropeptide signaling genes

We used the thresholded dataset (threshold 2) to probe expression of selected gene families with predicted functions in the nervous system (Hobert, 2013). Genes families involved in specifying the usage of classic neurotransmitters (acetylcholine [ACh], glutamate [Glu], GABA, monoamines) have been previously mapped throughout the nervous system (Chase and Koelle, 2007; Gendrel et al., 2016; Pereira et al., 2015; Serrano-Saiz et al., 2013) and those assignments were largely confirmed by our scRNA dataset (Figure S7). Next, we examined gene sets encoding proteins involved in neuropeptide signaling, including neuropeptides, receptors, and neuropeptide processing and degradation enzymes (Table S7, Figure 4A). Neuropeptide-encoding genes (31 FMRFamide-like peptides [*flp*], 33 insulin-related peptides [*ins*] and 77 neuropeptide-like proteins [*nlp*] genes, total of 141 genes) were highly expressed and were detected in every neuron class (a minimum of 6, maximum of 62 neuropeptide-encoding genes per neuron) (Figure 4A-D, Figure S9A). Each neuron class expressed on average 23 neuropeptide-encoding genes and each neuron expresses a unique combinatorial code of neuropeptide genes. Sensory neurons and interneurons expressed more neuropeptide genes than motor neurons (Figure S9A, B). Strikingly, neuropeptide encoding genes are among the most highly expressed transcripts in our data set, an observation that parallels similar findings in *Hydra*, *Drosophila* and mouse neurons (Siebert et al., 2019; Allen et al., 2020; Smith et al., 2019). Moreover, the subset of *nlp* genes with homologs in other species (25 of 77, (Husson et al., 2009; Koziol et al., 2016; Mirabeau and Joly, 2013)), along with the *flp* family genes, were detected at much higher levels than *ins* and non-conserved *nlp* genes (Figure 4B).

Whereas several neuropeptide-encoding genes were widely expressed in many neuron types, including *flp-9, flp-5,* and *nlp-21,* we also detected neuropeptide-encoding genes with expression restricted to just one or two neuron types, including exclusive expression of *flp-1* in AVK, *flp-23* in HSN, *nlp-56* in RMG, *nlp-*2 and *nlp-23* in AWA and *ins-13* in RMED/V (Figure 4B). We validated the restricted expression of *nlp-56* in the RMG cluster and *flp-1* in AVK with transcriptional reporters that were selectively expressed in RMG and AVK, respectively, *in vivo* (Figure 4E). Transcriptional reporters for several additional neuropeptide-encoding genes were largely consistent with the single-cell data (Figure S8).

Among neuropeptide-encoding classes, the *ins* genes were the most restricted, expressed in a median of 10 neuron types, compared to medians of 22 and 19 for *flp* and *nlp* genes, respectively. The *ins* genes were primarily expressed in amphid sensory and pharyngeal neurons (Figure 4D). Many *ins* genes are organized in genomic arrays of two to seven genes, but none reside within operons (Pierce et al., 2001). Hierarchical clustering revealed that *ins* genes from the same genomic region showed similar patterns of expression (Figure 4D). For instance, *ins-2*, *ins-3, ins-4, ins-5* and *ins-6* are located in a continuous 12.6 Kb region of chromosome II with no intervening genes. All of these *ins* genes are enriched in the sensory neurons ASI and ASJ (Figure 4D) and may regulate dauer entry or exit and/or aversive olfactory learning (Bargmann and Horvitz, 1991; Chen et al., 2013; Cornils et al., 2011; Hu, 2007; Li et al., 2003; Murphy and Hu, 2013). Two additional genomic clusters of *ins* genes (*ins-14, ins-15* and *ins-21, ins-22, ins-23*) are co-expressed in the ASE and AWC chemosensory neuron pairs which could be indicative of roles in modulating downstream responses to stimuli. An additional cluster (*ins-28, ins-25*, *ins-29*) is primarily expressed in head interneurons and pharyngeal neurons.

Of the more than 140 neuropeptide receptor-encoding genes, most also show highly restricted expression patterns, with a few notable exceptions (Figure 4A). The predicted neuropeptide receptor encoding genes *pdfr-1, npr-23* and *F59D12.1* were widely expressed, in over 100 neuron types. DAF-2, the only insulin/IGF receptor-like tyrosine kinase in *C. elegans*, was also detected in a majority of neurons (103 of 128 neuron types). Most other neuropeptide receptor genes, however, are expressed in a restricted subset of neurons; half of the detected predicted neuropeptide receptors are expressed in 29 or fewer cell types (Figure 4A). Each individual neuron class expressed on average 36 neuropeptide receptor-encoding genes, and each neuron type expressed a unique set of neuropeptide receptors. Sensory neurons and interneurons expressed more neuropeptide receptor genes than pharyngeal neurons, but not more than motor neurons (Figure S9B). With on-going efforts to match neuropeptide GPCRs to their cognate ligands (https://worm.peptide-gpcr.org/project/), these expression data covering all neuropeptide genes and predicted receptors provide a basis for establishing a nervous-system wide map of modulatory neuropeptide signaling.

Consistent with the detection of at least six neuropeptide-encoding transcripts in every neuron type (Figure 4B), neuropeptide processing genes were broadly expressed throughout the nervous system (Figure 4A). Neuropeptide signaling is terminated via degradation of neuropeptides by specific proteases, including neprilysins and dipeptidyl or tripeptidyl peptidases (Hobert, 2013; Turner et al., 2001). These enzymes displayed a bimodal distribution in our data set, with neprilysins showing restricted expression (e.g., 10 of 11 are expressed in < 30 neuron types) and the dipeptidyl and tripeptidyl peptidases largely expressed in > 100 neuron types (Figure 4A). Although neprilysins exhibit broad substrate recognition (Turner et al., 2001), their restricted patterns of expression in specific neurons point to local roles in specific behaviors. For example, the neuropeptide FLP-1, released from the interneuron AVK, acts on SMB and VC motor neurons to regulate movement (Oranth et al., 2018). SMB shows strong expression of *nep-21* and *nep-16* whereas VC motor neurons express dipeptidyl peptidases thus suggesting that different methods of degradation could attenuate signaling by a single neuropeptide (FLP-1) at different post-synaptic targets (SMB vs VC motor neurons). No significant differences were detected across neuron modalities in the numbers of peptide degradation genes (Figure S9B).

We also analyzed the expression of the unusual *irld* and *hpa* gene families, which encode insulin/EGF-receptor-like proteins containing the characteristic extracellular cysteine-rich L domain but lacking tyrosine kinase domains (Dlakic, 2002). *hpa-1* and *hpa-2* modulate lifespan and age-related locomotor decline through negative regulation of EGF signaling (Iwasa et al., 2010). In striking contrast to insulin/IGF receptor-like tyrosine kinase *daf-2,* which is broadly expressed in the nervous system, the 38 *irld* genes and the one *hpa* gene detected (Table S7) in the neuronal dataset showed highly restricted patterns of expression (Figure 4A). 21 of the 39 genes in the *irld* and *hpa* families were detected in only one or two neuron classes and were largely restricted to sensory neurons (Figure S9B); the sensory neuron ASJ expressed 27 of the 38 *irld* genes, and ten *irld* genes were only expressed in ASJ. The expression of several *ins* and *irld* genes in ASJ suggests the possibility of a unique role for ASJ in the regulation of *ins* signaling.

### Neurosecretory machinery

Genes related to vesicle release (SNARE proteins and others, see Table S7) are broadly expressed throughout the nervous system, as expected for a gene family that mediates a fundamental property of neurons. However, the *C. elegans* genome has undergone expansion of genes predicted to be involved in neurosecretion and some of these genes are exclusively expressed in selected neuron types. For example, the *C. elegans* genome encodes 7 synaptotagmins (calcium sensor for vesicle fusion), 9 synaptobrevin-related R-SNAREs and 21 syntaxin-related Q-SNARES (Figure 4A, Figure S9C) (Hobert, 2013). Whereas most of these genes are widely expressed, the R-SNARE encoding transcript, *snb-2*, is only detected in 8 neuron classes (Figure S9C). Similarly, members of the synaptotagmin family, *snt-1*, *snt-3* and *snt-4*, are broadly expressed whereas *snt-2*, *snt-5*, *snt-6*, *snt-7* are detected in a subset of neuron classes. Selective expression of synaptotagmins in specific neurons could be functionally important for fine-tuning calcium dependent neurotransmitter release (Chen and Jonas, 2017). A related question is whether co-expressed members of neurosecretory gene families are differentially distributed to functionally distinct presynaptic domains of a common neuron. Taken together, these patterns of expression suggest that neurosecretory mechanisms may be subjected to cell-type specific modulation.

### Differential expression of ion channels

As in other organisms, *C. elegans* has 2 types of ionotropic receptors: (1) pentameric Cys-loop ligand-gated ion channels (LGICs) and (2) tetrameric ionotropic glutamate receptors (Hobert 2013). Ionotropic receptor genes show specific expression profiles throughout the nervous system (Figure S10). Each neuron expresses on average 20 ionotropic receptors, and each individual neuron type expresses a unique combination of these genes. We detected expression of a large subset of LGICs in cholinergic ventral nerve cord motor neurons (VNC MNs) with the exception of the VC class (Figure S10A). Interestingly, these motor neuron-expressed LGICs include subunits of both acetylcholine-gated cation (e.g., *acr-2, acr-12, unc-63*) and chloride channels (e.g., *lgc-46, acc-1*), as well as GABA-gated cation (*exp-1, lgc-35*) and chloride channels (e.g., *unc-49, lgc-37, gab-1*) (Jones and Sattelle, 2008; Jones et al., 2007), indicating that ACh and GABA may be capable of eliciting both excitatory and inhibitory responses in the same postsynaptic cells (Figure S10A) (Fox et al., 2005; Gendrel et al., 2016; Pereira et al., 2015). VNC MNs also expressed several orphan LGICs with unknown ligand specificity and predicted anion selectivity (“Diverse” LGICs in Figure 4D, e.g., *lgc-41, lgc-42, ggr-1*). Together, motor neurons expressed more ionotropic receptors than any other neuron modality (Figure S9D).

Ionotropic glutamate-gated cation channels were mostly expressed in interneurons (particularly the command interneurons AVA, AVD, AVE, and PVC) and in motor neurons (particularly head motor neurons), consistent with previous reports (Brockie et al., 2001). Three unnamed genes homologous to *glr* genes are highly enriched in subsets of pharyngeal neurons. T25E4.2, ZK867.2 and C08B6.5 overlap closely with *glr-8* expression in I6 and NSM, and W02A2.5 is co-expressed with *glr-7* in I3. These unnamed genes, along with *glr-7* and *glr-8*, may belong to a subtype of ionotropic receptors that serve as chemosensory receptors in flies (Croset et al., 2010; Hobert, 2013).

Additionally, several uncharacterized orphan LGIC genes are expressed selectively in pharyngeal neurons (Figure S10B). For example, *lgc-5, lgc-6, lgc-7* and *lgc-8* are strongly expressed in the I1 pharyngeal neuron, whereas a different group of LGICs (*lgc-1, lgc-17, lgc-18, lgc-19* and *lgc-29)* is detected in I2 and M3 (Figure S10B). Many of these genes reside close together in the genome (but are not within operons) and belong to an orphan sub-family of Cys-loop gated channels (Jones and Sattelle, 2008; Jones et al., 2007). Members of the *C. elegans* Cys-loop LGIC superfamily have been shown to contribute to sensory function (Cohen et al., 2014), including the detection of protons (Beg et al., 2008). The pharyngeal neurons I1 and I2 modulate pharyngeal function in response to light and hydrogen peroxide (Bhatla and Horvitz, 2015; Bhatla et al., 2015), raising the possibility that these uncharacterized LGICs may contribute to pharyngeal neuron-to-neuron communication or to the response to environmental stimuli.

### Matching ionotropic receptor expression to synaptic connectivity

A previous analysis detected expression of ionotropic GABA receptors in many neurons that are not innervated by GABAergic neurons, suggesting pervasive extra-synaptic, volume transmission (Gendrel et al., 2016). We used the expression patterns of ionotropic Glu and predicted ACh receptors to ask whether volume transmission is also likely for these neurotransmitter systems. We find that predicted ionotropic ACh receptors (52 nAChR-like genes and 7 *acc*-like genes) are detected in all neurons, including those that are not postsynaptic to ACh-expressing neurons. Ionotropic Glu receptors (16 genes, including *nmr*, *glr*, *avr* and *glc* genes) are detected in all 33 neuron types that are not postsynaptic to Glu-expressing neurons (Figure S10C), thus suggesting wide-spread extra-synaptic signaling for ACh and Glu as well as for GABA.

Voltage-gated channels make up another major family of ion channels, of which there are two types, voltage-gated calcium channels (VGCC) and voltage-gated potassium channels. In contrast to ionotropic receptors, voltage-gated calcium channels are broadly expressed across the nervous system (Figure S10C). With the exception of *ccb-2*, which was selectively detected in SIB head neurons and in AS, DA, and VA VNC motor neurons (Lainé et al., 2011), all VGCC genes were expressed in > 50 neuron types (Figure S10C). Potassium channels show diverse expression throughout the nervous system (Figure S10C). For example, the TWK-type 2-pore channels, of which the *C. elegans* genome encodes an unusually large number (47), are selectively expressed in specific neurons across the nervous system (Figure S9C). The number of TWK genes per neuron ranged from 1 to 18; all neuron types expressed at least one TWK gene. Notably, VNC motor neurons expressed a large subset of TWK genes. Overall, motor neurons express more TWK channels than sensory, interneuronal or pharyngeal neurons (Figure S9C, D).

Members of the DEG/ENaC family of cation channels also display a restricted pattern of expression (Figure S10C). Lastly, members of a large family of ion channels of unexplored function, the bestrophin-type chloride channels, of which there are 26 in the *C. elegans* genome (Hobert, 2013), also show selective expression throughout the nervous systems with the exception of *best-18* and *best-19* which are detected in 49 and 67 neurons, respectively (Figure S9C).

### Sensory receptors and downstream signaling components

Our scRNA-Seq data confirm the reporter gene-based predictions of sensory/rhodopsin-type chemoreceptor GPCRs (G-protein-Coupled Receptors) in the nervous system (Troemel et al., 1995; Vidal et al., 2018) and also identify many additional neurons that express members of the GPCR family. Whereas GPCR expression is clearly enriched in sensory neurons, all non-sensory neurons also express at least one sensory-type GPCR gene (Figure S11A, C). Within the sensory system, GPCR expression is biased for subsets of ciliated amphid and phasmid neurons. For example, >300 GPCRs are expressed in ADL and >200 GPCRs each in ASH, ASJ and ASK (Figure S11A). Of the 780 chemoreceptor GPCRs detected in the neuronal dataset (Table S7), 137 (17.5%) were detected in single neuron classes, and 723 (92.6%) were detected in fewer than 10 cell types (Figure S11D).

Signaling components downstream of GPCR genes include heterotrimeric G-proteins, arrestin-like (arrd) genes that modulate GPCR activity and RGS proteins, which regulate G-alpha protein function. Each family of these regulatory factors is expanded in the *C. elegans* genome (21 G-alpha genes, 31 arrd genes, 21 rgs genes) (Hobert, 2013). The canonical G-alpha genes, *gsa-1* (G-alpha S), *goa-1* (G-alpha O) and *egl-30* (G alpha Q) are broadly expressed throughout the nervous system (Figure S9B) (Korswagen et al., 1997; Mendel et al., 1995; Moghal et al., 2003). Most other gene family members show sporadic patterns of expression in selected neurons with the exception of a subset of sensory neurons in which most G-protein encoding genes are detected. As noted above, this group of ciliated sensory neurons (ADF, ADL, ASH, ASI, ASJ, ASI, PHA, PHB) also express high numbers of GPCRs (Figure S11B). Most *rgs* genes were broadly expressed with the exception of *rgs-3*, which was limited to sensory neurons (Figure S11B, D). Several arrestin genes showed selective expression, including *arrd-23*, which was expressed almost exclusively in dopaminergic neurons (ADE, CEP, PDE), and *arrd-28*, which was only detected in the IL2 inner labial neurons and the O2-sensing neurons AQR, PQR, and URX (Figure S11B, D). Many other arrestin genes were broadly expressed.

### Differential expression of gene regulatory factors

We also interrogated gene families involved in gene regulation, including all predicted transcription factors (TFs) [wTF 3.0, (Fuxman Bass et al., 2016)] and RNA-binding proteins (Tamburino et al., 2013) (Figure 5A, B, Figure S12). C2H2 zinc finger proteins can be involved in both DNA and RNA binding (Tamburino et al., 2013); we treated this class as TFs and removed them from the list of RNA-binding proteins. We also removed ribosomal proteins from the RNA-binding protein list. 705 of 941 (75%) of predicted transcription factors and 497 of 587 (86%) of predicted RNA-binding proteins were detected in at least one neuron type. Overall, transcription factors were more restricted in their expression than RNA-binding proteins (Figure 5B).

**Figure 5.**
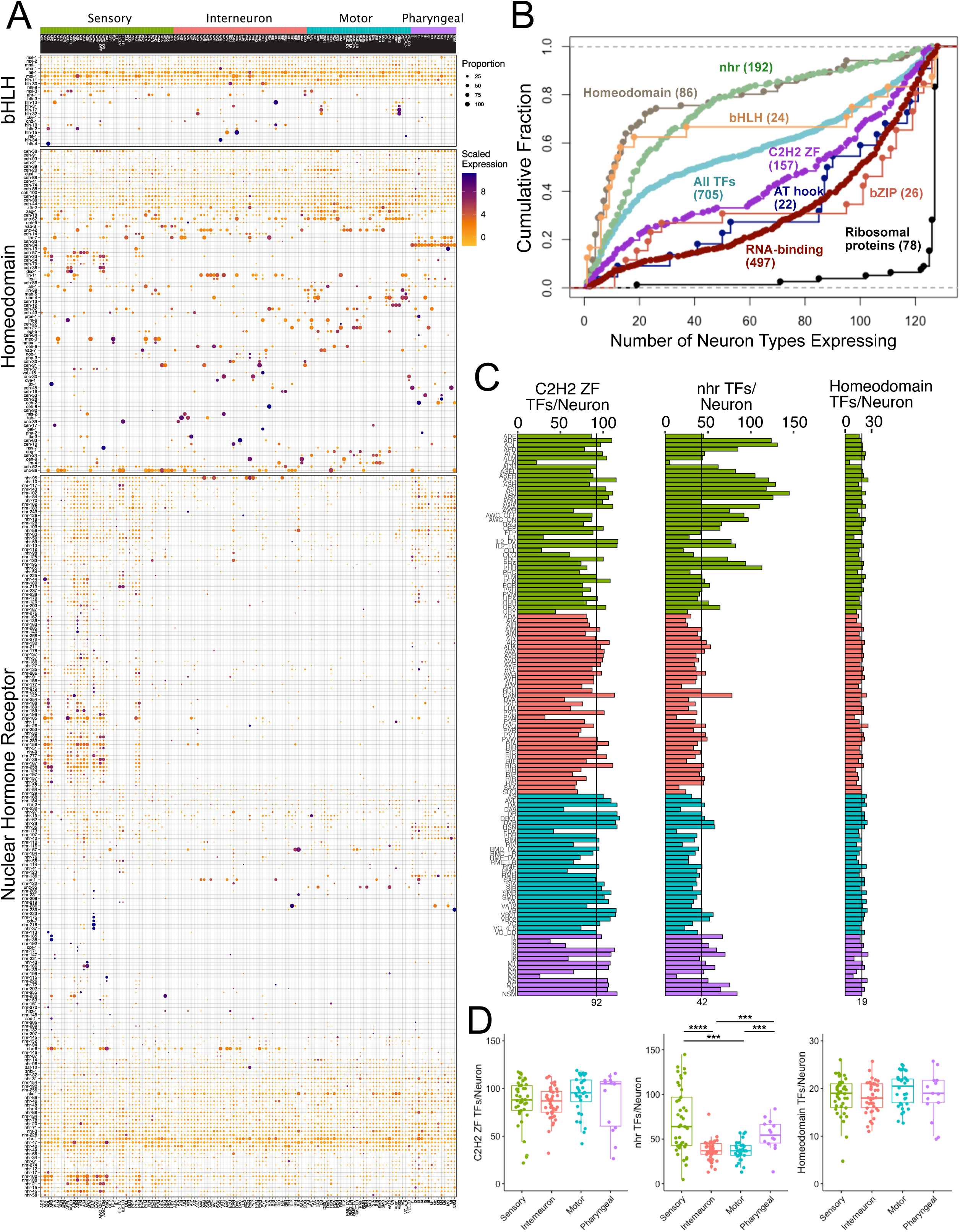
Expression of transcription factor families. A) Heatmap showing expression of bHLH (basic helix-loop-helix), Homeodomain and Nuclear Hormone Receptor (NHR) transcription factors (TFs) across 128 neuron types (columns) grouped by neuron modality. TFs are clustered for each subfamily. Circle diameter is correlated with the proportion of cells in each neuron-specific cluster that expresses a given gene. B) Cumulative distribution plot of number of neuron types expressing genes encoding Homeodomain, bHLH, NHR, C2H2 ZF (Zinc Finger), AT hook, bZIP transcription factor (TF) families, RNA binding proteins and ribosomal proteins (see also Figure 4A). C) Bar graphs showing the number of C2H2, NHR and Homeodomain TFs in each neuron type, grouped by modality (Interneuron, Sensory neuron, Motor neuron, Pharyngeal neuron). D) Quantitative comparison of TFs per neuron for C2H2 (left), NHR (middle) and Homeodomain TFs (right) shows enrichment in sensory neurons for NHR TFs, but no differences across neuronal functional groups for C2H2 or Homeodomain TFs. Boxplots show median and interquartile range (25^th^ – 75^th^ percentiles), statistical tests: Kruskal-Wallis. ***p < 0.001, ****p < 0.0001. See also Figure S12.

We analyzed expression of all TF classes that contain more than 15 members (homeodomain, nuclear hormone receptor [*nhr*], helix-loop-helix [bHLH], C2H2 zinc finger, bZIP, AT hook and T-box genes) and found distinct themes for individual gene families. On one end of the spectrum are T-box genes, very few (2) of which are expressed in the postembryonic nervous system (Figure S12). On the other end of the spectrum are AT hook and bZIP gene family members, where individual members are expressed broadly throughout the nervous system. Members of the bHLH and large family of C2H2 TFs show a combination of genes with broad vs selective expression in the nervous system (Figure 5A, B, S12). Homeodomain and *nhr* TF encoding genes show the most restricted patterns of expression but there are obvious differences between these two families. Each neuron expressed multiple different *nhr* TFs, but sensory and pharyngeal neurons expressed many more *nhr* TFs than either motor neurons or interneurons (Figure 5A-D). Ciliated sensory neurons expressed the highest numbers of *nhr* genes. Thirteen neuron types (ADF, ADL, ASER, ASG, ASH, ASI, ASJ, ASK, AWA, AWC_ON, AWC_OFF, PHA, PHB) expressed more than 90 different *nhr* TFs. Notably, ASJ expressed 144 *nhr* TFs, 75% of the 191 *nhr* TFs detected in the entire neuronal dataset (Figure 5A, C). Abundant expression of a broad array of *nhr* genes in sensory neurons is suggestive of specific roles in mediating transcriptional responses to sensory stimuli. In agreement with a recent report (Reilly et al., 2020), most homeodomain TFs are sparsely expressed throughout the nervous system (Figure 5A, B) and each neuron type expresses its own unique combination of homeodomain transcription factors. A similar discriminatory power is observed for *nhr* genes, but since *nhr* genes are generally more broadly expressed, each individual combinatorial code is composed of many more genes than for the homeobox gene code (Figure 5A, B).

### Single neuron-expressed genes

Depending on threshold values, between 160 to 1348 genes are exclusively expressed in a single neuron type (Tables S8-S11). At the least stringent threshold (threshold 1), 160 genes selectively label 41 of the 118 neuron classes, whereas 1348 genes at the most stringent threshold (threshold 4) individually mark 105 of the 118 canonical neuron classes as well as specific sub-classes. The single-neuron specificities of many genes in the most stringent threshold dataset (1348 genes) are validated by published reporter gene analysis. For example, fosmid-based reporters for the *ceh-63* (DVA), *ceh-28* (M4) and *ceh-8* (RIA) homeobox genes match the neuron specificity of the scRNA-Seq results (Reilly et al., 2020). The cis-regulatory control regions of these genes are candidate drivers for genetic access individual cells in the nervous system (Lorenzo et al., 2020). For example, they could be used for manipulating the activity of individual neuron classes or for rescue analysis of mutant phenotypes. Neurons that are not covered by single neuron-specific drivers can be genetically accessed by the unique intersection of drivers that are more broadly expressed.

### Bulk RNA-sequencing confirms scRNA-Seq results

To validate our scRNA-Seq dataset with an orthogonal approach, we generated bulk RNA-Seq profiles for eight neuron types: ASG, AVE, AVG, AWA, AWB, PVD, VD, and DD. In our scRNA-Seq experiments, we used FACS to collect samples with multiple neuron types which were then disambiguated during *post hoc* analysis. By contrast, in our bulk RNA-Seq experiments, we used FACS to collect each neuron type separately (Spencer et al., 2014). For each neuron type, we collected 2-5 biological replicates and averaged the results. To compare bulk RNA-Seq results for each neuron type to its corresponding single-cell profile, we selected the top marker genes for the corresponding single cell clusters (average log fold change > 2 and adjusted p-value < 0.001 when compared against all other neurons). Importantly, ‘marker’ in this analysis indicates enrichment rather than exclusive expression within a single cluster. We performed differential gene expression analysis for each bulk RNA-seq profile vs a pan-neuronal bulk RNA-Seq data set. We then checked for enrichment of the single cell cluster markers in the bulk data sets. We found that for every set of scRNA-Seq markers, the bulk neuron type with the most enrichment of these markers was the expected neuron class (Figure 6A). For example, the ASG markers defined by our scRNA-Seq analysis (left column) are enriched ∼24-fold (2^4.61^) in ASG bulk RNA-Seq profile (top left cell). By contrast, markers for other cells defined by scRNA-Seq analysis are depleted in ASG bulk data (remainder of top row; log_2_ change < 0). Thus, the independently-derived single cell and bulk RNA-Seq data sets yielded consistent gene expression profiles.

**Figure 6.**
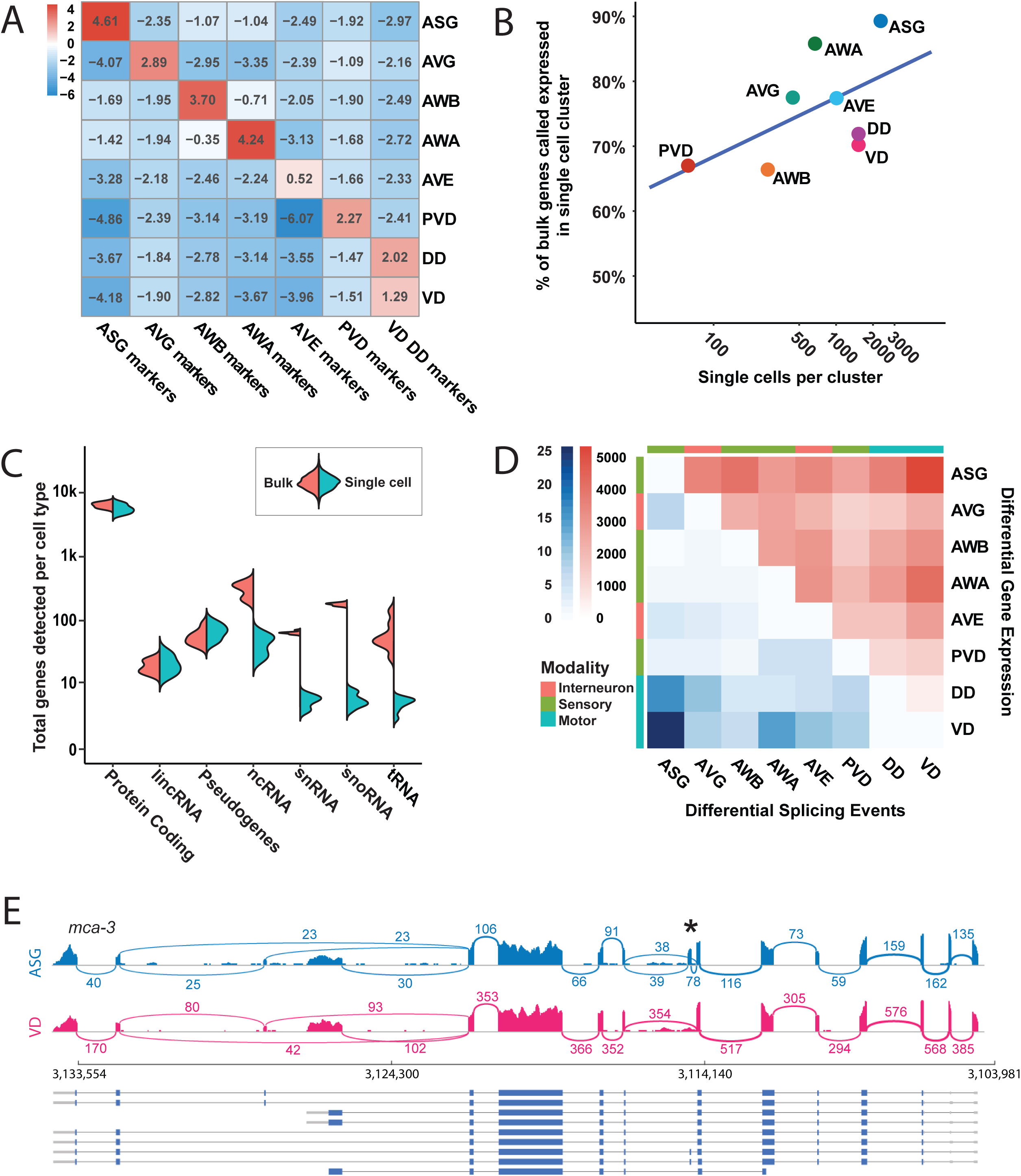
Comparison of bulk and single-cell RNA-Seq. A) For each set of marker genes determined from scRNA-Seq (Methods), we determined its bulk enrichment in each neuron type compared to a pan-neuronal sample set. Heatmap showing the enrichment of single cell marker genes (columns) in bulk RNA-Seq comparisons of one cell type to a pan-neuronal sample set (rows) for ASG, AVE, AVG, AWA, AWB, DD, PVD, and VD neurons. Pairwise Wilcoxon test within each marker gene set (comparing the enrichment in each bulk cell type to all other cell types) showed p-values < 0.001 for all comparisons except for AVE markers (all comparisons p-value > 0.05). B) Linear regression showing the relationship between the single cell cluster coverage of protein coding genes detected by bulk RNAseq datasets, and the number of cells in the cluster. VD and DD bulk datasets both compared to the same VD/DD cluster. R^2^ = 0.34. C) Split violin plot showing the number of genes detected in bulk and single cell datasets for seven classes of RNA. For each RNA class, bulk data is shown as the left violin-half, and single cell data is shown as the right violin-half. Colors reflect the neuron type, as in B. D) Heatmap showing the number of differentially expressed genes (red, upper half), and the number of differential splicing events (blue, lower half), in pairwise comparisons of bulk RNAseq datasets. E) Sashimi plot representing the splicing pattern of the gene *mca-3* in representative samples of neurons ASG and VD. The vertical bars represent exonic read coverage while the arcs indicate the number of junction-spanning reads. The (*) highlights an alternatively spliced exon included in ASG but excluded in VD. See also Figure S13.

The number of cells in each neuron type cluster in our scRNA-Seq dataset varies widely (12 - 3189, Figure 2C). We compared gene detection against our bulk RNA-Seq samples to assess how cluster size relates to sequencing depth at the gene level. For this experiment, we used gene expression values for protein coding genes (threshold 2) in single cell clusters and in bulk data for individual neurons at a comparable threshold (See Methods). We determined that the number of cells in a single cell cluster correlates with how well that cluster detects protein coding genes in the bulk dataset (Figure 6B) (R-squared = 0.387, regression coefficient = 13.256, std err = 6.818). This analysis indicates that even clusters with low numbers of cells (Figure S6J) can include useful information about protein coding gene expression but may also fail to provide a comprehensive profile.

### Bulk RNA-sequencing detects additional classes of non-coding RNAs

We also sought to interrogate the extent to which the scRNA-seq dataset is biased against certain categories of genes. Non-poly adenylated genes, for example, should be under-represented in the single cell dataset. We found that protein coding genes, lincRNAs and pseudogenes (which often have poly(A) tails) show similar coverage in both bulk and scRNA-Seq data sets. However, ncRNAs, snRNAs, and snoRNAs, which generally lack poly(A) tails, are abundantly represented in bulk RNA-Seq data but, as expected, are rarely detected in single cell samples (Figure 6C). The smallest species of ncRNAs, miRNAs and piRNAs, are rarely detected in our bulk profiles because of a size exclusion step in library preparation, and their characterization awaits further studies.

### Bulk RNA-sequence data sets distinguish closely related GABAergic motor neuron types

Although recognized as anatomically distinct neuron classes (White et al., 1986), DD and VD GABA neurons were not distinguishable as separate clusters in our scRNA-Seq data set (Figure 2B). However, previous studies identified a limited subset of markers that can distinguish DD and VD neurons and we used the corresponding reporter genes to isolate each class separately for bulk RNA-Seq (Table S1) (He et al., 2015; Kim and Li, 2004; Mimi Zhou and Walthall, 1998; Petersen et al., 2011; Shan et al., 2005). We compared gene expression in VD and DD bulk samples, and detected 270 differentially expressed genes (p < 0.01). Notably, the neuropeptide gene *flp-13* and the transcription factor *irx-1* are selectively expressed in DD neurons in this comparison (Petersen et al., 2011; Shan et al., 2005). Substantially more genes were identified as differentially expressed in all other pairwise comparisons of our neuron-specific bulk RNA-Seq data sets (Figure 6D). Together, these results suggest that DD and VD GABAergic neurons are more closely related than are other pairs of different neuron types and that methods for distinguishing neuron types from single cell data are relatively insensitive to small differences in gene expression. Thus, we expect that deeper sequencing may reveal additional neuron subtypes that this work has not uncovered.

### Widespread differential splicing between neuron types

Alternative splicing plays a critical role in the development and function of the nervous system (Raj and Blencowe, 2015; Vuong et al., 2016). Differential splicing has been reported between individual neuron types in *C. elegans* (Moresco and Koelle, 2004; Norris et al., 2014; Thompson et al., 2019; Tomioka et al., 2016). However, despite recent progress (Dehghannasiri et al., 2020; Patrick et al., 2020) the strong 3’ bias of the 10x Genomics scRNA-Seq method limits its use for distinguishing alternatively spliced transcripts (Arzalluz-Luqueángeles and Conesa, 2018). We thus took advantage of the bulk RNA-Seq profiles of specific neuron types to look for evidence of differential patterns of alternative splicing between *C. elegans* neurons.

We used the software SplAdder (Kahles et al., 2016) to discover 111 high confidence occurrences of differential use of splicing sites between these 8 neuron classes (Figure 6D, E, Table S12). Most neuron pairs displayed some differential use of splicing sites (Figure 6D), with wide variations between pairs. For example, we detected 16 differential splicing events between ASG and VD, and only 2 differences between ASG and AWA.

In addition, we noticed several instances of previously unannotated exons expressed in a specific neuron type. For example, the *mbk-2* transcript in AWA includes an additional 77 nt sequence corresponding to an alternative 5’ exon that is not expressed in the other seven neuron types in our data set (Figure S13). Inclusion of this exon alters the N-terminal end of the predicted MBK-2 protein, likely encoding a functional isoform with additional putative phosphorylation sites identified by Prosite (Sigrist et al., 2013). This *mbk-2* exon is predicted by GenemarkHMM (Pavy et al., 1999) but its expression was not detected in whole-worm RNA-Seq (Tourasse et al., 2017). By filtering known exons from the splicing graph generated by SplAdder, we recovered 63 putative novel exons (Table S13, see Methods). Thus, our data underscores the capacity of bulk RNA-Seq of single neuron types to detect differential splicing events that could not be reliably detected either by whole animal bulk RNA-Seq or by 10x Genomics scRNA-Seq.

### Consensus gene modules reveal cell type homology across species

We next sought to resolve the cell type evolution between the nervous systems of *C. elegans* and other species at the molecular level. We took advantage of the single-cell transcriptomic data from the entire mouse nervous system to examine the molecular signatures from both *C. elegans* and mammals (Zeisel et al., 2015). Consensus modules, which describe subsets of genes coherently co-expressed in both species (Langfelder and Horvath, 2008), offer an avenue to identify common cell type signatures across species. With this approach, we uncovered a consensus module (C1, Table S14) comprising 83 homologous genes characterized by their high expression in neurons of both species (Figure 7A). Similarly, a second consensus module (C2, Table S15), which contained 40 homologous genes in both species, showed specific expression enrichment in sensory neurons of *C. elegans* and neurons from the mouse peripheral nervous system (PNS), again demonstrating a conservation of a core signature across evolution (Figure 7B).

**Figure 7.**
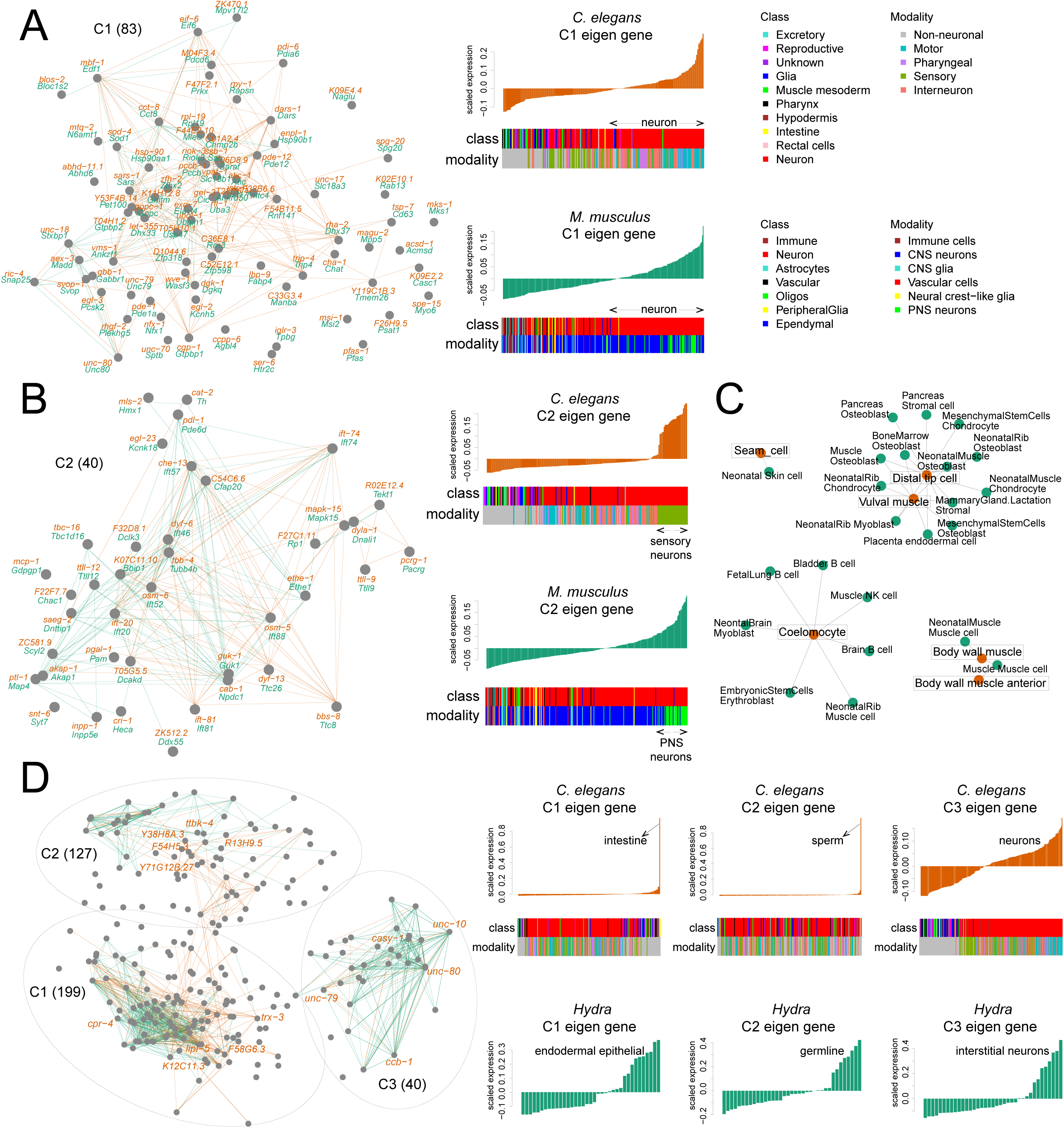
Molecular evolution of cell types in *C. elegans*. A) Left: network showing the expression similarity of 83 homologous genes from the consensus module 1 (C1) detected in *C. elegans* and mouse. Gene names are marked for each node (orange, *C. elegans*; green, mouse). Edges indicate expression profile similarity between pairwise genes measured by Pearson correlation coefficient (≥ 0.7) in each species (orange, *C. elegans*; green, mouse). (Right) Eigen gene patterns of C1 in *C. elegans* (top) and mouse (bottom), color coded by cell type class and modality. B) Network for 40 homologous genes in C2 and eigen gene patterns showing expression in sensory neurons (*C. elegans*) and PNS neurons (Mouse). C) Two-mode network demonstrating the transcriptome similarities between cell types from *C. elegans* (orange) and mouse (green). Cell types with Pearson correlation coefficients < 0.7 are not shown. D) Consensus network comprising three modules co-expressed in both *C. elegans* and *Hydra*, and their eigen gene expression patterns in each species. The top five highly connected genes in each module are labeled.

We also observed a robust correspondence between *C. elegans* non-neuronal cells and certain mouse non-neuronal cell types (Han et al., 2018). This included an expected similarity between muscle cells of both species, and between *C. elegans* coelomocytes and different mouse immune cell types (Figure 7C). Moreover, we discovered a strong transcriptomic match between nematode seam cells and mouse neonatal skin cells, as well as between *C. elegans* distal tip cells and vulval muscle cells and mouse stromal cells, osteoblasts and chondrocytes (Figure 7C).

We further probed the molecular evolution between *C. elegans* and the cnidarian polyp *Hydra* (Siebert et al., 2019). A consensus gene network constructed across the two species revealed three modules, two of which were centered on non-neuronal cell types and one neuronal module (Figure 7D). Module C1 (199 homologous genes, Table S16) was characterized by selective expression in *C. elegans* intestine and in *Hydra* endodermal epithelial cells. In module C2 (127 homologous genes, Table S17), we detected a prominent signature for *C. elegans* sperm cells and *Hydra* germline cells. Lastly, in module C3 (Table S18), 40 homologous genes were overrepresented in neurons and interstitial neurons of *C. elegans* and *Hydra*, respectively. The findings of conserved molecular signature provide a window into the evolution of neuronal cell types.

### Analysis of cis-regulatory elements reveals a rich array of 5’ and 3’ motifs

Transcription factors regulate gene expression at cis-regulatory elements in promoter and enhancer regions. Similarly, miRNAs and RNA binding proteins interact with 3’ UTR sequences to modulate gene expression. To identify candidate cis-regulatory elements that underlie the unique patterns of gene expression among neuron types, we used the FIRE motif discovery algorithm. FIRE detects DNA motifs within promoter sequences and linear RNA motifs in 3’ untranslated regions (UTRs) among cohorts of similarly regulated genes (See Methods) (Elemento et al., 2007).

We calculated z-scores of log-transformed expression values to normalize gene expression relative to the mean and variance across all neurons. For each neuron, we grouped genes into bins ranging from high (far right column) to low relative gene expression (far left column) and conducted de novo motif discovery (Figure 8A). FIRE detects motifs that are significantly informative of relative gene expression in each neuron type, revealing a gradient of enrichment and depletion across genes with similar expression. Motifs of positive regulators, for example, should be significantly over-represented (yellow squares, red borders) in genes with high z-scores (right columns). A subset of the 5’ DNA motifs matched transcription factor DNA binding preferences from the CIS-BP and JASPAR databases (Khan et al., 2018; Weirauch et al., 2014). For example, a motif corresponding to the DNA binding sequence (CTACA) of several *nhr* transcription factors, including ODR-7, is over-represented in genes that are highly enriched in the AWA neuron class (Figure 8 A). Notably, ODR-7 is exclusively expressed in AWA where it regulates neuron identity (Colosimo et al., 2003; Sengupta et al., 1994, 1996).

**Figure 8.**
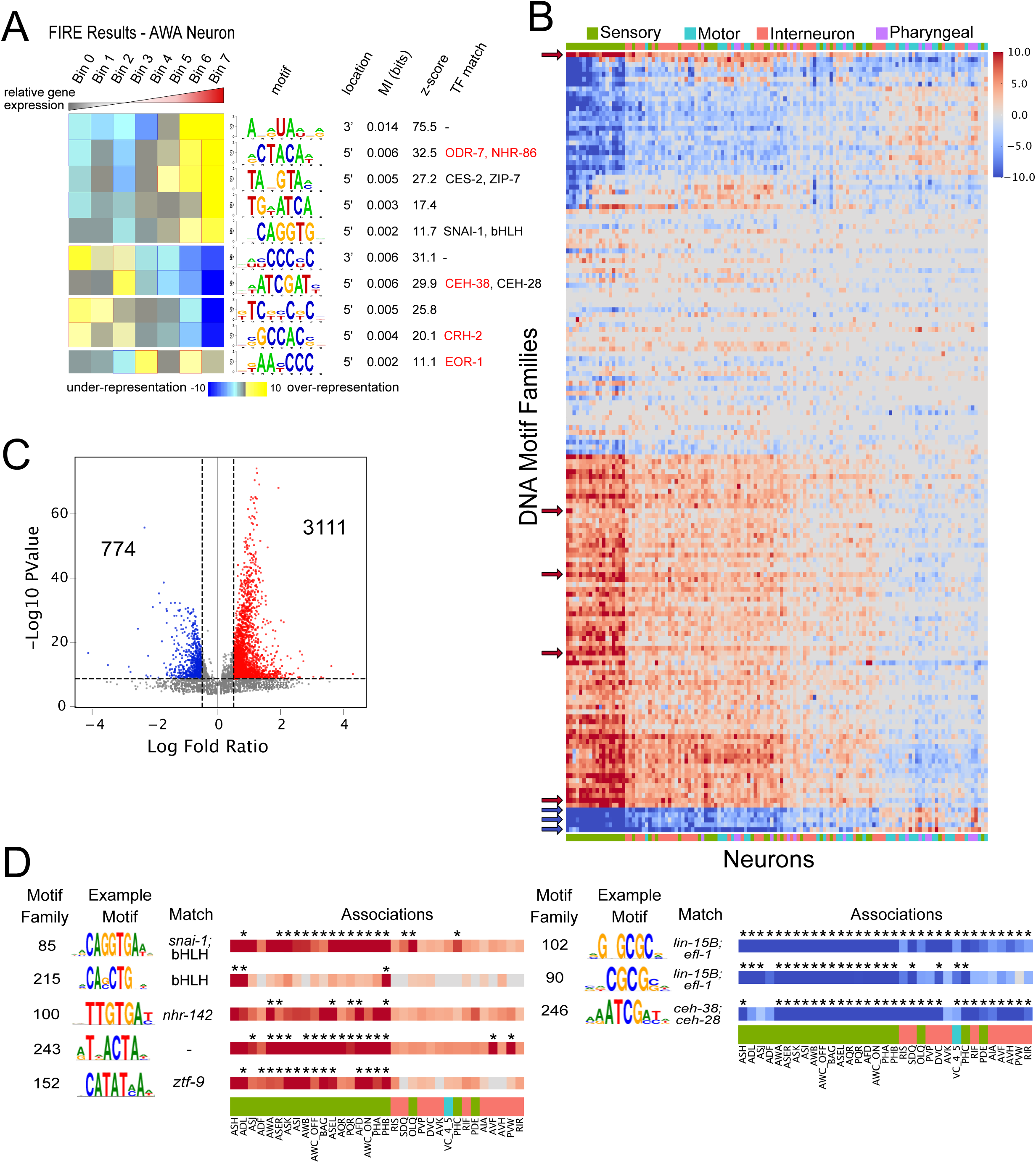
Cis-regulatory elements are abundantly discovered in neuronal transcriptomes. A) FIRE results for the AWA sensory neuron. Genes were grouped into seven bins based on their z-scores from lowest (left) to highest (right). Heatmap denotes over-representation (yellow) or under-representation (blue) of each motif (rows) in genes within each bin. Significant over-representation is indicated by red outlines, whereas significant under-representation is indicated by blue outlines. For each motif, the motif logo, location (5’ promoter or 3’ UTR), mutual information values, z-scores associated with a randomization-based statistical test and matching transcription factors from public databases are listed. Transcription factors in red are expressed in AWA. B) Heatmap showing the enrichment of clustered motifs (rows) in each neuron class (columns). Red color indicates enrichment in genes with highest relative expression, whereas blue color indicates enrichment in genes with lowest relative expression. Color intensity represents log10(p-value) from a hypergeometric test. Motif families and neurons are ordered by similarity. Color bar across x-axis indicates neuron modality. Arrows denote motif families featured in panel D. C) Volcano plot showing log fold ratio and −log10 p-value for all motif family-neuron associations. Numbers within the plot indicate the significant associations with p-value < 1e-5 and log fold ratio > 0.5 (3111) or < −0.5 (774). D) Eight selected motif families with different patterns of positive or negative associations with a subset of neurons from panel C: E-box motifs (motif families 85 and 215), as well as motifs matching NHR TFs (family 100), homeodomain TFs (family 246), and a motif with no known TF matches (family 243). Asterisks denote significant associations. See also Figure S14, S15.

Clustering all discovered motifs based on sequence similarity and overlap of the set of genes with regulatory sequences that include the motif (see Methods), we identified 159 distinct DNA and 65 RNA motif families. A small number (10%) of these motif families came from additional FIRE analyses of co-regulated gene clusters across all neuron types (See Methods, Figure S14A, B). 101 of 159 DNA motif families showed similarity to DNA binding sequences from available databases. For example, FIRE discovered a DNA motif family (core sequence of TAATCC, Figure S14C) which corresponds to the core DNA binding sequence of the K50 class of homeodomain transcription factors (TFs) (Driever and Nüsslein-Volhard, 1989; Treisman et al., 1989). FIRE discovered this motif in genes enriched in ASEL, ASER, AWC^ON^, AWC^OFF^, BAG, and AWA neurons (Figure S14C). The TAATCC sequence matches *in vitro*-derived binding motifs for *C. elegans* K50 class homeodomain genes that are expressed in these neurons (*ceh-36* in ASE and AWC, *ceh-37* in BAG and AWA) (Figure S14C, D) and are required for their development (Chang et al., 2003; Koga and Ohshima, 2004; Lanjuin et al., 2003; Serrano-Saiz et al., 2013). These results indicate that our approach revealed known functional regulatory elements.

To limit false positives, the FIRE algorithm uses stringent criteria for motif discovery and therefore generates conservative results. Although each motif family was discovered in an average of 5 neurons, we reasoned that the identified motif families might also regulate gene expression in additional neuron types. To test for this possibility, we generated motif-neuron associations for each motif family (FIRE figure main B) by calculating the occurrence of motifs in the most enriched genes compared to the most depleted genes for each neuron type (See Methods, FIRE Supplement 1E). In this case, we detected an average of 9 significant neuron associations for each motif family (log fold change > 0.5 and p-value < 1e-5). Results for motif family 184 provide a striking example of how this additional analysis significantly expanded the list of associations for neurons with previously established co-regulated genes. Motif family 184 matches the X-box sequences bound by DAF-19, which regulates cilia formation in all 28 ciliated neuron types (Efimenko et al., 2005; Swoboda et al., 2000). This X-box motif was initially discovered in FIRE runs of 10 ciliated neurons, but was significantly associated with another 12 ciliated sensory neurons by our additional analysis (FIRE Figure Supplement 1G). In another example, we detected additional significant association for the TAATCC motif with the sensory neuron AFD, consistent with the expression of the TAATCC-binding transcription factor *ttx-1*, which is required for AFD development (Satterlee et al., 2001) (FIRE Figure Supplement 1F).

In addition to confirming motifs for known TF targets (e.g., DAF-19), our approach also points to previously undetected roles for TFs in neuron-specific gene regulation. For example, motif family 85 corresponds to the E-box motif CAGGTG, and is strongly associated with nearly all amphid and phasmid neurons (Figure 8D). This particular E-box sequence has been shown to be enriched in *hlh-4* target genes in the nociceptive sensory neuron ADL (Masoudi et al., 2018), but can also bind at least 10 distinct bHLH dimers (Grove et al., 2009). Interestingly, motif family 215 contained a different E-box sequence which was positively associated only with the chemorepulsive sensory neurons ADL, ASH, and PHB (Figure 8D). Based on the expression patterns of bHLH TFs in the adult nervous system, motif 215 may be a target of a HLH-2 homodimer (Masoudi et al., 2018).

Intriguingly, a substantial number of the motifs with strong positive associations with sensory neurons correspond to motifs that do not match known TFs or that match TFs with uncharacterized roles in the nervous system (Figure 8D). For example, motif family 100 showed a strong association with several sensory neurons and is similar to the binding site of the nuclear hormone receptor protein, NHR-142*. nhr-142* is almost exclusively expressed in a subset of amphid sensory neurons (Figure 5A), and the binding domain of *nhr-142* is closely related to several other *nhr* TFs (Lambert et al., 2019) which are expressed primarily in sensory neurons (*nhr-45, nhr-213, nhr-18, nhr-84, nhr-178*) thus suggesting roles for these *nhr* TFs in sensory neuron function. Additionally, several motifs showed strong negative associations with enriched genes across many neurons (Figure 8D, right), indicating possible cis-regulatory elements of transcriptional repressors.

RNA motif analysis revealed that a majority of RNA motif families showed positive associations with many neurons (indicating over-representation of RNA motifs in the enriched genes for each neuron type). Similar to DNA motifs, the strongest effects for RNA motifs were seen in sensory neurons (Figure S15A). In contrast to all other RNA motif families, motif family 23 showed negative associations with most neuron types. This motif family corresponds to a poly-C sequence (Figure S15B). A subclass of RNA binding proteins with KH domains interacts with poly-C regions in RNA and microRNAs (Choi et al., 2009). The *C. elegans* poly-C binding protein HRPK-1 positively regulates the function of several microRNA families, including those that act in the nervous system (Li et al., 2019). The over-representation of the poly-C motif family in depleted genes in most neurons indicates a potential role for this motif in cooperation with microRNA-mediated repression. Overall, our analysis of neuron-specific gene expression across the entire *C. elegans* nervous system identified over 200 cis-regulatory elements that could be sites for *trans*-acting factors such as transcription factors, RNA-binding proteins and microRNAs.

### Cell adhesion molecules (CAMs) are differentially expressed among neurons that are synaptically connected and that define anatomically distinct fascicles in the nerve ring

We compared our transcriptomic data to the *C. elegans* connectome to identify candidate genetic determinants of neurite bundling and synaptic connectivity. For this analysis, we utilized the nerve ring (Figure 9A), the largest expanse of neuropil in the *C. elegans* nervous system, because electron microscope reconstructions from multiple animals have detailed both membrane contacts and synapses in this region (Brittin et al., 2020; Cook et al., 2019; White et al., 1986; Witvliet et al., 2020). We limited our analyses to putative cell adhesion molecules (CAMs), many of which have documented roles in axon pathfinding, selective axon fasciculation and synapse formation within nervous systems (Bruce et al., 2017; Colón-Ramos et al., 2007; Kim and Emmons, 2017; Sanes and Zipursky, 2020; Shen and Bargmann, 2003; Siegenthaler et al., 2015; Spead and Poulain, 2020; Sperry, 1963; Wang et al., 2011). 141 CAMs were detected in neurons in our single cell RNA-Seq dataset (list of CAMs compiled from (Cox et al., 2004; Hobert, 2013), see Table S7).

**Figure 9.**
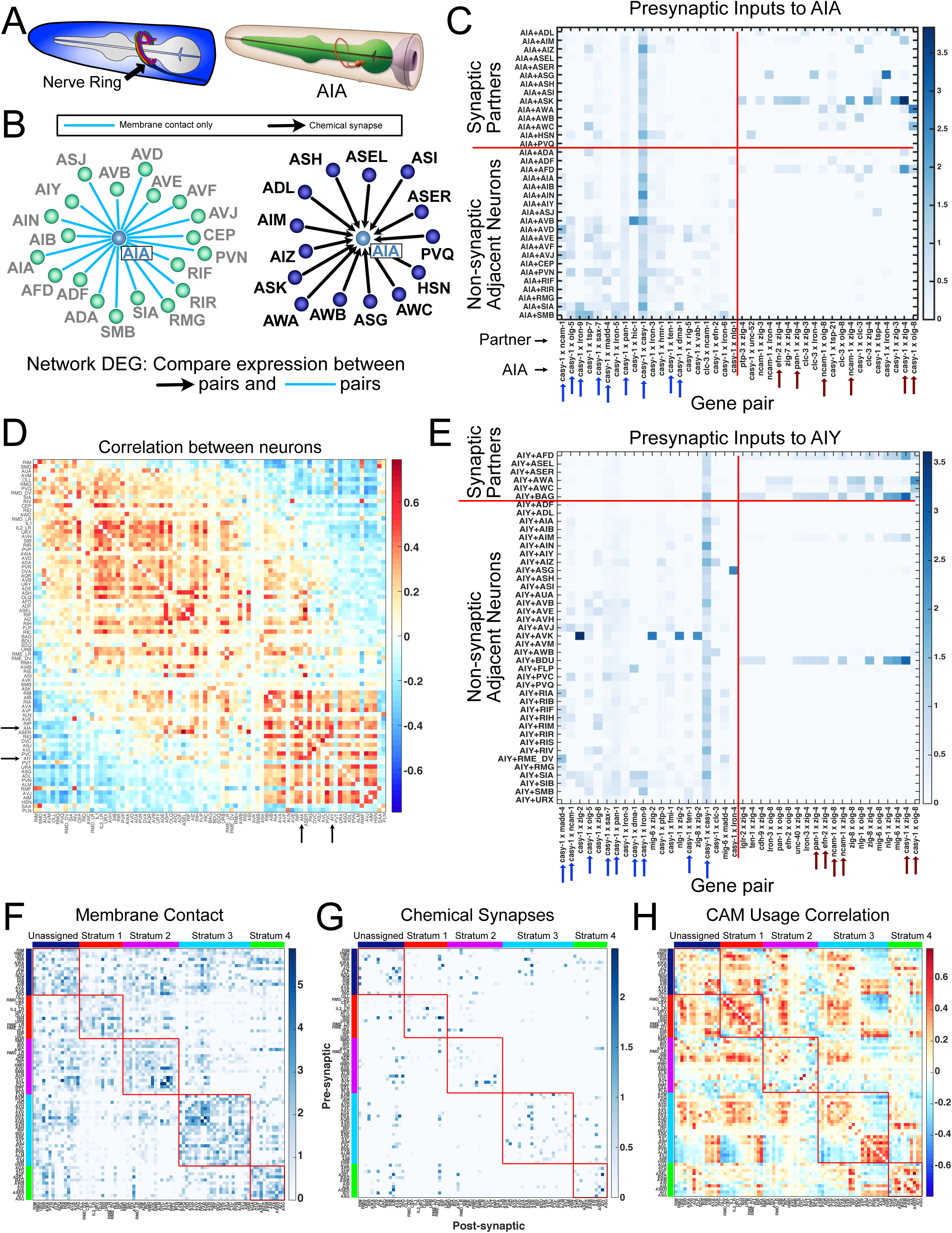
Differential expression of cell adhesion molecules among neurons and their presynaptic partners. A) Left: Cartoon representation of the *C*. *elegans* nerve ring. Right: Diagram of AIA ring interneuron morphology. Courtesy of WormAtlas. B) Diagram of neurons with chemical presynaptic input to AIA (right) and neurons with membrane contact but no synapses with AIA (left). Differential cell adhesion molecule (CAM) gene expression was determined between AIA + synaptic partners vs AIA + non-synaptic adjacent neurons (see Methods). C) Heatmap of the 20 gene pairs with highest log fold change in AIA + presynaptic inputs compared to AIA + non-synaptic adjacent neurons (right of vertical red line). The 20 gene pairs with highest log fold change in AIA + non-synaptic adjacent neurons compared to AIA + presynaptic partners are shown to the left of vertical red line. Blue and dark red arrows indicate gene pairs common among analyses of AIA and AIY (panel E). D) Correlation matrix showing the relationship of CAM usage (see text) across all neurons in the nerve ring (84 neuron types). Neurons split into two main groups. The black arrows indicate AIA and AIY, which showed a correlation of 0.568 with each other. E) Heatmap as in C, for AIY. Six of the top 20 gene pairs (dark red arrows) higher in AIY and its presynaptic partners compared to AIY and adjacent neurons are common with the AIA analysis in C. On the left, 9 of the top 20 genes pairs higher in AIY and adjacent neurons are common with AIA (blue arrows). F) The membrane adjacency matrix was grouped according to nerve ring strata as defined (Moyle et al., 2020). Within each stratum, neurons were ordered according to their CAM usage correlations (see panel H). Membrane contact is generally denser within individual strata than among strata, but not for the unassigned neurons which broadly contact neurons within each stratum. Colored bars indicate neuron strata assignments. Each stratum is outlined by a thin red box. G) The same ordering as in F was imposed upon the chemical connectome revealing that most synapses are detected between neurons within the same stratum. H) The CAM usage correlation matrix (as in D) was grouped by strata, then sorted by similarity (using multidimensional scaling) within each stratum. CAM usage for neurons within stratum 1 is similar to each other, as are those in stratum 4. Stratum 3 shows two distinct populations. See also Figures S16-S19.

Recent computational analysis has revealed a modular structure in which the nerve ring is organized into four distinct bundles of neurites or “strata” as well as a fifth group of unassigned neurons that contacts neurons in multiple strata (Moyle et al., 2020) (Figure S16A). A similar modular organization of the nerve ring has been independently described (Brittin et al., 2020). Nerve ring formation begins in the embryo, but this structure continues to be modified during larval development in which new axons extend into the nerve ring and form synapses (Moyle et al., 2020; Witvliet et al., 2020). In addition, strata and synapses established at early larval stages are maintained throughout the life of the animal. Together, these results point to the importance of both periodic as well as sustained expression of genetic determinants that initiate, modify or maintain the overall structure of the nerve ring and its connectome. We first determined CAMs that were differentially expressed between strata (Figure S16B, C). Six CAMs were significantly enriched in the neurons in one stratum compared to the neurons in all other strata (Figure S16C). Notably, the transcript for MADD-4/punctin, a secreted protein that has been shown to direct process outgrowth as well synaptic placement (Zhou and Bessereau, 2019), is significantly enriched in stratum 1 neurons. In addition, *tsp-7*, which is highly expressed in most neurons of stratum 2, is homologous to the human protein CD63, a member of the tetraspanin superfamily. Tetraspanins interact with integrins and have been implicated in membrane trafficking and synaptogenesis (Murru et al., 2018; Pols and Klumperman, 2009). Two CAM genes, *lron-5* and *lron-9* (extracellular leucine rich repeat proteins) are selectively expressed in a subset of neurons in stratum 2 which could be indicative of roles in organizing these specific fascicles (Figure S16B). Thus, our approach has identified candidate genes that can now be experimentally tested for roles in organizing and maintaining structurally and functionally distinct domains of the nerve ring.

In addition to testing for possible roles in axon fasciculation, we reasoned that expression of specific CAMs among synaptically connected neurons might contribute to synaptic maintenance in the mature nervous system. For our analysis, we surmised that CAMs mediating synaptic stability would be more highly expressed in synaptically connected neurons than in adjacent neurons with membrane contacts but no synapses. We generated high-confidence membrane adjacency and chemical synaptic connectomes by retaining only those contacts and synapses that are preserved across animals in EM reconstructions of the nerve ring (see methods, Table S19, Table S20) (Brittin et al., 2020; Cook et al., 2019; White et al., 1986; Witvliet et al., 2020). These datasets include 84 of the 128 neuron classes. The importance of genetic determinants of connectivity in this circuit is underscored by the observation that membrane contacts between neurons in the nerve ring are much more numerous than synapses; on average, in the nerve ring, each neuron synapses with only 15% of the neurons it contacts (means of 6.42 presynaptic inputs, 6.42 postsynaptic outputs, 42 contacted cells) (Brittin et al., 2020; White et al., 1986).

For each neuron, we compared the expression of all possible combinations of pairs of CAMs in the neuron and its synaptic partners relative to the neuron and its non-synaptic adjacent neurons (Figure 9B, C). Two independent comparisons were generated, one for presynaptic partners (Figure 9C) and a second result for postsynaptic neurons (Figure S17). Our analysis revealed multiple CAM gene pairs with enrichment in synaptically connected neurons compared to adjacent but not synaptically connected neurons (see below). A representative example for presynaptic inputs to the interneuron AIA shows that CAM pairs enriched in synaptically connected neurons were not uniform for the different presynaptic partners of AIA (Figure 9C). For example, AIA and its presynaptic partner, ASK, show strong enrichment for *casy-1* (Calsyntenin) and *zig-4* (secreted 2-Ig domain protein) whereas the AIA-ASG pair is enriched for *casy-1* (Calsyntenin) and *lron-4* (extracellular leucine rich repeat protein). This finding is consistent with the prediction that unique combinatorial codes of CAMs could be required for patterning connectivity between individual pairs of neurons (Kim and Emmons, 2017). Additionally, we identified distinct CAM pairs that are enriched in adjacent, not synaptically connected neurons (Figure 9C). This observation raises the possibility that some of these CAM interactions may functionally inhibit either the formation or maintenance of synapses between neurons. Anti-synaptic effects have been documented for the axon guidance molecules netrin, sema-5B and their cell surface receptors (O’Connor et al., 2009; Poon et al., 2008; Tran et al., 2009).

To examine patterns across the nerve ring, we restricted our analysis to gene pairs with a log fold change > 0.2 in either direction (e.g., enriched in either synaptically connected or in adjacent but not connected neurons) for at least one neuron type. We refer to this pattern of CAM pairs enriched in synaptic or solely adjacent neurons as “CAM usage.” We separately analyzed CAM usage in presynaptic and postsynaptic connections for each neuron. Of the 19,881 possible CAM pairs, 439 pairs passed our log fold change threshold for presynaptic connections (Figure S18A), whereas 443 pairs showed > 0.2 log fold change for postsynaptic connections. (Figure S18B). To identify neurons with similar patterns of CAM usage, we generated correlation matrices from pairwise comparisons of all neurons and sorted neurons by similarity using multidimensional scaling (presynaptic CAM usage in Figure 9D, postsynaptic CAM usage in Figure S17D). For example, CAM usage for presynaptic inputs to AIA and AIY is strongly correlated (correlation 0.568) due to the co-occurrence for each neuron of multiple shared combinations of CAMs (Figure 9E, blue and red arrows). This analysis also separated neurons into two main groups based on CAM usage which could be indicative of underlying shared roles for CAMs among these distinct sets of neurons.

We sought to understand the relationship between stratum membership and synaptic CAM usage for nerve ring neurons. Both membrane contact and chemical synapses are denser among neurons within strata than across strata (Figure 9F, G, Figure S17F, G), a finding also observed for an independent assessment of nerve ring axon bundles (Brittin et al., 2020). We sorted neurons by CAM usage within each stratum (Figure 9H – presynaptic, Figure S17H – postsynaptic) to assess intra-stratum correlations. This approach revealed high correlations among neurons within strata. Additionally, neurons in some strata split into distinct groups based on CAM usage (Stratum 3 in both pre- and postsynaptic analyses, Strata 2 and 4 in postsynaptic analysis, Figure 9H, S17H). This observation suggests that CAM usage at synaptic connections is likely distinct from CAMs that may be involved in strata formation and/or maintenance. We also note although CAM usage correlations were often elevated among neurons within strata, high correlations were also detected among neurons in different strata that are not synaptically connected and with minimal contacts thus suggesting the potential synaptogenic effects for CAMs likely depend on additional factors. We suggest that that the overall results of our analysis point to specific CAMs that can now be investigated for roles in the formation and maintenance of synapses as well as fasciculation between specific neurons in the *C. elegans* nerve ring.

### Data interface

We developed a web application, CengenApp (http://cengen.shinyapps.io/CengenApp) to facilitate analysis of the scRNA-Seq data produced by this project. Users can generate gene expression profiles by neuron class or by gene at different thresholds, and perform differential gene expression analysis between either individual neurons or between groups of neuron types. In addition, an interactive graphical interface is available for generating heat map representations (e.g., Figure 4C) of gene expression across the nervous system. Raw data have been deposited with the Gene Expression Omnibus (www.ncbi.nlm.nih.gov/geo) (GSE136049). The data as an R object and additional supporting files can be downloaded from the CeNGEN website, www.cengen.org. Finally, code used in the analyses presented here is available at Github, www.github.com/cengenproject.

## CONCLUSIONS

We have produced a gene expression map for the mature *C. elegans* nervous system. To our knowledge, these results constitute the first transcriptome of every neuron type in an entire nervous system and complement earlier partial profiles of the *C. elegans* nervous system at embryonic and early larval stages (Cao et al., 2017; Packer et al., 2019). This complete catalog of neuronal gene expression provides an essential foundation for a comprehensive exploration of transcriptional and gene regulatory patterns that lead to neuronal diversity, connectivity and function. To aid our analysis, we developed a novel thresholding approach for single-cell data to generate high confidence profiles for each neuron type. Our findings indicate that neuropeptide signaling is widely utilized throughout the entire *C. elegans* nervous system, and likely crucial for a variety of functions. Multiple lines of evidence support these conclusions. First, neuropeptide-encoding genes are among the most abundantly detected genes in the dataset. Second, at the most stringent threshold examined, each neuron expresses at least four different neuropeptide-encoding genes. Third, each neuron expresses a unique combination of both neuropeptide genes and putative neuropeptide receptors. Our findings are consistent with recent reports of abundant and widespread neuropeptide expression in *Hydra* (Siebert et al., 2019), *Drosophila* (Allen et al., 2020) and mouse cortical neurons (Smith et al., 2019), indicating that that these salient features of neuropeptide signaling are conserved across phyla.

Our analysis of transcription factor expression across an entire nervous system reveals that different transcription factor families appear to have segregated into distinct functions during cellular differentiation. Some families are underrepresented in the mature nervous system (T-box genes), others show very broad expression patterns in the nervous system (Zn finger), while others are sparsely expressed and appear to exquisitely track with neuronal identity (homeodomains) (Reilly et al., 2020). The nuclear hormone receptors may have acquired a unique function in sensory neuron biology, as inferred by their striking enrichment in sensory neurons. The identification of enriched cis-regulatory motifs in neuronal gene batteries provides an opportunity for future experiments that will dissect the mechanisms of gene regulation in the nervous system.

Finally, we devised computational strategies that exploit our gene expression profile of the *C. elegans* nervous system to reveal the genetic underpinnings of neuron-specific process placement and connectivity. Previous computational efforts to forge a link between neuron-specific gene expression and the *C. elegans* wiring diagram have been hampered by incomplete and largely qualitative expression data (Barabási and Barabási, 2020; Baruch et al., 2008; Kaufman et al., 2006; Kovacs et al., 2020; Varadan et al., 2006). Here, we leveraged our nervous-system wide catalog of gene expression to deduce combinatorial codes for cell adhesion protein genes that likely contribute to the maintenance and formation of this complex neuropil. Importantly, this analysis can now be extended to unique groups of neurons and to any gene family to generate specific hypotheses of connectivity for direct experimental validation.

We anticipate that these data will be useful for future studies of individual genes, neurons, and circuits, as well as global analyses of an entire nervous system. Additionally, we anticipate that these data will be of use for the development of single-cell RNA sequencing analysis methods. Coupled with the fully described cell lineages (Sulston and Horvitz, 1977; Sulston et al., 1983), neuronal anatomy (Albertson and Thomson, 1976; Brittin et al., 2020; Cook et al., 2019; White et al., 1986; Witvliet et al., 2020), and powerful functional analyses, such as pan-neuronal calcium imaging and neuronal identification (Kato et al., 2015; Kotera et al., 2016; Nguyen et al., 2016; Venkatachalam et al., 2016; Yemini et al., 2019), our dataset provides the foundation for discovering the genetic programs underlying neuronal development, connectivity and function.

## Acknowledgements

We acknowledge HaoSheng Sun for examining neuropeptide reporter strains. We thank the CeNGEN Advisory Board for helpful suggestions and M. Zhen for providing the AVG marker strain, ZM9592. This work was funded by NIH grant R01NS100547 to MH, OH, DMM, and NS and by Vanderbilt Trans-Institutional Program funds to DMM. Flow Cytometry experiments were performed in the Vanderbilt Flow Cytometry Shared Resource which is supported by the Vanderbilt Ingram Cancer Center (P30 CA68485) and the Vanderbilt Digestive Disease Research Center (DK058404). The Vanderbilt VANTAGE Core provided technical assistance for this work and is supported by CTSA Grant (5UL1 RR024975-03), the Vanderbilt Ingram Cancer Center (P30 CA68485), the Vanderbilt Vision Center (P30 EY08126), and NIH/NCRR (G20 RR030956). Some strains were provided by the CGC, which is funded by NIH Office of Research Infrastructure Programs (P40 OD010440). G.S. received the support of a fellowship from “la Caixa” Foundation (ID 100010434). The fellowship code is LCF/BQ/PI19/11690010. G.S. is also supported by Ministerio de Ciencia e Innovación, Spain (PID2019-104700GA-I00).

## Author Contributions

M.H., O.H., N. S., D.M.M. originated project.

S.R.T. generated scRNA-Seq data, performed cell calling, batch correction, UMAP and clustering analyses, assigned neuron identities to clusters, analyzed gene family expression patterns, helped A.W., C.X., E.V. and P.O. with data analysis, designed figures and supplemental tables, wrote the original draft, edited final version.

G.S. developed CengenApp with input from the group.

A.W. designed thresholding strategy with input from S.R.T., analyzed alternative splicing and implemented web application for meta data.

A.B. generated bulk RNA sequence data and compared with scRNA-Seq results.

M.R. generated reporter-based expression data for identifying neuron clusters and for implementing thresholding strategy, performed exploratory analysis for transcription factor family expression patterns.

C.X. refined gene models with extended 3’ UTRs for read mapping and expression level quantification, performed evolutionary comparisons.

E.V. developed computational strategy to identify cell adhesion proteins correlated with neuron-specific synapses and process placement, input on statistical and quantitative analysis.

P.O. Implemented FIRE analysis to identify candidate cis-regulatory motifs.

L.G. organized and managed BrainAtlas.

R.M. used FACS to isolate neurons and extracted RNA for bulk RNA seq.

A.P. used FACS to isolate neurons.

I.R., E.Y., S.J.C., B.V., C.C., M.B., A.A., S.R.T. generated and analyzed reporter strains.

E. Y. provided the NeuroPAL strain and assisted with neuron identification.

S.T. provided guidance for FIRE analysis conducted by P.O.

N.S. directed work of G.S. and C.X. and provided guidance for evolutionary comparisons

M.H. directed work of G.S., A.W., A.B., M.B. and A.A., contributed to original draft and helped edit final version.

O.H. directed work of M.R., L.G., I.R., E.Y., S.J.C., B.V., C.C., contributed to original draft and helped edit final version.

D.M.M oversaw scope of work, directed efforts of S.R.T., R.M., A.P., contributed to original draft and edited final version.

## STAR Methods

### RESOURCE AVAILABILITY

#### Lead Contact

Requests for resources and reagents should be directed to the Lead Contact, David Miller

#### Materials Availability

##### Data and Code Availability

The raw data are available at GEO (Accession Number 136049). The full and neuron only datasets are available at www.cengen.org. Analysis code is available at github https://github.com/cengenproject.

### Nematode Strains

The strains used in the study are listed in Table S1.

## EXPERIMENTAL MODEL AND SUBJECT DETAILS

### Preparation of larval strains and dissociation

Worms were grown on 8P nutrient agar 150 mm plates seeded with *E. coli* strain NA22. To obtain synchronized cultures of L4 worms, embryos obtained by hypochlorite treatment of adult hermaphrodites were allowed to hatch in M9 buffer overnight (16-23 hours at 20° C) and then grown on NA22-seeded plates for 45-48 hours at 23° C. The developmental age of each culture was determined by scoring vulval morphology (>75 worms) (Mok et al., 2015). Single cell suspensions were obtained as described (Kaletsky et al., 2016; Spencer et al., 2014; Zhang et al., 2011) with some modifications. Worms were collected and separated from bacteria by washing twice with ice-cold M9 and centrifuging at 150 rcf for 2.5 minutes. Worms were transferred to a 1.6 mL centrifuge tube and pelleted at 16,000 rcf for 1 minute. 250 µL pellets of packed worms were treated with 500 µL of SDS-DTT solution (20 mM HEPES, 0.25% SDS, 200 mM DTT, 3% sucrose, pH 8.0) for 2-4 minutes. In initial experiments, we noted that SDS-DTT treatment for 2 minutes was sufficient to dissociate neurons from the head and tail, but longer times were required for effective dissociation of neurons in the mid-body and ventral nerve cord. The duration of SDS-DTT was therefore selected based on the cells targeted in each experiment. For example, NC3582, OH11746, and *juIs14* L4 larvae were treated for 4 minutes to ensure dissociation and release of ventral cord motor neurons. NC3579, NC3580 and NC3636 L4 larvae were treated with SDS-DTT for 3 minutes. All other strains were incubated in SDS-DTT for 2 minutes. Following SDS-DTT treatment, worms were washed five times by diluting with 1 mL egg buffer and pelleting at 16,000 rcf for 30 seconds. Worms were then incubated in pronase (15 mg/mL, Sigma-Aldrich P8811, diluted in egg buffer) for 23 minutes. During the pronase incubation, the solution was triturated by pipetting through a P1000 pipette tip for four sets of 80 repetitions. The status of dissociation was monitored under a fluorescence dissecting microscope at 5-minute intervals. The pronase digestion was stopped by adding 750 µL L-15 media supplemented with 10% fetal bovine serum (L-15-10), and cells were pelleted by centrifuging at 530 rcf for 5 minutes at 4 C. The pellet was resuspended in L-15-10, and single-cells were separated from whole worms and debris by centrifuging at 100 rcf for 2 minutes at 4 C. The supernatant was then passed through a 35-micron filter into the collection tube. The pellet was resuspended a second time in L-15-10, spun at 100 rcf for 2 minutes at 4 C, and the resulting supernatant was added to the collection tube.

### FACS isolation of fluorescently-labeled neuron types for RNA sequencing

Fluorescence Activated Cell Sorting (FACS) was performed on a BD FACSAria™ III equipped with a 70-micron diameter nozzle. DAPI was added to the sample (final concentration of 1 µ g/mL) to label dead and dying cells. To prepare samples for scRNA-sequencing, our general strategy used fluorescent reporter strains to isolate subgroups of cells. For example, we used an *eat-4::mCherry* reporter (OH9625) to target glutamatergic neurons and an *ift-20::NLS-TagRFP* reporter (OH11157) to label ciliated sensory neurons. We used an intersectional labeling strategy with a nuclear-localized pan-neural marker (*otIs355 [rab-3(prom1)::2xNLS-TagRFP]* IV) to exclude cell fragments labeled with cytosolic GFP markers (NC3582). In other cases, we used an intersectional strategy to exclude non-neuronal cells. For example, *stIs10447* [*ceh-34p::HIS-24::mCherry*] is expressed in pharyngeal muscles, pharyngeal neurons and coelomocytes. To target pharyngeal neurons, we generated strain NC3583 by crossing *stIs10447 [ceh-34p::HIS-24::mCherry*] with the pan-neural GFP marker *evIs111* to isolate cells that were positive for both mCherry and GFP. Non-fluorescent N2 (wild-type reference strain) (Brenner, 1974) standards and single-color controls (in the case of intersectional labeling approaches) were used to set gates to exclude auto-fluorescent cells and to compensate for bleed-through between fluorescent channels. For two experiments, single-cell suspensions from separate strains were combined (OH16003 plus PS3504 and *nIs175*, NC3635 plus NC3532) prior to FACS. In some cases, we expanded FACS gates to encompass a wide range of fluorescent intensities to ensure capture of targeted cell types. This less stringent approach may contribute to the presence of non-neuronal cells in our dataset (see Results). Cells were sorted under the “4-way Purity” mask.

For 10X Genomics single-cell experiments, sorted cells were collected into L-15-33 (L-15 medium containing 33% fetal bovine serum), concentrated by centrifugation at 500 rcf for 12 minutes at 4° C, and counted on a hemocytometer. Single-cell suspensions used for 10x Genomics single-cell sequencing ranged from 300-900 cells/µL.

For bulk RNA-sequencing of individual cell types, sorted cells were collected directly into TRIzol LS. At ∼15-minute intervals during the sort, the sort was paused, and the collection tube with TRIzol was inverted 3-4 times to ensure mixing. Cells in TRIzol LS were stored at −80° C for RNA extractions (see below).

### Single-cell RNA sequencing

Each sample (targeting 5,000 or 10,000 cells per sample) was processed for single cell 3’ RNA sequencing utilizing the 10X Chromium system. Libraries were prepared using P/N 1000075, 1000073, and 120262 following the manufacturer’s protocol. The libraries were sequenced using the Illumina NovaSeq 6000 with 150 bp paired end reads. Real-Time Analysis software (RTA, version 2.4.11; Illumina) was used for base calling and analysis was completed using 10X Genomics Cell Ranger software (v3.1.0). Most samples were processed with 10x Genomics v2 Chemistry, except for samples from *juIs14*, NC3583, NC3636, CX5974, OH16003, PS3504, *nIs175*, NC3635 and NC3532, which were processed with v3 Chemistry. Detailed experimental information is found in Table S2.

### Single-cell RNA-Seq Mapping

Reads were mapped to the C. elegans reference transcriptome from WormBase, version WS273. Due to the possibility that 3’ untranslated region (UTR) annotations in the reference transcriptome may be too short (Packer et al., 2019), we dynamically extended the 3’ UTR of each gene to its optimal length, thereby enabling the additional mapping of reads to the 3’ extremity of the gene body. We generated eight versions of gene annotations based on WormBase WS273 annotation, with 3’ UTRs in each version elongated by 50, 100, 150, 200, 250, 300, 400 and 500 base pairs (bps), respectively. Elongation of genes which overlapped with other genes during the extension process was terminated before encountering an adjacent exon. Subsequently, eight custom genome indexes, which respectively combined the *C. elegans* WS273 reference genome with the eight extended gene annotation versions, were generated using CellRanger (version 3.1.0).

All sequenced reads from each of the 17 single-cell samples were mapped to the eight reference genomes using the CellRanger pipeline. We next selected the best UTR extension length of each annotated gene independently for the 17 samples, as a number of genes were heavily enriched in specific samples. First, we calculated the total number of mapped reads for each of the expressed genes in each sample, resulting in eight mapped-read values representing the eight gene annotation versions. To discard the UTR extension intervals which harbor sparse additional reads, as well as to allow for the intervals which harbor fewer reads but are surrounded by read-enriched intervals, we took advantage of the trimming algorithm in Burrows-Wheeler Alignment (Li and Durbin, 2009) to find the best extension. Specifically, a cutoff of 20 reads was applied to each extension interval (50, 50, 50, 50, 50, 100, and 100 bps). Cumulative sums from 3’ to 5’ end were then calculated after subtracting the cutoff in each interval, and the smallest sum of less than 0 was located as the trimming point for a given sample. Considering all 17 samples, the trimming point agreed by most samples (or at least two samples if one gene is expressed in limited samples) was chosen as the ultimate one. Consequently, we extended the UTRs for 1,012 *C. elegans* genes, encompassing 40, 216, 175, 113 and 468 genes with UTRs extended by 150, 200, 250, 300 and 400 bps at the 3’ end, respectively. Lastly, with the gene annotation file containing the optimal extension length for each gene, we remapped and quantified the gene expression in all 17 samples using CellRanger.

### Downstream Processing

We distinguished cells from empty droplets, corrected background RNA expression and generated quality control metrics for each sample independently, then merged the files together into one dataset. The default barcode filtering algorithm in CellRanger can fail to capture cells in some conditions, especially with cells with variable sizes and RNA content (Lun et al., 2019). Neurons in particular tend to have lower UMI counts than other cell types and can be missed by the default algorithm (Packer et al., 2019). We therefore used the EmptyDrops method (with a threshold of 50 UMIs for determining empty droplets) from the R package DropletUtils (Lun et al., 2019) to determine which droplets contained cells. This approach detected significantly more cells than the CellRanger method, and we were able to confidently annotate these additional cells as neurons.

The SoupX R package (Young and Behjati, 2020) was used to correct for background RNA. We used a more conservative threshold for determining background RNA for SoupX than for EmptyDrops to exclude low-quality cells in the background correction. We therefore set a threshold of droplets with fewer than 25 UMIs to estimate the background RNA. Genes with patterns of strong expression in restricted sets of cells (from the literature or from preliminary clustering analysis for each single-cell experiment) were selected for each dataset (Table S2). SoupX uses these genes, preliminary clustering, and the calculated background RNA profile (from droplets with fewer than 25 UMIs) to estimate the percent of contamination in each sample. The estimated background contamination ranged from 4.15-13.56%, with a mean of 8.01%. For the *ceh-28_dat-1* experiment, no combination of genes tested resulted in satisfactory performance, so the contamination was set manually to 10.00%. SoupX uses the calculated contamination level to correct the expression of genes that are abundant in the background RNA profile, and returns a corrected gene by cell count matrix. The background corrected count matrices produced by SoupX were rounded to integer counts and used for subsequent downstream processing.

Following background correction, quality control metrics were calculated for each dataset with the R package scater (McCarthy et al., 2017), using the percentage of UMIs from the mitochondrial genes *nduo-1, nduo-2, nduo-3, nduo-4, nduo-5, nduo-6, ctc-1, ctc-2, ctc-3, ndfl-4, atp-6,* and *ctb-1*. Droplets with greater than twenty percent of UMIs coming from mitochondrial genes were removed. Datasets from individual experiments were merged using Seurat (v3) (Stuart et al., 2019). Genes detected in fewer than five cells were removed. Log-normalized expression matrices were then used for downstream analysis using monocle (2.99.3), monocle3 (0.2.1) (Cao et al., 2019; Qiu et al., 2017a, 2017b; Trapnell et al., 2014) and Seurat (v3) packages.

### Dimensionality reduction and batch correction

We imported the merged dataset into monocle3, and reduced the dimensionality of the dataset with PCA (135 principal components, based on examination of an elbow plot showing the variance explained by each principal component), followed by the Uniform Manifold Approximation and Projection (UMAP) (Becht et al., 2019; McInnes et al., 2018) algorithm in monocle3 (reduce_dimension function, parameters were default other than: umap.min_dist = 0.3, umap.n_neighbors = 75). We then clustered cells using the Louvain algorithm in monocle3 (res = 3e-4). Batch correction between experiments was performed using the align cds function (Cao et al., 2019; Haghverdi et al., 2018). We processed the neuron-only dataset with the following parameters (125 PCs, umap.min_dist = 0.3, umap.n_neighbors = 75, alignment_k (for align_cds) = 5, clustering resolution 3e-3).

We assigned tissue and cell identity to the majority of cells in our dataset based on a manually compiled list of reported gene expression profiles with an average of > 20 molecular markers per neuron type (Hobert et al., 2016), and a recently described protein expression atlas of >100 homeodomain proteins (Reilly et al., 2020) (Table S3). We manually excluded clusters we identified as doublets due to co-expression of cell-type specific markers. We manually merged multiple clusters that corresponded to the same neuron type. We noted that coelomocytes were most abundant in experiments using strains expressing mCherry (*otIs292* and *otIs447)*. This effect likely results from neurons shedding mCherry+ exophers, which are then taken up by coelomocytes (Melentijevic et al., 2017), causing them to be isolated along with mCherry-labeled neurons.

Some clusters in the initial global dataset appeared to contain multiple closely related neuron types (i.e., cholinergic motor neurons, dopaminergic neurons, oxygen sensing neurons AQR, PQR, URX and pharyngeal neurons). Additional analysis of these separate clusters (i.e., reapplication of PCA, UMAP, and Louvain clustering to just these clusters) separated these cell types into individual clusters. Finally, we identified separate clusters for the neuron classes RIV and SMD. In both instances, however, one of the putative clusters showed strong expression of stress-related transcripts rather than sub-type specific markers and therefore likely correspond to a subset of RIV and SMD neurons damaged by the isolation protocol. These two aberrant clusters were excluded from further analyses.

### Neuron network analysis

The neuron network containing all neuron types was constructed on the basis of the transcriptome similarity between each pair of neuron types. We obtained the transcriptional profile of each neuron type by averaging gene expression across all cells within the given type, resulting in the gene expression trajectory for each neuron type. We next calculated transcriptome similarity (after log transformation) as the Pearson correlation coefficient between pairwise neuron types, using 7,390 highly variable genes identified by Seurat based on their variance and mean expression. The neuron network in a graphopt layout was constructed by the package “igraph” (Csárdi and Nepusz 2006) in R using the force-directed graphopt algorithm based on the above similarity matrix.

### Gene expression analyses

Averaged gene expression profiles for each neuron class were generated as described (Cao et al., 2017). Quantitative expression data for a subset of genes are distorted by overexpression from fosmid reporters or co-selectable markers (*lin-15A, lin-15B, pha-1, rol-6, unc-119, dpy-20, cho-1*), the promoter regions used for marking cell types (*unc-53, unc-47, gcy-35)* or from a gene-specific 3’ UTR included in fluorescent reporter constructs (*eat-4*, *unc-54*). These genes are annotated in the CengenApp web application.

For visualization of gene expression data in the web application and for differential gene expression tests, data were imported into Seurat (v3) and raw counts were normalized using the variance stabilizing transformation (VST) implemented in the function sctransform with default parameters and regressing out the percent of mitochondrial reads (Hafemeister and Satija, 2019; Stuart et al., 2019). Differential gene expression tests used the Seurat v3 default Wilcoxon rank sum test with default parameters (a gene must be detected in > 10% of the cells in the higher-expressing cluster and have an adjusted p-value < 0.05).

### Thresholding

The wealth of known gene expression data in *C. elegans* from fluorescent reporter strains provides a unique opportunity to set empirical thresholds for our scRNA-Seq data based on ground truth. We first compiled a ground truth dataset of 160 genes with expression patterns across the nervous system previously determined with high confidence fosmid fluorescent reporters, CRISPR strains or other methods (Bhattacharya et al., 2019; Harris et al., 2020; Reilly et al., 2020; Stefanakis et al., 2015; Yemini et al., 2019) (Table S5). For each gene, we then aggregated expression across the single cells corresponding to each neuron type and calculated several metrics, including the total UMI count, the number of single cells of each neuron type in which each gene was detected with at least one UMI, the proportion of single cells of each neuron type in which gene was detected with at least one UMI and a normalized transcripts per million (TPM) expression value. We generated receiver operating characteristic (ROC) and precision recall (PR) curves for each metric by thresholding the data across a range of values, and calculated true positive, false positive, and false discovery rates by comparing the single-cell data to the ground truth. We used the area under the curve to decide which metric to use for thresholding. The proportion of cells in which a gene was detected performed the best (had the highest AUC) and was thus used to establish gene-level thresholds.

We first set initial thresholds to retain ubiquitously-expressed genes and to remove non-neuronal genes. Genes detected in ≥ 1% of the cells in every neuron cluster were considered expressed in all neuron types (193 genes), whereas transcripts detected in ≤ 2% of the cells in every neuron cluster were considered non-neuronal (4806 genes; no genes were detected in ≥ 1 % and ≤ 2 % of the cells in every neuron). As most genes displayed different levels of expression, we found that a single threshold failed to reliably capture expression for all genes. Thus, we applied percentile thresholding for each gene individually. For example, the AFD cluster showed the highest proportion of cells (76.3%, Figure S6A) expressing the homeodomain transcription factor *ttx-1*. For *unc-25*/GAD, the VD_DD cluster had the highest proportion of cells (94.4%, Figure S6G), whereas for the homeodomain transcription factor *ceh-13*, the DA neuron cluster had the highest proportion (13.4%, not shown). Thresholds were calculated as a fraction of the highest proportion of cells for each individual gene. For example, a threshold of 0.04 results in different absolute cut-offs for each gene. For *ttx-1,* with a highest proportion of 76.3%, we scored *ttx-1* as “not expressed” in clusters in which it was detected in < 3.05% of cells (0.04*76.3 = 3.05%). For *unc-25,* with a highest proportion of expressing cells of 94.4%, we scored *unc-25* as “not expressed” in clusters in which it was detected in < 3.77% of cells (0.04*94.4 = 3.77%). Similarly, and we scored *ceh-13* as “not expressed” in clusters in which it was detected in < 0.536% of cells (0.04*13.4 = 0.536%).

For each threshold percentile, we generated 5,000 stratified bootstraps of the ground truth genes using the R package boot (Canty and Ripley, 2019; Davison and Hinkley, 1997) and computed the True Positive Rate (TPR), False Positive Rate (FPR) and False Discovery Rate (FDR) for the entire dataset as well as for each neuron type. We estimated 95% confidence intervals with the adjusted percentile (BCa) method, and plotted the ROC and PR curves (Figure S6C, D). Finally, we selected 4 thresholds of increased stringency (1-4, see Table S6 for statistics for each neuron type). Threshold 2 was used for analyses profiling gene expression across all neuron types and across gene families.

### Determining unique combinations of gene families

Expression matrices of selected gene families from threshold 2 were binarized. Genes were clustered following default parameters in the R package hclust. We determined if neurons expressed a unique combinatorial code for given gene families by determining whether any two columns (neurons) of the binarized expression matrix were identical.

### Connectivity Analysis

To determine neurons postsynaptic to either ACh or glutamate-releasing neurons, we used the *C. elegans* hermaphrodite chemical connectome data from (Cook et al., 2019). For this analysis, we scored synapses as connections detected in more than 3 electron micrograph sections.

### Reporter strains

GFP reporters for the neuropeptide genes *flp-33, nlp-17, nlp-42, nlp-52* and *nlp-*56 were created by PCR Fusion (Hobert, 2002) whereby the 5’ intergenic region of the gene of interest and the coding sequence of GFP with 3’ UTR of *unc-54* were fused in subsequent PCR reactions. We used the entire intergenic region of the genes of interest: 1519 bp for *flp-33* (forward primer: aggaagttgataaacttgcttgttttaatg, reverse primer: ggtagggggaccctggaag), 372 bp for *nlp-17* (forward primer: tcatctaaaatatattttcaaaacgattttctgtgc, reverse primer: attttctgtgaaaaagcctgactttttc), 3250 bp for *nlp-42* (forward primer: ttgtctgaaaatatgggttttgcatgg, reverse primer: tttacctgaaaatttgcaatttttcagatttttac), 3731 bp for *nlp-52* (forward primer: ttgcttgcattttctgaaataagatgg, reverse primer: ttttgggaagaggtacctggaac), and 2954 bp for *nlp-56* (forward primer: ggttcactggaataaatatatgcactgtatc, reverse primer: ctggaagagttgaatcatatggtttagaag). Reporters were injected directly into NeuroPAL *pha-1* strain (OH15430 *pha-1(e2123); otIs669[NeuroPAL 15])* (Yemini et al., 2019) as a complex array with OP50 DNA (linearized with ScaI) and *pBX [pha-1 (+)]* (Granato et al., 1994) as a co-injection marker. For *flp-33* and *nlp-52*, the reporter, *pBX [pha-1 (+)]* and OP50 DNA were injected at concentrations of 7.75 ng/µl, 6.2 ng/µl, 99.96 ng/µl, respectively. For *nlp-42*, the reporter, *pBX [pha-1 (+)]* and OP50 DNA were injected at 11.80 ng/µl, 8.7 ng/µl and 88.86 ng/µl. For *nlp-17*, the reporter, *pBX [pha-1 (+)]* and OP50 DNA were injected at 10 ng/µl, 6.2 ng/µl and 99.96 ng/µl. For *nlp-56*, the reporter, *pBX [pha-1 (+)]* and OP50 DNA were injected at concentrations of 9.5 ng/µl, 5.2 ng/µl and 94.9 ng/µl. After injection, animals were kept at 25°C for selection of the array positive worms and maintained for at least three generations before imaging (see below). CRISPR reporter strains for *flp-1* and *nlp-51* were generated by engineering a T2A::3xNLS::GFP cassette into the respective gene loci just before the stop codons. The *nspc-1* promoter fusion reporter was constructed using the entire 713 bp intergenic region upstream of *npsc-1* fused driving GFP.

Sequences of C39H7.2 and *nhr-236* were acquired from *C. elegans* BioProject PRJNA13758 browser (via WormBase). We combined 1447 bp upstream of the C39H7.2 sequence (forward primer: Gtatggtctgcaggagtatc, reverse primer: Gcccatggaagtgtcgaatt) with 2044 bp of UberPN::3xNLS-intronGFP (forward primer: CCCAAAGgtatgtttcgaat, reverse primer: AACTGTTTCCTACTAGTCGG) via overlap PCR. For *nhr-236*, we combined 802 bp immediately upstream of the ATG sequence of the first exon of *nhr-236* (forward primer: Tcttgaagggcacgccgatt, reverse primer: Gctctgtgtcggtattccgg) with 2044 bp of UberPN::3xNLS-intronGFP (primers as above) via overlap PCR. The resulting overlap PCR products were injected with 50 ng/µl of *pha-1* rescue construct *pBX [pha-1 (+)]* and 1Kb+ladder (Promega Corporation, G5711) into GE24 [*pha-1(e2123)* III]. The injected lines were grown at 25 C for selection of the *pha-1+* worms and were maintained for at least five generations before imaging with a Spinning Disk Confocal microscope (Nikon). The images were analyzed using Volocity Imaging Software and also crossed into the NeuroPAL strain *otIs669* to identify the neurons expressing the reporters.

### Imaging

Confocal images were obtained on either a Nikon A1R confocal laser scanning microscope or a Zeiss LSM 880 microscope using 20x or 40x oil immersion objectives. Brightness and contrast adjustments were performed with FIJI.

### RNA Extraction

Cell suspensions in TRIzol LS (stored at −80° C) were thawed at room temperature. Chloroform extraction was performed using Phase Lock Gel-Heavy tubes (Quantabio) according to the manufacturer’s protocol. The aqueous layer from the chloroform extraction was combined with an equal volume of 100% ethanol and transferred to a Zymo-Spin IC column (Zymo Research). Columns were centrifuged for 30 sec at 16,000 rcf, washed with 400 µL of Zymo RNA Prep Buffer and centrifuged for 16,000 rcf for 30 sec. Columns were washed twice with Zymo RNA Wash Buffer (700 µL, centrifuged for 30 sec, followed by 400 µL, centrifuged for 2 minutes). RNA was eluted by adding 15 µL of DNase/RNase-Free water to the column filter and centrifuging for 30 sec. A 2 µL aliquot was submitted for analysis using the Agilent 2100 Bioanalyzer Picochip to estimate yield and RNA integrity and the remainder stored at −80° C.

### Bulk sequencing and mapping

Each bulk RNA sample was processed for sequencing using the SoLo Ovation Ultra-Low Input RNAseq kit from Tecan Genomics according to manufacturer instruction, modified to optimize rRNA depletion for *C. elegans* (A. Barrett, R. McWhirter, S. Taylor, A. Weinreb, D. Miller, and M. Hammarlund, personal communication). Libraries were sequenced on the Illumina Hiseq 2500 with 75 bp paired end reads. Reads were mapped to the *C. elegans* reference transcriptome from WormBase (version WS274) using STAR version 2.7.0. Duplicate reads were removed using SAMtools (version 1.9), and a counts matrix was generated using the featureCounts tool of SubReads (version 1.6.4).

### Comparing single cell and bulk gene expression profiles

Differential gene expression comparing sorted cell samples with sorted pan-neuronal samples was performed using TMM-normalized counts in edgeR (version 3.28.1). Two to five replicates per cell type were used in each sample (ASG: 4, AVE: 3, AVG: 3, AWA: 4, AWB: 5, DD: 3, PVD: 2, VD: 4, pan-neuronal: 5). Marker genes from the single cell dataset were selected using a Wilcoxon test in Seurat v3, calling enriched genes by comparing individual neuronal clusters to all other neuronal clusters. Marker genes were defined as genes with a log fold change >2, and adjusted p-value < 0.001. To examine marker gene enrichment in each bulk cell type, pairwise Wilcoxon tests were performed in R comparing the corresponding bulk cell type’s enrichment against the enrichment in all other bulk cell types.

To compare the overlap of gene detection between bulk and single cell datasets, bulk TMM counts were normalized to gene length, and the true positive rate (TPR) for detecting ground truth markers (see **Thresholding**) was calculated for a range of length normalized TMM values. At each expression threshold, if > 65% of samples showed expression equal to or higher than the threshold, the gene was called expressed. TPR, FPR, and FDR rates were calculated with 5,000 stratified bootstraps of the ground truth genes, which were generated using the R package boot (Canty and Ripley, 2019; Davison and Hinkley, 1997). We used a threshold of 5.7 length normalized TMM, to match the TPR (0.81) of the single cell Threshold 2. To calculate the relationship between single cell cluster size and the overlap between bulk and single cell gene expression, only protein coding genes were considered. Classifications from WormBase were used to define each gene’s RNA class.

### Alternative Splicing

Alternative splicing events were detected using the software SplAdder (Kahles et al., 2016). The common splicing graph was built based on all 32 individual samples and each pair of neurons was tested for differential use of AS events (with confidence level of 3 and parameters --ignore-mismatches, --validate-sg and sg_min_edge_count=3). The resulting tables were loaded in R to adjust the p-value for multiple testing, and events with FDR > 0.1 were discarded. Sashimi plots for the genes *mca-3* and *mbk-2* were generated using the Integrated Genomics Viewer (Robinson et al., 2011).

For the putative novel exons, the splicing graph generated by SplAdder was recovered. It consisted of 197,576 exons; of these, 3,860 were not annotated in WormBase WS274. To avoid counting exons resulting from intron retention events or imprecise annotation of neighboring exons, we filtered out exons sharing their start and end positions with annotated exons, to keep 2,142 exons displaying an unannotated start or end. As many of these had extensive overlap with annotated exons, we further filtered the set to keep 63 exons, 42 of them displaying no overlap with annotated exons, and 21 exons having less than 90% of their sequence overlapping with annotated exons.

### Comparative Analyses

We obtained gene orthologs between *C. elegans* and *M. musculus* by downloading the ortholog table from the Ensembl BioMart (Ensembl Genes 101) on the basis of stable Ensembl gene IDs. To increase the confidence of downstream interspecies comparisons, only one-to-one orthologs were taken into account. For the gene orthology between *C. elegans* and *Hydra*, we derived the high-quality ortholog table by three steps. First, we got the *Hydra*’s orthologs as the best Blast hits from the Swiss-Prot database (Siebert et al. 2019). The unique suffix of each gene (e.g., CP4B1_MOUSE) was converted to a standard gene name (e.g., Cyp4b1) of the associated species (e.g., mouse) using the online ID mapping tool (https://www.uniprot.org/uploadlists/) in UniProt. Second, we included *Hydra* genes whose best hits were from the defined 59 species: 42 vertebrate species (BOVIN, MOUSE, HUMAN, CHICK, XENLA, RAT, DANRE, PONAB, XENTR, MACFA, ORYLA, TAKRU, PANTR, PONPY, ICTPU, CRIGR, CANLF, PIG, SHEEP, SAIBB, MACMU, AILME, COTJA, LOPSP, RABIT, PHOSU, SALSA, CAVPO, MESAU, CHLAE, FELCA, ATEGE, HORSE, COLLI, TETNG, ONCMY, TAEGU, SQUAC, PAPAN, MELGA, CARAU and TETFL), 14 Drosophila species (DROPS, DROPB, DROYA, DROVI, DROMO, DROME, DROFU, DROMA, DROGR, DROSI, DROSE, DROER, DROAN and DROWI), two Caenorhabditis species (CAEEL and CAEBR), and one Nematostella species (NEMVE). For vertebrates, the gene names were further transferred to the format of mouse gene symbols (i.e., only first letter as an upper case). Third, genes from the three major categories were mapped back to the *C. elegans* gene IDs using corresponding ortholog tables between worms and mice, flies, and sea anemones respectively from Ensembl. The transcriptomic homology between cell types of *C. elegans* and mouse major organs was uncovered by checking their transcriptome similarities. Specifically, in each species we derived the transcriptome profile of each type by averaging gene expression across cells within the given cell type. We next calculated the interspecies transcriptome correlation (Pearson correlation coefficient) between each pair of cell types, and organized these cell types in a network with the graphopt layout using the package “igraph” in R (Csardi and Nepusz, 2006). The genes used during this procedure were the one-to-one orthologs between worms and mice which were also among the 7,390 highly variable genes used in the neuron network analysis. To remove weak connections and weakly linked cell types in the network, we discarded the edges (correlations) and corresponding nodes (cell types) with a transcriptome correlation < 0.7 with any cell types (Figure 7).

To investigate the molecular signatures that robustly exist between neurons of *C. elegans* and cell types from the mouse nervous system, we used the consensus module detection implemented in the “WGCNA” package in R (Langfelder and Horvath, 2008). First, we located the best scaling power (8), which is the minimum power to render the networks scale-free in both *C. elegans* and mouse. Next, we constructed the consensus network spanning the two species using the function “blockwiseConsensusModules” in WGCNA (tag-value pairs: networkType = “signed”, power = 6, minModuleSize = 20, deepSplit = 2, pamRespectsDendro = FALSE, mergeCutHeight = 0.25, numericLabels = TRUE, minKMEtoStay = 0), resulting in two confident consensus modules across species and corresponding eigen gene patterns (Figure 7B, C). This construction procedure of consensus network was repeated for single-cell *Hydra* data, with the scaling power selected to be 6 to make the network scale-free in this context and other procedures being constant (Figure 7D). The genes used here were the orthologs between worms and Hydra which were also among the 7,500 highly variable genes. Since many genes in Hydra were not matched in the Swiss-Port database, we also included one-to-many orthologs between worms and Hydra to incorporate more gene information. The Hydra genes orthologous to one given worm gene were collapsed to a pseudogene by averaging their gene expression in each cell subtype.

### Generation of invariant membrane contact and chemical connectome datasets

We compiled membrane contact and chemical synapse matrices from published electron microscope reconstructions, N2U (Cook et al., 2019; White et al., 1986) and Adults 7 and 8 (Witvliet et al., 2020). Membrane contact data are available for N2U and Adult 8. Chemical synapse data was obtained from three adult animals (N2U, Adult 7 and Adult 8). These sources contain data for each individual neuron (e.g., for each of the six IL2 neurons). Data were summed across the individual neurons corresponding to each neuron type in the single-cell data (e.g., IL2DL, IL2DR, IL2VL, IL2VR were summed for the IL2_DV class, IL2L and IL2R were summed for IL2_LR). Only contacts and synapses present across all animals were retained to generate high confidence sets of invariant contacts and synapses.

### Regulatory patterns of neuronal transcriptomes

In order to identify distinct regulatory patterns for the transcriptome of each neuron, log-transformed expression values were converted to z-scores from the distribution of expression across all neurons for each gene. A high (low) z-score for a particular gene in a specific neuron type indicates an up-regulated (down-regulated) gene relative to the expression in other neurons. For motif discovery in promoters and 3’UTRs, gene z-scores were mapped to their isoform transcripts. Unique isoforms were maintained by applying a simple duplicate removal procedure, which guarantees that no pair of promoters and no pair of 3’UTRs will have a Blast local alignment with E-value < 10^-10^ (Elemento et al., 2007). For promoter sequences we considered sequences 1KB upstream of the transcriptional start site of each isoform, while for 3’UTRs we considered 1KB within the from the start of each annotated 3’UTR sequence (or 1KB downstream of the stop codon for transcripts without annotated 3’UTRs). To identify expression patterns of co-regulated transcripts, z-score values across all neuron types were clustered using hierarchical clustering with three different cut-offs (python/scipy fcluster implementation, cosine metric, criterion=’distance’, cophenetic threshold= 1.2, 1.25, 1.37). We chose these thresholds to provide clustering of the data ranging from coarse to fine (16, 48, and 76 transcript clusters). For individual neurons, transcripts were categorized into bins with high to low z-scores based on the distribution of all z-scores across transcripts and neuron types. Z-score bin intervals were defined considering the following percentiles of the overall distribution of z-scores: 2.5%, 5%, 10%, 20%, 80%, 90%, 95%, 97.5%. For each neuron type, the top bin included transcripts with z-scores above the 97.5^th^ percentile, the second to top included z-scores between the 95^th^ and 97.5^th^ percentile, etc. The bottom bin included transcripts with z-scores below the 2.5^th^ percentile, the second to bottom included z-scores between the 2.5^th^ and 5^th^ percentile, etc. To avoid poorly populated bins, any given category containing less than 350 transcripts was merged with the next closest bin towards the center of the distribution.

### Cis-regulatory element discovery

To systematically explore the regulatory effect of short DNA and RNA cis-regulatory elements, we utilized FIRE, a computational framework for de novo discovery of linear motifs in DNA and RNA whose presence or absence in a transcript’s promoter and 3’UTR regions is informative of regulatory patterns. We ran FIRE in discrete mode including transcript identifiers (Wormbase transcript IDs) along with either their z-score bin categories (for individual neurons) or transcript cluster IDs (for patterns of co-regulated genes). Over representation (yellow) and under representation (blue) patterns are shown for each discovered motif within each category (bin or cluster) of transcripts as well as mutual information (MI) values and z-scores associated with a randomization-based statistical test. All discovered motifs pass a three-fold jackknifing test more than 6 out of 10 times. Each time one-third of the transcripts was randomly removed and the statistical significance of the MI value of the motif was reassessed. For each of the 10 tests, the remaining two-thirds of the transcripts was shuffled 10,000 times and the motif was deemed significant if its MI was greater than all 10,000 MI scores from the randomized sets (Elemento et al., 2007). For every motif identified through FIRE, we defined the regulon for that motif as the collection of transcripts that harbored instances of the motif in their promoters (DNA motifs) or 3’UTRs (RNA motifs).

### Motif families

Motifs with similar nucleotide compositions and regulons were discovered across individual neurons and gene expression patterns. We sought to identify the extent of redundancy between individual motifs and group them into motif families based on their similarity. We included additional motifs in this analysis for known transcription factors (CIS-BP, JASPAR), RNA binding proteins (CISBP-RNA) and miRNA 6-mer seeds (5’ extremity of known miRNA sequences of C. elegans). To quantify the similarity between nucleotide compositions between motifs we applied TOMTOM (MEME version 5.0.5). For each motif, we used its IUPAC motif sequence to convert it into a MEME formatted motif (iupac2meme function) as input to TOMTOM and compared it against all other discovered and known motifs. We specified a minimum overlap of 5, and an E-value threshold of 10 to identify significant matches. To quantify the extent of overlap between two motif modules, we defined a similarity measure between a module A and B as 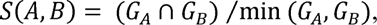 where *C_k_* is the set of transcripts in module *K*. We calculated TOMTOM and module similarity scores for all motif pairs. Module similarity scores were deemed significant if p<10^-4^ (hypergeometric test). To ensure that motifs are considered redundant only when they are similar both in nucleotide and module composition, we set the module similarity scores to 0 if either the TOMTOM or the module similarity scores were not significant. We clustered the motifs into motif families based on the masked similarity measures of all motif pairs using hierarchical clustering (python/scipy fcluster implementation, cosine metric, criterion=’distance’, cophenetic threshold= 0.9). We set out to identify potential known regulators that represent a given motif family. To this end, we applied TOMTOM to match the motif family members with the binding preferences of known regulators. For each motif family, we counted all the significant TOMTOM scores for every family member compared to a known regulator. We considered a known regulator as a potential match for the motif family, if it had a significant TOMTOM score for more than 2/3 of the family members.

### Associations of motif families and neuron types

We set out to assess the regulatory potential of each motif family on each neuron type. Motifs with positive regulatory potential should have consistent patterns across the z-score bins, i.e., predominantly over-represented in genes with high z-scores or under-represented in genes with low z-scores. On the other hand, motifs with negative regulatory potential should be over-represented in genes with low z-scores or under-represented in genes with high z-scores. For each neuron type and each motif, we considered the frequency of transcripts carrying the motif in the top two z-score bins combined (*f_t_*), as well as the bottom two z-score bins (*f_b_*). To consider a positive association of the motif with the neuron type we required that the motif is: *over-represented in the top two bins (p<0.005) and not over-represented in the bottom two bins (p>0.05)*, or, *under-represented in the bottom (p<0.005) two bins and not under-represented in the top two bins (p>0.05)*. To consider a negative association of the motif with the neuron type we required that the motif is: *over-represented in the bottom two bins (p<0.005) and not over-represented in the top two bins (p>0.05)*, or, *under-represented in the top two bins (p<0.005) and not under-represented in the bottom two bins (p>0.05)*. We calculated a Log_2_-fold ratio 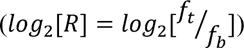 and an associated p-value (hypergeometric test) between the two categories. We reported significant associations 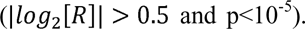. For each motif for the family, we report the Log_2_-fold ratio and signed p-value 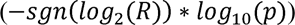 for the motif member with the lowest p-value.

### Differential gene expression analysis of transcripts encoding Cell Adhesion Molecules (CAMs)

Given a set of gene expression profiles for the neurons classes in the nerve ring and their memberships in different strata, we can execute standard differential gene expression (DGE) analysis (Soneson and Delorenzi, 2013) to determine which genes are enriched in members of particular strata. Standard DGE analysis involves performing univariate t-tests between the gene expression levels of members of a particular stratum versus the members of all the remaining strata. The visual representation of this test can be seen in Figure S17A. In detail, the DGE model involves fitting a regression model where the response variables are the gene expression levels for every neuron and the design matrix is a vector of 1s and −1s corresponding to the neurons in the two groups that are being compared. The gene expression is logarithm transformed to Gaussianize count-based data (Love et al., 2014). The output of this test is a vector of t-statistics and log-fold changes for every single gene in which this tuple of information can be visualized via volcano plots (Figure S16C). We deem that genes that pass the Bonferroni threshold for multiple comparisons (q<0.05) are significantly enriched or depleted in particular strata.

### Network differential gene expression analysis

Whereas standard DGE analysis is useful for delineating univariate differences between groups of neurons, here we introduce a generalization of DGE, termed “network” DGE (nDGE), to establish the genetic determinants of synaptic formation and maintenance. Unlike DGE where gene expression levels of disjoint groups of neurons are compared, in nDGE, **the multiplicative co-expression of genes**, between sets of **pairs** of neurons (representing edges in a network) is compared. The visual representation of the nDGE statistical model can be seen in Figure S19B. In nDGE, the response variables are the pairwise co-expression of all genes in all pairs of neurons. On the other hand, the design matrix captures two sets of pairs of neurons, one for each group. Similar to standard DGE, the output of this test is a set of t-statistics and log-fold changes for gene associations. However, unlike standard DGE, the t-statistics and log-fold changes in nDGE capture the effect of **co-expression of pairs of genes**, one corresponding to the gene observed in the pre-synaptic neuron partner and the other corresponding to the gene observed in the post-synaptic one. To deem a pair of genes significant under nDGE analysis, we also utilize the Bonferroni correction for p-values. However, the number of comparisons in nDGE is the square of the number of genes interrogated.

Since nDGE is a generalization of standard DGE, it enables the testing of a variety of hypotheses in addition to what is testable in standard DGE. The types of hypotheses that are tested are encoded in the design matrix of nDGE of which several examples are displayed in Figure S19C, D, E, F. Figure S19C shows how standard DGE can be executed through nDGE, by placing 1s and −1s in the diagonal of the design matrix corresponding to the neuron groups. Three other types of hypotheses that can be tested are whether particular gene pairs have global effects of synaptic formation across all the neurons (Figure S19D), whether there are differential gene co-expression differences in the synapses of two different neurons (Figure S19E), or which gene co-expression patterns are implicated in the synapses of an individual neuron (Figure S19F). In these scenarios, the design matrix has 1s where there is a synapse and a −1 where there is membrane contact, but no synapse, restricted to the sets of neurons of interest (all, pair, or one, respectively).

The main caveat in nDGE is the lack of independence of samples that are compared between groups. Since “samples” in nDGE are the co-expression of genes in pairs of neurons, the information from a particular neuron will inevitably be represented multiple times and possibly in different groups e.g., the gene expression from neuron AIA is represented in multiple synaptic gene co-expression values for all synaptic partners of AIA as well as the non-synaptic adjacent partners of AIA (Figure 9B). This lack of independence in the test samples can falsely inflate/deflate the sample variance, which can introduce excess false positives and false negatives. To accurately estimate the null distribution of the nDGE test statistics, we generate randomized “pseudoconnectomes” that respect the topology of the original connectome. Specifically, the pseudoconnectomes preserve the same number of synaptic partners for each neuron and the shuffled synaptic partners are confined to be neurons that have membrane contact (Milo et al., 2003). The latter constraint prevents infeasible pseudoconnectomes where synapses exist between neurons that do not share a membrane contact. Examples of pseudoconnectomes that are generated using the chemical connectome and membrane contact adjacency matrices are displayed in (Figure S19F). We execute nDGE analysis with the design matrices corresponding to 1000 pseudoconnectomes and compute a t-statistic using the mean and variance of the resulting null distribution.

While the nDGE technique introduced here is a generalization of standard DGE, interrogating the contribution of pairs of genes in the formation and maintenance of synapses between pairs of neurons, nDGE can only account for a single co-expressed gene in either of the two synaptic terminals (pre/post). For this reason, the nDGE model will tend to underestimate the effects of trimer (or higher-order) proteins in the formation and maintenance of synapses. Therefore, it is imperative to keep in mind that lack of significant hits for a particular neuron might not mean that there are no genes implicated in the formation of synapses for that neuron, but rather that higher-order gene interactions might be at play. Conceptually, it is straightforward to extend the model to higher-order gene interactions, but the prohibitive number of combinatorial gene co-expression enumeration is a computational bottleneck.

Another feature of nDGE is that it is a mass-univariate method, which does not take into account the possibility of interaction of different co-expressed genes in forming or inhibiting synapses. Therefore, the significance results output by nDGE tends to be very conservative with strict control of type 1 errors. This is in contrast with multivariate methods for explaining the genetic bases of connectivity (Kovacs et al., 2020). Due to the relatively high dimensionality of the gene expression data compared to the number of synapses in the chemical connectome, multivariate models tend to overfit and introduce type 1 errors.

### Supplemental Information titles and legends

**Figure S1. Isolating L4 larval stage neurons from fluorescent marker strains for single-cell RNA-Sequencing (scRNA-Seq)**. (**A**) Confocal images of the pan-neural marker, *rab-3::TagRFP* (*otIs355)*, ventral cord cholinergic motor neurons marked with *acr-2::GFP (juIs14)* and neurons in the head region dual-labeled with the GABA neuron-specific reporter, *unc-47::GFP (oxIs12)* and the pan neural marker strain, *rab-3::TagRFP*. Scale bars = 10 µm. L4 animals were treated with SDS-DTT and dissociated with pronase to produce single-cell suspensions. Targeted subgroups of neurons were isolated by Fluorescence Activated Cell Sorting (FACS, red boxes) and collected for scRNA-Seq using the 10x Genomics 3’ platform. B) Results from 17 separate profiling experiments were submitted to a series of processing steps to produce a final merged data set (see Methods). Right panel shows UMAP projection of all 100,955 cells, colored by cluster. C) Graphical depiction of relative abundance of each neuron class in cells isolated from the pan-neural marker strain (*rab-3::TagRFP*). The fraction of observed cells for each neuron type was divided by the expected ratio (i.e., # neuron type/302) and annotated for each cell type according to heat map index. Note under-representation (dark blue/black) of neurons in the anterior pharynx, anterior ganglion and pre-anal ganglion (dashed lines). * For pairs ASE and AWC, the individual neurons are treated separately. Related to Figure 1, 2.

**Figure S2. Distinguishing non-neuronal vs neuronal cells.** A) UMAP projection of 100,955 single cell profiles, colored by cluster. B) Known markers identify clusters of non-neuronal cells (*unc-122* + *galt-1*, coelomocytes; *myo-3* + *pat-10*, body wall muscles; *sqt-3* and *col-12*, epidermis). Boxes correspond to sub-regions in A. C) The neuropeptide processing gene *sbt-1* is expressed in all neuronal clusters (blue-magenta) and largely absent from non-neuronal clusters (gray). D) Sub-UMAP of all non-neuronal cells, labeled by cell type. E) *nspc-1*, a member of the nematode-specific peptide c (*nspc*) gene family, is restricted to a single non-neuronal cluster (arrow). F) The transcriptional reporter *nspc-1::GFP* is exclusively expressed in the excretory gland cell (yellow). DIC (Differential Interference Contrast). Scale bar = 10 µm. Related to Figure 1.

**Figure S3. Annotation of neuronal clusters.** A) Co-expression of the glutamate receptor genes *glr-3* and *glr-6* demarcate the RIA cluster. B) Co-expression of the homeodomain transcription factors *ceh-10* and *lim-4* label the RID cluster. C) Co-expression of the transient receptor potential channel (trp) gene *ocr-2* and tryptophan hydroxylase *tph-1* label the sensory neuron ADF. D) The nuclear hormone receptor *nhr-236* is primarily detected in three clusters, corresponding to RIP, M2, and MC. E) Z-projection of confocal stack showing *nhr-236::NLS-GFP* expression in RIP and MC in a NeuroPAL strain. M2 (not shown) also consistently expressed GFP. Scale bar = 5 µm. Related to Figure 2.

**Figure S4. Identifying pharyngeal neuron types and neuron clustering.** A) Diagram denoting neurons (blue) in pharynx (gray). B) Sub-UMAP of neurons expressing the pharyngeal neuron marker *ceh-34* revealed independent clusters for each known pharyngeal neuron type. C-E) Expression of known specific marker genes identifies individual pharyngeal neuron types. C) The M4-specific homeodomain transcription factor, *ceh-28* is exclusively detected in a small but distinct group of 12 cells. D) *ceh-2*, a marker for I3, M3 and NSM pharyngeal neurons, is restricted to 3 clusters. E) Restricted expression of *gur-3*, a marker for I2 and I4 pharyngeal neurons. F) Network analysis of all transcriptionally distinct neuron types and subtypes. *C. elegans* neuron types organized in a force-directed network according to transcriptomic similarities. Colors denote distinct neuron modalities, and widths of edges show the strengths of the transcriptome similarity between each pair of neuron types. Edges with Pearson correlation coefficients > 0.7 are shown. Related to Figure 2.

**Figure S5. Identification of neuron sub-types.** A) Sub-UMAPs showing expression of known markers for neuron sub-types for ASE (*gcy-3*, ASER and *gcy-6*, ASEL), AWC (*str-2*, AWC^ON^ and *srsx-3*, AWC^OFF^), VC (*vab-7 + unc-4*, VC1-3,6 and *cat-1 + unc-4*, VC4-5), RMD (*unc-42 + unc-46*, RMD DV and *unc-42 + cog-1*, RMD LR), RME (*unc-25 + lim-6*, RME LR and *unc-25 + slt-1*, RME DV), DA9 (*unc-53 + egl-5*), VA12 (*bnc-1 + egl-5*), DB (*vab-7*, DB1-7 and *vab-7 + sptf-1*, DB1). B) Volcano plots of genes that are differentially expressed between neuron subtypes. Inset depicts color coding for selected gene families. Related to Figure 3.

**Figure S6. Establishing expression thresholds.** A) Jitter plot of normalized *ttx-1* expression (Y-axis) in all neuronal clusters (X-axis) shows strongest expression in AFD, M2, RIB and RIP. Confocal image of *ttx-1^crispr^::GFP* shows expression in AFD, M2, RIB and RIP neurons and in glia and epidermal cells. C) Receiver-Operator Characteristic (ROC) curve of True Positive Rate (TPR) vs False Positive Rate (FPR) for a range of thresholds (1-4) (red dots) compared to ground truth expression data (see Methods). Increased stringency diminishes both the TPR and FPR. Grey shading represents 95% confidence intervals. D) Thresholds 1-4 (red dots) plotted on Precision-Recall (PR) Curve of Recall (TPR) vs Precision [1 – False Discovery Rate (FDR)]. Grey shading represents 95% confidence intervals. E) The proportion of cells in each neuron-specific cluster expressing *ttx-1*. Inset shows expanded view of boxed region. Thresholds (1-4) are set to different proportions of *ttx-1*-expressing cells in each cluster (see Methods). Note that neurons RIB and M2, that show expression of native *ttx-1^crispr^::GFP,* are excluded by the thresholds 3 and 4. Numbers (1-4) correspond to thresholds in C and D. F) Jitter plot shows strong expression of *unc-25*/GAD in seven known GABAergic neuron types (AVL, DVB, RIB, RIS, RME, DD, VD) but scattered *unc-25/GAD* expression is also detected in other cell types. G) Bar graph plotting the proportion of cells in each neuron type that express *unc-25/GAD*. Thresholds of increasing stringency (1-4) are set to different proportions of cells in a given cluster that express *unc-25/GAD*. Note that threshold 2 distinguishes known *unc-25/GAD*-positive neurons from other neuron types with lower detected levels of *unc-25/GAD* (see Methods). H) Cell soma for each neuron type in the head, mid-body and tail regions are colored according to the number of genes detected using threshold 2. Neuron types in the anterior ganglion (dashed line) are among those with the fewest cells and also lower numbers of detected genes. I) Number of genes detected with threshold 2 for each neuron type plotted against the number of cells in each neuron-type cluster. Spearman’s rank correlation = 0.783, p < 2.2e-16). J) Number of genes detected with threshold 2 plotted against the True Positive Rate (TPR) for each neuron type. Spearman’s rank correlation = 0.678, p < 2.2e-16. Neurons with fewest cells, low TPR and number of genes/type are denoted (red). Related to Figure 4, 5, S9-S12.

**Figure S7. Expression of neurotransmitter-related genes.** Heatmap showing scaled expression within each neuron type for genes specifically required for mediating synthesis and release of different classes of chemical neurotransmitters.

**Figure S8. Neuropeptide gene reporter strains validate single cell RNA-Seq data**. A) Diagrams depicting upstream regions (black lines) in transcriptional GFP reporter genes. B) Heatmap showing single cell RNA-Seq expression of four selected neuropeptides (*nlp-42*, *nlp-17*, *nlp-52*, *flp-33*) and (C) corresponding confocal micrographs. Neuron types expressing each neuropeptide reporter were determined by co-localization with NeuroPAL markers (not shown). *nlp-42p::gfp* is robustly expressed in AIN and RMF with faint expression in NSM and PHB. *nlp-17* expression at threshold 2 was 480x stronger in PVQ than for six additional neuron types and the *nlp-17p::gfp* reporter was selectively detected in PVQ. GFP reporters for *nlp-52* and *flp-33* (C) are largely congruent with single-cell RNA-Seq results (B). Related to Figure 4.

**Figure S9. Expression of selected gene families for neuronal signaling and ion channel function in different neuron classes.** A) Bar graph showing expression of neuropeptide encoding genes at threshold 2 for neurons grouped by modality (Sensory, Interneuron, Motor, Pharyngeal). B) Number of genes (threshold 2) expressed in each neuron type grouped by modality for all genes and neuropeptide signaling genes (neuropeptide-encoding genes, neuropeptide receptors, peptide degradation genes, *irld* genes). Boxes are interquartile ranges. Statistical tests: ANOVA, with Tukey post-hoc comparisons for neuropeptide receptors, Kruskal-Wallis test for other genes. *p < 0.05, **p < 0.01, ***p < 0.001. C) Heatmaps showing expression of genes for neurosecretory components, TWK K^+^ channels, and Bestrophin anion channels across 128 neuron types (columns) grouped by neuron modality. Protein subfamilies are clustered separately. D) Number of genes (threshold 2) expressed in each neuron type grouped by modality Ligand-Gated Ion Channels (LGICs) and TWK K^+^ Channels. Boxes are interquartile ranges. Statistical tests: Kruskal-Wallis. *p < 0.05, **p < 0.01, ***p < 0.001, ****p < 0.0001. Related to Figure 4.

**Figure S10. Expression of ligand-gated ion channels.** A) Heatmaps of ionotropic receptors including 89 Cys-loop ligand-gated ion channel (LGIC) genes grouped by neurotransmitter (Acetylcholine, GABA, Amine, Glutamate) and by ion selectivity (Cl^-^ vs cation) and 14 glutamate-gated cation channels across 128 neuron types (columns). B) (Top) Inset of selected region of heatmap in A showing 18 Cys-loop LGICs enriched in pharyngeal neurons (bottom). C) Cumulative distribution plot of number of neuron types expressing genes encoding ionotropic receptors (Cys-loop LGICs and glutamate-gated cation channels), DEG/ENaC channels, potassium (K) channels, chloride channels, VGCC (Voltage-Gated Calcium Channels) and ribosomal proteins (see also Figure 4A). Related to Figure 4.

**Figure S11. Expression of chemoreceptor GPCRs and downstream signaling pathways.** A) Bar graph showing expression of chemoreceptor GPCRs across all neuron types. All neurons express at least two chemoreceptor GPCRs. Note enrichment in a subset of ciliated amphid/phasmid sensory neurons. B) Heatmaps showing expression of genes for G-alpha, arrestin and RGS families. Note that neurons expressing the highest numbers of chemoreceptors (Panel A, ADL, ASH, ASJ, ASK, AWA, PHA, PHB) also abundantly express G-alpha and RGS proteins. C) The number of chemoreceptor GPCRs (left) and G-alpha genes (right) grouped by neuron modality (Sensory, Interneuron, Motor, Pharyngeal). Boxes are interquartile ranges. Kruskal-wallis test, *p < 0.05, **p < 0.01, ***p < 0.001, ****p < 0.0001. D) Cumulative distribution plot of chemoreceptor GPCRs, G-alpha, arrestin and RGS families.

**Figure S12. Expression of transcription factor subfamilies.** Heatmaps showing expression of the T-box, AT hook, bZIP and C2H2 zinc-finger (ZF) transcription factor families. Neurons are grouped according to modality (Sensory, Interneuron, Motor, Pharyngeal).and genes are clustered within each family. Related to Figure 5.

**Figure S13. Detection of neuron-specific splicing of a novel exon by bulk RNA-Seq.** Sashimi plot representing the splicing pattern of the gene *mbk-2* in neurons DD, AWA and pan-neuronal samples. Two representative samples are represented for each condition. The (*) indicates an alternative 5’ exon used exclusively in AWA and undetectable in pan-neuronal samples. Related to Figure 6.

**Figure S14. FIRE discovered known regulatory motifs in individual neurons.** A) Heatmap of scaled gene expression (z-scores for each gene in each neuron). Both genes (columns) and neurons (rows) are ordered by similarity. X-axis color bar (top) denotes 48 gene clusters. Row color bar (left) indicates neuron modality (Sensory, Motor, Interneuron, Pharyngeal). B) Selected results from FIRE analysis of 48 gene clusters. C) Left: Motif family 109 includes highly similar motifs discovered in independent FIRE analyses from individual neurons. Center: Motifs from the CIS-BP database for six different K50 homeodomain transcription factors (CEH-36, CEH-37, TTX-1, CEH-53, UNC-30, CEH-45) match motif family 109. Right) Expression (TPM or transcripts per million) of two transcription factors (*ceh-36, ceh-37*) in our scRNA-Seq data corresponds to the neurons in which the cognate motif was discovered. D) The TAATCC motif (red boxes) in the 5’ regions of genes enriched in ASEL, ASER, AWC^ON^ or AWC^OFF^, including genes expressed in all four neurons (top rows) as well as genes expressed in subsets of these neurons. Expression of *eat-4*, *tax-2*, *gcy-6*, *gcy-7* and *srt-2* is *ceh-36*-dependent in these neurons. E) Schematic showing the z-score bins tested for relative enrichment for motif-neuron associations (See Methods). F) Motif-neuron associations (from FIRE Main Figure B for motif family 109 in a subset of neurons) showing correspondence to expression of K50 homeodomain proteins (black arrows). Colors indicate −log10(p-value) for positive associations and log10(p-value) for negative associations. Asterisks denote significant associations (pval < 1e-5, log fold change > 0.5). G) Top: Two similar motifs discovered in FIRE runs of individual neurons IL2_DV and ADL are members of motif family 184 that matches the DAF-19 motif from CIS-BP. Other members of the DAF-19 motif family were discovered in FIRE analysis of 8 additional individual neurons (black boxes). Bottom: Motif family 184 showed significant positive associations (asterisks) with 22 ciliated sensory neuron types (green boxes) and significant negative associations with some motor neurons (blue boxes with asterisks). Related to Figure 8.

**Figure S15. RNA motif families are positively associated with a majority of neuron types.** Heatmap of log-transformed p-values showing motif-neuron associations for RNA motif families (columns) across all neuron types (rows). Note the strong positive signal in sensory neurons, and the one motif family with largely negative associations with neurons (blue arrow). B) Representative RNA motif (from ADL FIRE run) of family 23 showing a poly-C RNA sequence in the 3’ UTR. Related to Figure 8.

**Figure S16. Differential expression of CAMs between strata.** A) Cartoon representation of nerve ring strata (Moyle et al., 2020). Figure courtesy of Daniel Colón-Ramos. B) Heatmap of CAM expression (columns) in neurons grouped by strata (rows). Black arrows indicate four significantly enriched CAMs with the highest log fold changes between strata. Colors indicate log-transformed expression values. Red lines separate strata. C) Volcano plots showing log-fold change (x-axis) by −log10 p-value (y-axis) for each CAM in the neurons in each stratum compared to all other neurons. The six labeled genes were significantly enriched in the strata shown. Related to Figure 9.

**Figure S17. Differential expression of cell adhesion molecules among neurons and their postsynaptic partners.** A) Left: Cartoon representation of the *C. elegans* nerve ring. Right: Subset of nerve ring neurites from the touch neurons AVM, ALM and their synaptic partner BDU. Courtesy of WormAtlas. B) Diagram of neurons receiving chemical postsynaptic output from AVM (right) and neurons with membrane contact but no synapses with AVM (left). Differential cell adhesion molecule (CAM) gene expression was determined between AVM + synaptic partners and AVM + non-synaptic adjacent neurons. C) Heatmap of the 20 gene pairs with highest log fold change in AVM and its postsynaptic outputs compared to AVM and adjacent neurons (right of vertical red line). The 20 gene pairs with highest log fold change in AVM and adjacent neurons compared to AVM and synaptic partners are shown to the left of vertical red line. Blue and dark red arrows indicate gene pairs common among analyses of AVM and ALM (panel D). The first gene listed is expressed in AVM, the second gene is expressed in the synaptic or adjacent partner. D) Correlation matrix showing the relationship of CAM usage across all neurons in the nerve ring (84 neuron types). Note the similarities in CAM usage among the neurons in the lower right-hand corner. The black arrows indicate AVM and ALM (correlation 0.607). E) Heatmap as in C, for ALM. 10 of the top 20 gene pairs (dark red arrows) higher in ALM and its synaptic partners compared to ALM and adjacent neurons are common with the AVM analysis in B. On the left, 7 of the top 20 genes pairs higher in ALM/adjacent are common with AVM (blue arrows). F) The membrane adjacency matrix was grouped according to nerve ring strata as defined in Moyle, et al. Within each stratum, neurons were ordered according to their CAM usage correlations (see panel H). Membrane contact is denser within strata than between strata, but not for the unassigned neurons which generally contact neurons in other strata. Colored bars indicate neuron strata assignments. Each stratum is outlined by a thin red box. G) The same ordering as in F was imposed upon the chemical connectome revealing that most synapses are detected between neurons within the same stratum. H) The CAM usage correlation matrix (as in D) was grouped by strata, then sorted by similarity (using multidimensional scaling) within each stratum. Neurons within stratum 1 are similar to each other. Strata 2, 3, and 4 all show two distinct groups of neurons based on CAM usage at postsynaptic outputs. Related to Figure 9.

**Figure S18. CAM usage across nerve ring neurons.** A) Heatmap of CAM usage in presynaptic connections for each neuron in the nerve ring. Neurons are rows, gene pairs are columns. Colors indicate the log of the t-statistic comparing gene pair expression in each neuron + presynaptic partners vs each neuron + non-synaptic adjacent neurons. Positive values (orange and red) indicate higher expression in synaptically-connected neurons relative to adjacent neurons. Negative values (blue) indicate higher expression in adjacent neurons relative to synaptically-connected neurons. Only gene pairs with a log fold change in expression > 0.2 in either direction are displayed. Black arrows indicate AIA and AIY, and red arrows indicate two gene pairs with the high log fold changes in synaptic neurons for AIA and AIY. B) Heatmap as in A, but for CAM usage in postsynaptic connections for each neuron in the nerve ring. Black arrows indicate AVM and ALM, red arrows indicate gene pairs with high log fold change in synaptic connections relative to adjacent neurons for both AVM and ALM, and blue arrows indicate gene pairs with high log fold change in adjacent neurons compared to synaptically connected neurons for both AVM and ALM. Related to Figure 9.

**Figure S19. Network differential expression methods.** A) Overview of standard differential gene expression analysis (DGE): DGE can be represented as a regression model with the response variables as the neuron-wise gene expression data and the design matrix as 1s and −1s denoting the group memberships of neurons. B) Schematic of network differential gene expression analysis (nDGE): nDGE is a generalization of DGE that represents the pairwise co-expression of all genes in all pairs of neurons as the response variable. The design matrix is a square matrix of all pairs of neurons with 1s and −1s placed in locations where pairs of edges of the network are contrasted. C) Design matrix for nDGE that yields identical results as standard DGE. D) Design matrix for nDGE that explores the global genetic differences between synapses and non-synaptic membrane contact. E) Design matrix for nDGE that explores the differential gene co-expression differences between the synapses of two different neurons. F) Design matrix for nDGE that aims to demonstrate gene co-expression enrichment in synapses of a particular neuron compared to non-synaptic membrane contacts. G) The statistical significance of nDGE test statistic is obtained by computing the mean and variance of the null distribution of the test statistic resulting from pseudoconnectomes that respect the network topology of the original connectome. Related to Figure 9.

**Table S1. Strains used in the study**

**Table S2. 10X Genomics 3’ single-cell experiment details**

**Table S3. Tissue and cell type markers used for annotation**

**Table S4. Number of detected cells per cell type by experiment**

**Table S5. Ground truth matrix for thresholding**

**Table S6. Neuron-type thresholding metrics**

**Table S7. Genes and gene families used in cumulative fraction plots**

**Table S8. Genes detected in single neuron types under Threshold 1**

**Table S9. Genes detected in single neuron types under Threshold 2**

**Table S10. Genes detected in single neuron types under Threshold 3**

**Table S11. Genes detected in single neuron types under Threshold 4**

**Table S12. Differential use of splicing sites between 8 neuron classes**

**Table S13. Putative novel exons**

**Table S14. C. elegans-mouse consensus gene module C1**

**Table S15. C. elegans-mouse consensus gene module C2**

**Table S16. C. elegans-hydra consensus gene module C1**

**Table S17. C. elegans-hydra consensus gene module C2**

**Table S18. C. elegans-hydra consensus gene module C3**

**Table S19. Invariant membrane contact matrix for nerve ring**

**Table S20. Invariant chemical synapse matrix for nerve ring**

## References

Adorjan, I., Tyler, T., Bhaduri, A., Demharter, S., Finszter, C.K., Bako, M., Sebok, O.M., Nowakowski, T.J., Khodosevich, K., Møllgård, K., et al. (2019). Neuroserpin expression during human brain development and in adult brain revealed by immunohistochemistry and single cell RNA sequencing. J. Anat. 235, 543–554.

Albertson, D.G., and Thomson, J.N. (1976). The pharynx of Caenorhabditis elegans. Philos. Trans. R. Soc. Lond. B. Biol. Sci. 275, 299–325.

Allen, A.M., Neville, M.C., Birtles, S., Croset, V., Treiber, C.D., Waddell, S., Goodwin, S.F., and Mann, R.S. (2020). A single-cell transcriptomic atlas of the adult Drosophila ventral nerve cord. Elife 9, e54074.

Arzalluz-Luqueángeles, and Conesa, A. (2018). Single-cell RNAseq for the study of isoforms-how is that possible? Genome Biol. 110.

Barabási, D.L., and Barabási, A.L. (2020). A Genetic Model of the Connectome. Neuron 105, 435–445.

Bargmann, C.I., and Horvitz, H.R. (1991). Control of larval development by chemosensory neurons in Caenorhabditis elegans. Science 251, 1243–1246.

Baruch, L., Itzkovitz, S., Golan-Mashiach, M., Shapiro, E., and Segal, E. (2008). Using expression profiles of Caenorhabditis elegans neurons to identify genes that mediate synaptic connectivity. PLoS Comput. Biol. 4, e1000120.

Becht, E., McInnes, L., Healy, J., Dutertre, C.A., Kwok, I.W.H., Ng, L.G., Ginhoux, F., and Newell, E.W. (2019). Dimensionality reduction for visualizing single-cell data using UMAP. Nat. Biotechnol. 37, 38–44.

Beg, A.A., Ernstrom, G.G., Nix, P., Davis, M.W., and Jorgensen, E.M. (2008). Protons Act as a Transmitter for Muscle Contraction in C. elegans. Cell 132, 149–160.

Bhatla, N., and Horvitz, H.R. (2015). Light and Hydrogen Peroxide Inhibit C.elegans Feeding through Gustatory Receptor Orthologs and Pharyngeal Neurons. Neuron 85, 804–818.

Bhatla, N., Droste, R., Sando, S.R., Huang, A., and Horvitz, H.R. (2015). Distinct neural circuits control rhythm inhibition and spitting by the myogenic pharynx of C. elegans. Curr. Biol. 25, 2075–2089.

Bhattacharya, A., Aghayeva, U., Berghoff, E.G., and Hobert, O. (2019). Plasticity of the Electrical Connectome of C. elegans. Cell 176, 1174–1189.

Blondel, V.D., Guillaume, J.L., Lambiotte, R., and Lefebvre, E. (2008). Fast unfolding of communities in large networks. J. Stat. Mech. Theory Exp. 83, 036103.

Brenner, S. (1974). The genetics of Caenorhabditis elegans. Genetics 77, 71–94.

Brittin, C.A., Cook, S.J., Hall, D.H., Emmons, S.W., and Cohen, N. (2020). Beyond the connectome: A map of a brain architecture derived from whole-brain volumetric reconstructions. BioRxiv https://doi.org/10.1101/2020.05.24.112870.

Brockie, P.J., Madsen, D.M., Zheng, Y., Mellem, J., and Maricq, A. V. (2001). Differential expression of glutamate receptor subunits in the nervous system of Caenorhabditis elegans and their regulation by the homeodomain protein UNC-42. J. Neurosci. 21, 1510–1522.

Bruce, F.M., Brown, S., Smith, J.N., Fuerst, P.G., and Erskine, L. (2017). DSCAM promotes axon fasciculation and growth in the developing optic pathway. Proc. Natl. Acad. Sci. U. S. A. 114, 1702–1707.

Canty, A., and Ripley, B. (2019). boot: Bootstrap R (S-Plus) Functions. R package version 1.3–24.

Cao, J., Packer, J.S., Ramani, V., Cusanovich, D.A., Huynh, C., Daza, R., Qiu, X., Lee, C., Furlan, S.N., Steemers, F.J., et al. (2017). Comprehensive single-cell transcriptional profiling of a multicellular organism. Science 357, 661–667.

Cao, J., Spielmann, M., Qiu, X., Huang, X., Ibrahim, D.M., Hill, A.J., Zhang, F., Mundlos, S., Christiansen, L., Steemers, F.J., et al. (2019). The single-cell transcriptional landscape of mammalian organogenesis. Nature 566, 496–502.

Chang, S., Johnston, R.J., and Hobert, O. (2003). A transcriptional regulatory cascade that controls left/right asymmetry in chemosensory neurons of c. elegans. Genes Dev. 17, 2123– 2137.

Chase, D.L., and Koelle, M.R. (2007). Biogenic amine neurotransmitters in C. elegans. WormBook 1–15.

Chen, C., and Jonas, P. (2017). Synaptotagmins: That’s Why So Many. Neuron 694–696.

Chen, Z., Hendricks, M., Cornils, A., Maier, W., Alcedo, J., and Zhang, Y. (2013). Two Insulin-like Peptides Antagonistically Regulate Aversive Olfactory Learning in C. elegans. Neuron 77, 572–585.

Choi, H.S., Hwang, C.K., Song, K.Y., Law, P.Y., Wei, L.N., and Loh, H.H. (2009). Poly(C)-binding proteins as transcriptional regulators of gene expression. Biochem. Biophys. Res. Commun. 380, 431–436.

Cohen, E., Chatzigeorgiou, M., Husson, S.J., Steuer-Costa, W., Gottschalk, A., Schafer, W.R., and Treinin, M. (2014). Caenorhabditis elegans nicotinic acetylcholine receptors are required for nociception. Mol. Cell. Neurosci. 59, 85–96.

Colón-Ramos, D.A., Margeta, M.A., and Shen, K. (2007). Glia promote local synaptogenesis through UNC-6 (netrin) signaling in C. elegans. Science 318, 103–106.

Colosimo, M.E., Tran, S., and Sengupta, P. (2003). The Divergent Orphan Nuclear Receptor ODR-7 Regulates Olfactory Neuron Gene Expression via Multiple Mechanisms in Caenorhabditis elegans. Genetics 165, 1779–1791.

Cook, S.J., Jarrell, T.A., Brittin, C.A., Wang, Y., Bloniarz, A.E., Yakovlev, M.A., Nguyen, K.C.Q., Tang, L.T.H., Bayer, E.A., Duerr, J.S., et al. (2019). Whole-animal connectomes of both Caenorhabditis elegans sexes. Nature 571, 63–71.

Cornils, A., Gloeck, M., Chen, Z., Zhang, Y., and Alcedo, J. (2011). Specific insulin-like peptides encode sensory information to regulate distinct developmental processes. Development 138, 1183–1193.

Cox, E.A., Tuskey, C., and Hardin, J. (2004). Cell adhesion receptors in C. elegans. J. Cell Sci. 117, 1867–1870.

Croset, V., Rytz, R., Cummins, S.F., Budd, A., Brawand, D., Kaessmann, H., Gibson, T.J., and Benton, R. (2010). Ancient protostome origin of chemosensory ionotropic glutamate receptors and the evolution of insect taste and olfaction. PLoS Genet. 6, e1001064.

Csardi, G., and Nepusz, T. (2006). The igraph software package for complex network research. InterJournal Complex Syst.

Davison, A.C., and Hinkley, D. V. (1997). Bootstrap Methods and their Application (Cambridge: Cambridge University Press).

Dehghannasiri, R., Olivieri, J.E., and Salzman, J. (2020). Specific splice junction detection in single cells with SICILIAN. BioRxiv https://doi.org/10.1101/2020.04.14.041905.

Dlakic, M. (2002). A new family of putative insulin receptor-like proteins in C. elegans. Curr. Biol. 12, R155–7.

Driever, W., and Nüsslein-Volhard, C. (1989). The bicoid protein is a positive regulator of hunchback transcription in the early Drosophila embryo. Nature 337, 138–143.

Efimenko, E., Bubb, K., Mark, H.Y., Holzman, T., Leroux, M.R., Ruvkun, G., Thomas, J.H., and Swoboda, P. (2005). Analysis of xbx genes in C. elegans. Development 132, 1923–1934.

Elemento, O., Slonim, N., and Tavazoie, S. (2007). A Universal Framework for Regulatory Element Discovery across All Genomes and Data Types. Mol. Cell 28, 337–350.

Fox, R.M., Von Stetina, S.E., Barlow, S.J., Shaffer, C., Olszewski, K.L., Moore, J.H., Dupuy, D., Vidal, M., and Miller, D.M. (2005). A gene expression fingerprint of C. elegans embryonic motor neurons. BMC Genomics 6, 42.

Fuxman Bass, J.I., Pons, C., Kozlowski, L., Reece-Hoyes, J.S., Shrestha, S., Holdorf, A.D., Mori, A., Myers, C.L., and Walhout, A.J. (2016). A gene-centered C. elegans protein– DNA interaction network provides a framework for functional predictions. Mol. Syst. Biol. 12, 884.

Gendrel, M., Atlas, E.G., and Hobert, O. (2016). A cellular and regulatory map of the GABAergic nervous system of C. elegans. Elife 5, e17686.

Granato, M., Schnabel, H., and Schnabel, R. (1994). pha-1, a selectable marker for gene transfer in C.elegans. Nucleic Acids Res. 22, 1762–1763.

Grove, C.A., De Masi, F., Barrasa, M.I., Newburger, D.E., Alkema, M.J., Bulyk, M.L., and Walhout, A.J.M. (2009). A Multiparameter Network Reveals Extensive Divergence between C. elegans bHLH Transcription Factors. Cell 138, 314–327.

Hafemeister, C., and Satija, R. (2019). Normalization and variance stabilization of single-cell RNA-seq data using regularized negative binomial regression. Genome Biol. 20, 296.

Haghverdi, L., Lun, A.T.L., Morgan, M.D., and Marioni, J.C. (2018). Batch effects in single-cell RNA-sequencing data are corrected by matching mutual nearest neighbors. Nat. Biotechnol. 36, 421–427.

Hammarlund, M., Hobert, O., Miller, D.M., and Sestan, N. (2018). The CeNGEN Project: The Complete Gene Expression Map of an Entire Nervous System. Neuron 99, 430–433.

Han, X., Wang, R., Zhou, Y., Fei, L., Sun, H., Lai, S., Saadatpour, A., Zhou, Z., Chen, H., Ye, F., et al. (2018). Mapping the Mouse Cell Atlas by Microwell-Seq. Cell 172, 1091–1107.

Harris, T.W., Arnaboldi, V., Cain, S., Chan, J., Chen, W.J., Cho, J., Davis, P., Gao, S., Grove, C.A., Kishore, R., et al. (2020). WormBase: A modern Model Organism Information Resource. Nucleic Acids Res. 48, D762–D767.

He, S., Philbrook, A., McWhirter, R., Gabel, C. V., Taub, D.G., Carter, M.H., Hanna, I.M., Francis, M.M., and Miller, D.M. (2015). Transcriptional control of synaptic remodeling through regulated expression of an immunoglobulin superfamily protein. Curr. Biol. 25, 2541–2548.

Hobert, O. (2002). PCR fusion-based approach to create reporter Gene constructs for expression analysis in transgenic C. elegans. Biotechniques 32, 728–730.

Hobert, O. (2013). The neuronal genome of Caenorhabditis elegans. WormBook 1–106.

Hobert, O., Glenwinkel, L., and White, J. (2016). Revisiting Neuronal Cell Type Classification in Caenorhabditis elegans. Curr. Biol. 26, R1197–R1203.

Hu, P.J. (2007). Dauer. WormBook 1–19.

Husson, S.J., Lindemans, M., Janssen, T., and Schoofs, L. (2009). Comparison of Caenorhabditis elegans NLP peptides with arthropod neuropeptides. Trends Parasitol. 25, 171–181.

Iwasa, H., Yu, S., Xue, J., and Driscoll, M. (2010). Novel EGF pathway regulators modulate C. Elegans healthspan and lifespan via EGF receptor, PLC-γ, and IP3R activation. Aging Cell 9, 490–505.

Johnston, R.J., Chang, S., Etchberger, J.F., Ortiz, C.O., and Hobert, O. (2005). MicroRNAs acting in a double-negative feedback loop to control a neuronal cell fate decision. Proc. Natl. Acad. Sci. U. S. A. 102, 12449–12454.

Jones, A.K., and Sattelle, D.B. (2008). The cys-loop ligand-gated ion channel gene superfamily of the nematode, Caenorhabditis elegans. Invertebr. Neurosci. 8, 41–47.

Jones, A.K., Davis, P., Hodgkin, J., and Sattelle, D.B. (2007). The nicotinic acetylcholine receptor gene family of the nematode Caenorhabditis elegans: An update on nomenclature. Invertebr. Neurosci. 7, 129–131.

Kahles, A., Ong, C.S., Zhong, Y., and Rätsch, G. (2016). SplAdder: Identification, quantification and testing of alternative splicing events from RNA-Seq data. Bioinformatics 32, 1840–1847.

Kaletsky, R., Lakhina, V., Arey, R., Williams, A., Landis, J., Ashraf, J., and Murphy, C.T. (2016). The C. elegans adult neuronal IIS/FOXO transcriptome reveals adult phenotype regulators. Nature 529, 92–96.

Kato, S., Kaplan, H.S., Schrödel, T., Skora, S., Lindsay, T.H., Yemini, E., Lockery, S., and Zimmer, M. (2015). Global Brain Dynamics Embed the Motor Command Sequence of Caenorhabditis elegans. Cell 163, 656–669.

Kaufman, A., Dror, G., Meilijson, I., and Ruppin, E. (2006). Gene expression of Caenorhabditis elegans neurons carries information on their synaptic connectivity. PLoS Comput. Biol. 2, e167.

Khan, A., Fornes, O., Stigliani, A., Gheorghe, M., Castro-Mondragon, J.A., Van Der Lee, R., Bessy, A., Chèneby, J., Kulkarni, S.R., Tan, G., et al. (2018). JASPAR 2018: Update of the open-access database of transcription factor binding profiles and its web framework. Nucleic Acids Res. 46, D260–D266.

Kim, B., and Emmons, S.W. (2017). Multiple conserved cell adhesion protein interactions mediate neural wiring of a sensory circuit in C. elegans. Elife 6, e29257.

Kim, K., and Li, C. (2004). Expression and regulation of an FMRFamide-related neuropeptide gene family in Caenorhabditis elegans. J. Comp. Neurol. 475, 540–550.

Koga, M., and Ohshima, Y. (2004). The C. elegans ceh-36 Gene Encodes a Putative Homemodomain Transcription Factor Involved in Chemosensory Functions of ASE and AWC Neurons. J. Mol. Biol. 336, 579–587.

Korswagen, H.C., Park, J.H., Ohshima, Y., and Plasterk, R.H.A. (1997). An activating mutation in a Caenorhabditis elegans G(s) protein induces neural degeneration. Genes Dev. 11, 1493– 1503.

Kotera, I., Tran, N.A., Fu, D., Kim, J.H.J., Byrne Rodgers, J., and Ryu, W.S. (2016). Pan-neuronal screening in Caenorhabditis elegans reveals asymmetric dynamics of AWC neurons is critical for thermal avoidance behavior. Elife 5, e19021.

Kovacs, I.A., Barabási, D.L., and Barabási, A.L. (2020). Uncovering the genetic blueprint of the C. elegans nervous system. BioRxiv https://doi.org/10.1101/2020.05.04.076315.

Koziol, U., Koziol, M., Preza, M., Costábile, A., Brehm, K., and Castillo, E. (2016). De novo discovery of neuropeptides in the genomes of parasitic flatworms using a novel comparative approach. Int. J. Parasitol. 46, 709–721.

Lainé, V., Frøkjær-Jensen, C., Couchoux, H., and Jospin, M. (2011). The α1 subunit EGL-19, the α2/δ subunit UNC-36, and the β subunit CCB-1 underlie voltage-dependent calcium currents in Caenorhabditis elegans striated muscle. J. Biol. Chem. 286, 36180–36187.

Lambert, S.A., Yang, A.W.H., Sasse, A., Cowley, G., Albu, M., Caddick, M.X., Morris, Q.D., Weirauch, M.T., and Hughes, T.R. (2019). Similarity regression predicts evolution of transcription factor sequence specificity. Nat. Genet. 51, 981–989.

Langfelder, P., and Horvath, S. (2008). WGCNA: An R package for weighted correlation network analysis. BMC Bioinformatics 9, 559.

Lanjuin, A., VanHoven, M.K., Bargmann, C.I., Thompson, J.K., and Sengupta, P. (2003). Otx/otd homeobox genes specify distinct sensory neuron identities in C. elegans. Dev. Cell 5, 621–633.

Lesch, B.J., Gehrke, A.R., Bulyk, M.L., and Bargmann, C.I. (2009). Transcriptional regulation and stabilization of left-right neuronal identity in C. elegans. Genes Dev. 23, 345–368.

Li, H., and Durbin, R. (2009). Fast and accurate short read alignment with Burrows-Wheeler transform. Bioinformatics 25, 1754–1760.

Li, L., Veksler-Lublinsky, I., and Zinovyeva, A. (2019). HRPK-1, a conserved KH-domain protein, modulates microRNA activity during Caenorhabditis elegans development. PLoS Genet. 15, e1008067.

Li, W., Kennedy, S.G., and Ruvkun, G. (2003). daf-28 encodes a C. elegans insulin superfamily member that is regulated by environmental cues and acts in the DAF-2 signaling pathway. Genes Dev. 17, 844–858.

Lorenzo, R., Onizuka, M., Defrance, M., and Laurent, P. (2020). Combining single-cell RNA-sequencing with a molecular atlas unveils new markers for Caenorhabditis elegans neuron classes. Nucleic Acids Res. 48, 7119–7134.

Love, M.I., Huber, W., and Anders, S. (2014). Moderated estimation of fold change and dispersion for RNA-seq data with DESeq2. Genome Biol. 15, 550.

Lun, A.T.L., Riesenfeld, S., Andrews, T., Dao, T.P., Gomes, T., and Marioni, J.C. (2019). EmptyDrops: Distinguishing cells from empty droplets in droplet-based single-cell RNA sequencing data. Genome Biol. 20, 63.

Masoudi, N., Tavazoie, S., Glenwinkel, L., Ryu, L., Kim, K., and Hobert, O. (2018). Unconventional function of an Achaete-Scute homolog as a terminal selector of nociceptive neuron identity. PLoS Biol. 16, e2004979.

McCarthy, D.J., Campbell, K.R., Lun, A.T.L., and Wills, Q.F. (2017). Scater: Pre-processing, quality control, normalization and visualization of single-cell RNA-seq data in R. Bioinformatics 33, 1179–1186.

McInnes, L., Healy, J., Saul, N., and Großberger, L. (2018). UMAP: Uniform Manifold Approximation and Projection. J. Open Source Softw. 3.

Melentijevic, I., Toth, M.L., Arnold, M.L., Guasp, R.J., Harinath, G., Nguyen, K.C., Taub, D., Parker, J.A., Neri, C., Gabel, C. V., et al. (2017). C. elegans neurons jettison protein aggregates and mitochondria under neurotoxic stress. Nature 542, 367–371.

Melkman, T., and Sengupta, P. (2005). Regulation of chemosensory and GABAergic motor neuron development by the C. elegans Aristaless/Arx homolog alr-1. Development 132, 1935– 1949.

Mendel, J.E., Korswagen, H.C., Liu, K.S., Hajdu-Cronin, Y.M., Simon, M.I., Plasterk, R.H.A., and Sternberg, P.W. (1995). Participation of the protein Go in multiple aspects of behavior in C. elegans. Science 267, 1652–1655.

Milo, R., Kashtan, N., Itzkovitz, S., Newman, M.E.J., and Alon, U. (2003). On the uniform generation of random graphs with prescribed degree sequences. ArXiv https://arxiv.org/abs/cond-mat/0312028.

Mimi Zhou, H., and Walthall, W.W. (1998). UNC-55, an orphan nuclear hormone receptor, orchestrates synaptic specificity among two classes of motor neurons in Caenorhabditis elegans. J. Neurosci. 18, 10438–10444.

Mirabeau, O., and Joly, J.S. (2013). Molecular evolution of peptidergic signaling systems in bilaterians. Proc. Natl. Acad. Sci. U. S. A. 110, E2028–37.

Moghal, N., Garcia, L.R., Khan, L.A., Iwasaki, K., and Sternberg, P.W. (2003). Modulation of EGF receptor-mediated vulva development by the heterotrimeric G-protein G q and excitable α cells in C. elegans. Development 130, 4553–4566.

Mok, D.Z.L., Sternberg, P.W., and Inoue, T. (2015). Morphologically defined sub-stages of C. Elegans vulval development in the fourth larval stage. BMC Dev. Biol. 15.

Moresco, J.J., and Koelle, M.R. (2004). Activation of EGL-47, a Gαo-coupled receptor, inhibits function of hermaphrodite-specific motor neurons to regulate Caenorhabditis elegans egg-laying behavior. J. Neurosci. 24, 8522–8530.

Moyle, M.W., Barnes, K.M., Kuchroo, M., Gonopolskiy, A., Duncan, L.H., Sengupta, T., Shao, L., Guo, M., Santella, A., Christensen, R., et al. (2020). Structural and developmental principles of neuropil assembly in C. elegans. BioRxiv https://doi.org/10.1101/2020.03.15.992222.

Murphy, C.T., and Hu, P.J. (2013). Insulin/insulin-like growth factor signaling in C. elegans. WormBook 1–43.

Murru, L., Moretto, E., Martano, G., and Passafaro, M. (2018). Tetraspanins shape the synapse. Mol. Cell. Neurosci. 91, 76–81.

Nguyen, J.P., Shipley, F.B., Linder, A.N., Plummer, G.S., Liu, M., Setru, S.U., Shaevitz, J.W., and Leifer, A.M. (2016). Whole-brain calcium imaging with cellular resolution in freely behaving Caenorhabditis elegans. Proc. Natl. Acad. Sci. U. S. A. 113, E1074–81.

Norris, A.D., Gao, S., Norris, M.L., Ray, D., Ramani, A.K., Fraser, A.G., Morris, Q., Hughes, T.R., Zhen, M., and Calarco, J.A. (2014). A Pair of RNA-binding proteins controls networks of splicing events contributing to specialization of neural cell types. Mol. Cell 54, 946–959.

O’Connor, T.P., Cockburn, K., Wang, W., Tapia, L., Currie, E., and Bamji, S.X. (2009). Semaphorin 5B mediates synapse elimination in hippocampal neurons. Neural Dev. 4.

Oranth, A., Schultheis, C., Tolstenkov, O., Erbguth, K., Nagpal, J., Hain, D., Brauner, M., Wabnig, S., Steuer Costa, W., McWhirter, R.D., et al. (2018). Food Sensation Modulates Locomotion by Dopamine and Neuropeptide Signaling in a Distributed Neuronal Network. Neuron 100, 1414–1428.

Ortiz, C., Etchberger, J., Posy, S., Frokjaer-Jensen, C., Lockery, S., Honig, B., and Hobert, O. (2006). Searching for neuronal left/right asymmetry: Genome wide analysis of nematode receptor-type guanylyl cyclases. Genetics 173, 131–149.

Packer, J.S., Zhu, Q., Huynh, C., Sivaramakrishnan, P., Preston, E., Dueck, H., Stefanik, D., Tan, K., Trapnell, C., Kim, J., et al. (2019). A lineage-resolved molecular atlas of C. Elegans embryogenesis at single-cell resolution. Science 365.

Patrick, R., Humphreys, D.T., Janbandhu, V., Oshlack, A., Ho, J.W.K., Harvey, R.P., and Lo, K.K. (2020). Sierra: Discovery of differential transcript usage from polyA-captured single-cell RNA-seq data. Genome Biol. 21.

Pavy, N., Rombauts, S., Déhais, P., Mathé, C., Ramana, D.V.V., Leroy, P., and Rouzé, P. (1999). Evaluation of gene prediction software using a genomic data set: Application to Arabidopsis thaliana sequences. Bioinformatics 15, 887–899.

Pereira, L., Kratsios, P., Serrano-Saiz, E., Sheftel, H., Mayo, A.E., Hall, D.H., White, J.G., LeBoeuf, B., Garcia, L.R., Alon, U., et al. (2015). A cellular and regulatory map of the cholinergic nervous system of C. Elegans. Elife 4, e12432.

Petersen, S.C., Watson, J.D., Richmond, J.E., Sarov, M., Walthall, W.W., and Miller, D.M. (2011). A transcriptional program promotes remodeling of GABAergic synapses in Caenorhabditis elegans. J. Neurosci. 31, 15362–15375.

Pierce-Shimomura, J.T., Faumont, S., Gaston, M.R., Pearson, B.J., and Lockery, S.R. (2001). The homeobox gene lim-6 is required for distinct chemosensory representations in C. elegans. Nature 410, 694–698.

Pierce, S.B., Costa, M., Wisotzkey, R., Devadhar, S., Homburger, S.A., Buchman, A.R., Ferguson, K.C., Heller, J., Platt, D.M., Pasquinelli, A.A., et al. (2001). Regulation of DAF-2 receptor signaling by human insulin and ins-1, a member of the unusually large and diverse C. elegans insulin gene family. Genes Dev. 15, 672–686.

Pols, M.S., and Klumperman, J. (2009). Trafficking and function of the tetraspanin CD63. Exp. Cell Res. 315, 1584–1592.

Poon, V.Y., Klassen, M.P., and Shen, K. (2008). UNC-6/netrin and its receptor UNC-5 locally exclude presynaptic components from dendrites. Nature 455, 669–673.

Poulin, J.F., Tasic, B., Hjerling-Leffler, J., Trimarchi, J.M., and Awatramani, R. (2016). Disentangling neural cell diversity using single-cell transcriptomics. Nat. Neurosci. 19, 1131– 1141.

Qiu, X., Mao, Q., Tang, Y., Wang, L., Chawla, R., Pliner, H.A., and Trapnell, C. (2017a). Reversed graph embedding resolves complex single-cell trajectories. Nat. Methods 14, 979–982.

Qiu, X., Hill, A., Packer, J., Lin, D., Ma, Y.A., and Trapnell, C. (2017b). Single-cell mRNA quantification and differential analysis with Census. Nat. Methods 14, 309–315.

Raj, B., and Blencowe, B.J. (2015). Alternative Splicing in the Mammalian Nervous System: Recent Insights into Mechanisms and Functional Roles. Neuron 87, 14–27.

Reilly, M.B., Cros, C., Varol, E., Yemini, E., and Hobert, O. (2020). Unique homeobox codes delineate all the neuron classes of C. elegans. Nature 584, 595–601.

Sanes, J.R., and Zipursky, S.L. (2020). Synaptic Specificity, Recognition Molecules, and Assembly of Neural Circuits. Cell 181, 536–556.

Satterlee, J.S., Sasakura, H., Kuhara, A., Berkeley, M., Mori, I., and Sengupta, P. (2001). Specification of thermosensory neuron fate in C. elegans requires ttx-1, a homolog of otd/Otx. Neuron 31, 943–956.

Sengupta, P., Colbert, H.A., and Bargmann, C.I. (1994). The C. elegans gene odr-7 encodes an olfactory-specific member of the nuclear receptor superfamily. Cell 79, 971–980.

Sengupta, P., Chou, J.H., and Bargmann, C.I. (1996). odr-10 Encodes a seven transmembrane domain olfactory receptor required for responses to the odorant diacetyl. Cell 84, 899–909.

Serrano-Saiz, E., Poole, R.J., Felton, T., Zhang, F., De La Cruz, E.D., and Hobert, O. (2013). Modular control of glutamatergic neuronal identity in C. elegans by distinct homeodomain proteins. Cell 155, 659–673.

Shan, G., Kim, K., Li, C., and Walthall, W.W. (2005). Convergent genetic programs regulate similarities and differences between related motor neuron classes in Caenorhabditis elegans. Dev. Biol. 280, 494–503.

Shen, K., and Bargmann, C.I. (2003). The immunoglobulin superfamily protein SYG-1 determines the location of specific synapses in C. elegans. Cell 112, 619–630.

Siebert, S., Farrell, J.A., Cazet, J.F., Abeykoon, Y., Primack, A.S., Schnitzler, C.E., and Juliano, C.E. (2019). Stem cell differentiation trajectories in Hydra resolved at single-cell resolution. Science 365.

Siegenthaler, D., Enneking, E.M., Moreno, E., and Pielage, J. (2015). L1CAM/Neuroglian controls the axon-axon interactions establishing layered and lobular mushroom body architecture. J. Cell Biol. 208, 1003–1018.

Sigrist, C.J.A., De Castro, E., Cerutti, L., Cuche, B.A., Hulo, N., Bridge, A., Bougueleret, L., and Xenarios, I. (2013). New and continuing developments at PROSITE. Nucleic Acids Res. 41, D344–7.

Smith, S.J., Smbül, U., Graybuck, L.T., Collman, F., Seshamani, S., Gala, R., Gliko, O., Elabbady, L., Miller, J.A., Bakken, T.E., et al. (2019). Single-cell transcriptomic evidence for dense intracortical neuropeptide networks. Elife 8, e47889.

Soneson, C., and Delorenzi, M. (2013). A comparison of methods for differential expression analysis of RNA-seq data. BMC Bioinformatics 14.

Spead, O., and Poulain, F.E. (2020). Trans-axonal signaling in neural circuit wiring. Int. J. Mol. Sci. 21, 5170.

Spencer, W.C., McWhirter, R., Miller, T., Strasbourger, P., Thompson, O., Hillier, L.D.W., Waterston, R.H., and Miller, D.M. (2014). Isolation of specific neurons from C. Elegans larvae for gene expression profiling. PLoS One 9, e112102.

Sperry, R.W. (1963). Chemoaffinity in theorderly growth of nerve fiber paterns and connections. Proc. Natl. Acad. Sci. USA 50, 703–710.

Stefanakis, N., Carrera, I., and Hobert, O. (2015). Regulatory Logic of Pan-Neuronal Gene Expression in C. elegans. Neuron 87, 733–750.

Von Stetina, S.E., Fox, R.M., Watkins, K.L., Starich, T.A., Shaw, J.E., and Miller, D.M. (2007). UNC-4 represses CEH-12/HB9 to specify synaptic inputs to VA motor neurons in C. elegans. Genes Dev. 21, 332–346.

Stuart, T., Butler, A., Hoffman, P., Hafemeister, C., Papalexi, E., Mauck, W.M., Hao, Y., Stoeckius, M., Smibert, P., and Satija, R. (2019). Comprehensive Integration of Single-Cell Data. Cell 177, 1888–1902.

Sulston, J.E., and Horvitz, H.R. (1977). Post-embryonic cell lineages of the nematode, Caenorhabditis elegans. Dev. Biol. 56, 110–156.

Sulston, J.E., Schierenberg, E., White, J.G., and Thomson, J.N. (1983). The embryonic cell lineage of the nematode Caenorhabditis elegans. Dev. Biol. 100, 64–119.

Swoboda, P., Adler, H.T., and Thomas, J.H. (2000). The RFX-type transcription factor DAF-19 regulates sensory neuron cilium formation in C. Elegans. Mol. Cell 5, 411–421.

Tamburino, A.M., Ryder, S.P., and Walhout, A.J.M. (2013). A compendium of Caenorhabditis elegans RNA binding proteins predicts extensive regulation at multiple levels. G3 Genes, Genomes, Genet. 3, 297–304.

Tasic, B., Menon, V., Nguyen, T.N., Kim, T.K., Jarsky, T., Yao, Z., Levi, B., Gray, L.T., Sorensen, S.A., Dolbeare, T., et al. (2016). Adult mouse cortical cell taxonomy revealed by single cell transcriptomics. Nat. Neurosci. 19, 335–346.

Thompson, M., Bixby, R., Dalton, R., Vandenburg, A., Calarco, J.A., and Norris, A.D. (2019). Splicing in a single neuron is coordinately controlled by RNA binding proteins and transcription factors. Elife 8, e46726.

Tomioka, M., Naito, Y., Kuroyanagi, H., and Iino, Y. (2016). Splicing factors control C. elegans behavioural learning in a single neuron by producing DAF-2c receptor. Nat. Commun. 7, 11645.

Tourasse, N.J., Millet, J.R.M., and Dupuy, D. (2017). Quantitative RNA-seq meta-analysis of alternative exon usage in C. elegans. Genome Res. 27, 2120–2128.

Tran, T.S., Rubio, M.E., Clem, R.L., Johnson, D., Case, L., Tessier-Lavigne, M., Huganir, R.L., Ginty, D.D., and Kolodkin, A.L. (2009). Secreted semaphorins control spine distribution and morphogenesis in the postnatal CNS. Nature 462, 1065–1069.

Trapnell, C., Cacchiarelli, D., Grimsby, J., Pokharel, P., Li, S., Morse, M., Lennon, N.J., Livak, K.J., Mikkelsen, T.S., and Rinn, J.L. (2014). The dynamics and regulators of cell fate decisions are revealed by pseudotemporal ordering of single cells. Nat. Biotechnol. 32, 381–386.

Treisman, J., Gönczy, P., Vashishtha, M., Harris, E., and Desplan, C. (1989). A single amino acid can determine the DNA binding specificity of homeodomain proteins. Cell 59, 553–562.

Troemel, E.R., Chou, J.H., Dwyer, N.D., Colbert, H.A., and Bargmann, C.I. (1995). Divergent seven transmembrane receptors are candidate chemosensory receptors in C. elegans. Cell 83, 207–218.

Troemel, E.R., Sagasti, A., and Bargmann, C.I. (1999). Lateral signaling mediated by axon contact and calcium entry regulates asymmetric odorant receptor expression in C. elegans. Cell 99, 387–398.

Turner, A.J., Elwyn Isaac, R., and Coates, D. (2001). The neprilysin (NEP) family of zinc metalloendopeptidases: Genomics and function. BioEssays 23, 261–269.

Varadan, V., Miller, D.M., and Anastassiou, D. (2006). Computational inference of the molecular logic for synaptic connectivity in C. elegans. Bioinformatics 22, e497–506.

Venkatachalam, V., Ji, N., Wang, X., Clark, C., Mitchell, J.K., Klein, M., Tabone, C.J., Florman, J., Ji, H., Greenwood, J., et al. (2016). Pan-neuronal imaging in roaming Caenorhabditis elegans. Proc. Natl. Acad. Sci. U. S. A. 113, E1082–8.

Vidal, B., Aghayeva, U., Sun, H., Wang, C., Glenwinkel, L., Bayer, E.A., and Hobert, O. (2018). An atlas of Caenorhabditis elegans chemoreceptor expression. PLoS Biol. 16, e2004218.

Vuong, C.K., Black, D.L., and Zheng, S. (2016). The neurogenetics of alternative splicing. Nat. Rev. Neurosci. 17, 265–281.

Wang, L., Klein, R., Zheng, B., and Marquardt, T. (2011). Anatomical Coupling of Sensory and Motor Nerve Trajectory via Axon Tracking. Neuron 71, 263–277.

Weirauch, M.T., Yang, A., Albu, M., Cote, A.G., Montenegro-Montero, A., Drewe, P., Najafabadi, H.S., Lambert, S.A., Mann, I., Cook, K., et al. (2014). Determination and inference of eukaryotic transcription factor sequence specificity. Cell 158, 1431–1443.

White, J., Southgate, E., Thomson, J.N., and Brenner, S. (1986). The structure of the nervous system of the nematode Caenorhabditis elegans. Philos. Trans. R. Soc. London. B, Biol. Sci. 314, 1–340.

Witvliet, D., Mulcahy, B., Mitchell, J.K., Meirovitch, Y., Berger, D.R., Wu, Y., Liu, Y., Koh, W.X., Parvathala, R., Holmyard, D., et al. (2020). Connectomes across development reveal principles of brain maturation in C. elegans. BioRxiv https://doi.org/10.1101/2020.04.30.066209.

Yemini, E., Lin, A., Nejatbakhsh, A., Varol, E., Sun, R., Mena, G.E., Samuel, A.D.T., Paninski, L., Venkatachalam, V., and Hobert, O. (2019). NeuroPAL: A Neuronal Polychromatic Atlas of Landmarks for Whole-Brain Imaging in C. elegans. BioRxiv https://doi.org/10.1101/676312.

Young, M.D., and Behjati, S. (2020). SoupX removes ambient RNA contamination from droplet based single cell RNA sequencing data. BioRxiv https://doi.org/10.1101/303727.

Yu, S., Avery, L., Baude, E., and Garbers, D.L. (1997). Guanylyl cyclase expression in specific sensory neurons: A new family of chemosensory receptors. Proc. Natl. Acad. Sci. U. S. A. 94, 3384–3387.

Zeisel, A., Munoz-Manchado, A.B., Codeluppi, S., Lönnerberg, P., Manno, G. La, Juréus, A., Marques, S., Munguba, H., He, L., Betsholtz, C., et al. (2015). Brain structure. Cell types in the mouse cortex and hippocampus revealed by single-cell RNA-seq. Science 347, 1138–1142.

Zhang, S., Banerjee, D., and Kuhn, J.R. (2011). Isolation and culture of larval cells from C. elegans. PLoS One 6, e19505.

Zhou, X., and Bessereau, J.L. (2019). Molecular Architecture of Genetically-Tractable GABA Synapses in C. elegans. Front. Mol. Neurosci. 12.

Zhu, Y., Sousa, A.M.M., Gao, T., Skarica, M., Li, M., Santpere, G., Esteller-Cucala, P., Juan, D., Ferrández-Peral, L., Gulden, F.O., et al. (2018). Spatiotemporal transcriptomic divergence across human and macaque brain development. Science 362.

