## Supplemental Figures for "Molecular topography of an entire nervous system"

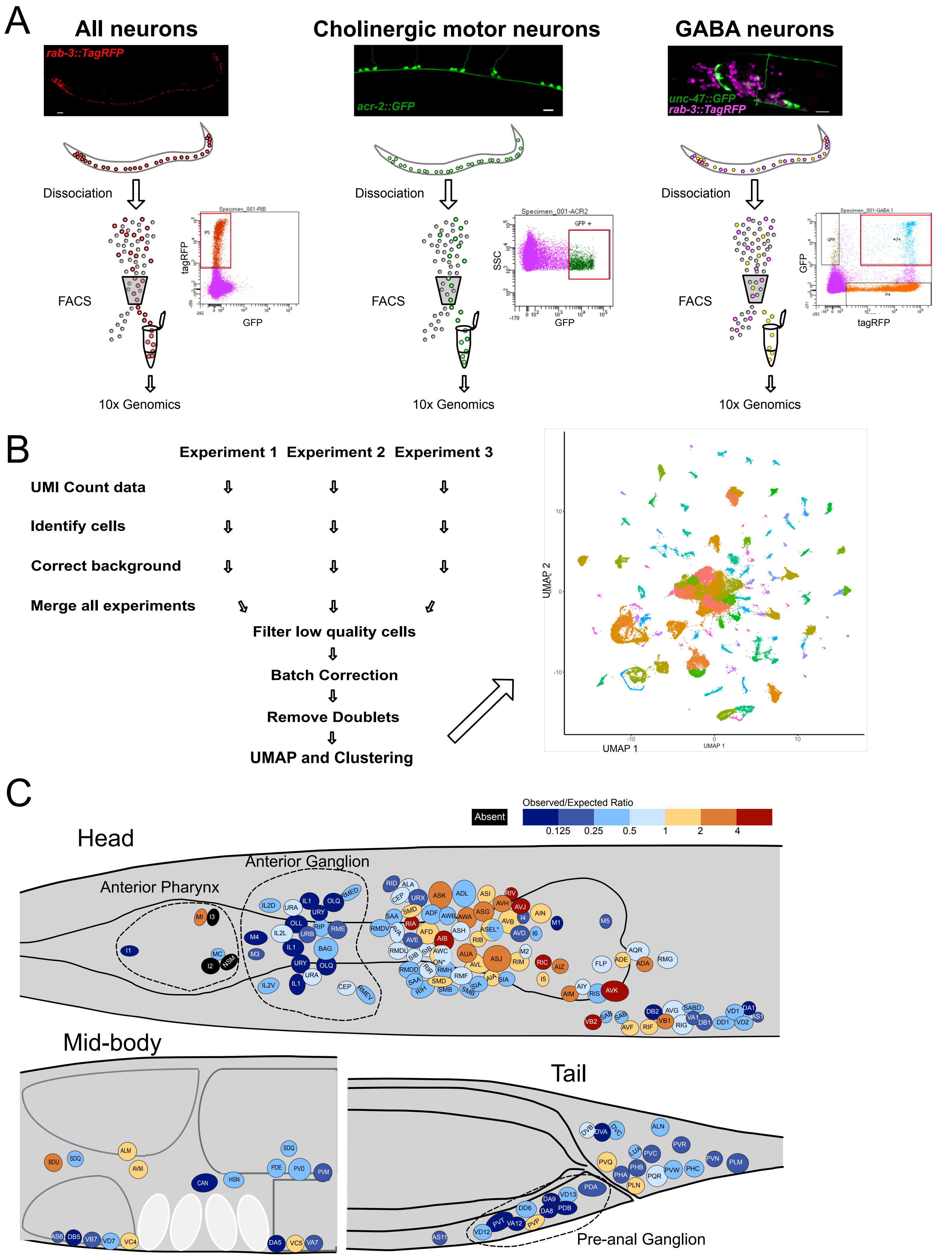

A
All neurons
rab-3::TagRFP
Cholinergic motor neurons
acr-2::GFP
GABA neurons
unc-47::GFP
rab-3::TagRFP
Dissociation
FACS
tagRFP
GFP
SSC
GFP +
10x Genomics
B
Experiment 1
Experiment 2
Experiment 3
UMI Count data
Identify cells
Correct background
Merge all experiments
Filter low quality cells
Batch Correction
Remove Doublets
UMAP and Clustering
UMAP 2
UMAP 1
C
Absent
Observed/Expected Ratio
0.125
0.25
0.5
1
2
4
Head
Anterior Ganglion
Anterior Pharynx
Mid-body
Tail
Pre-anal Ganglion

**Figure S1. Isolating L4 larval stage neurons from fluorescent marker strains for single-cell RNA-Sequencing (scRNA-Seq)**. (**A**) Confocal images of the pan-neural marker, *rab-3::TagRFP* (*otIs355)*, ventral cord cholinergic motor neurons marked with *acr-2::GFP (juIs14)* and neurons in the head region dual-labeled with the GABA neuron-specific reporter, *unc-47::GFP (oxIs12)* and the pan neural marker strain, *rab-3::TagRFP*. Scale bars = 10 µm. L4 animals were treated with SDS-DTT and dissociated with pronase to produce single-cell suspensions. Targeted subgroups of neurons were isolated by Fluorescence Activated Cell Sorting (FACS, red boxes) and collected for scRNA-Seq using the 10x Genomics 3' platform. B) Results from 17 separate profiling experiments were submitted to a series of processing steps to produce a final merged data set (see Methods). Right panel shows UMAP projection of all 100,955 cells, colored by cluster. C) Graphical depiction of relative abundance of each neuron class in cells isolated from the pan-neural marker strain (*rab-3::TagRFP*). The fraction of observed cells for each neuron type was divided by the expected ratio (i.e., # neuron type/302) and annotated for each cell type according to heat map index. Note under-representation (dark blue/black) of neurons in the anterior pharynx, anterior ganglion and pre-anal ganglion (dashed lines). * For pairs ASE and AWC, the individual neurons are treated separately. Related to Figure 1, 2.

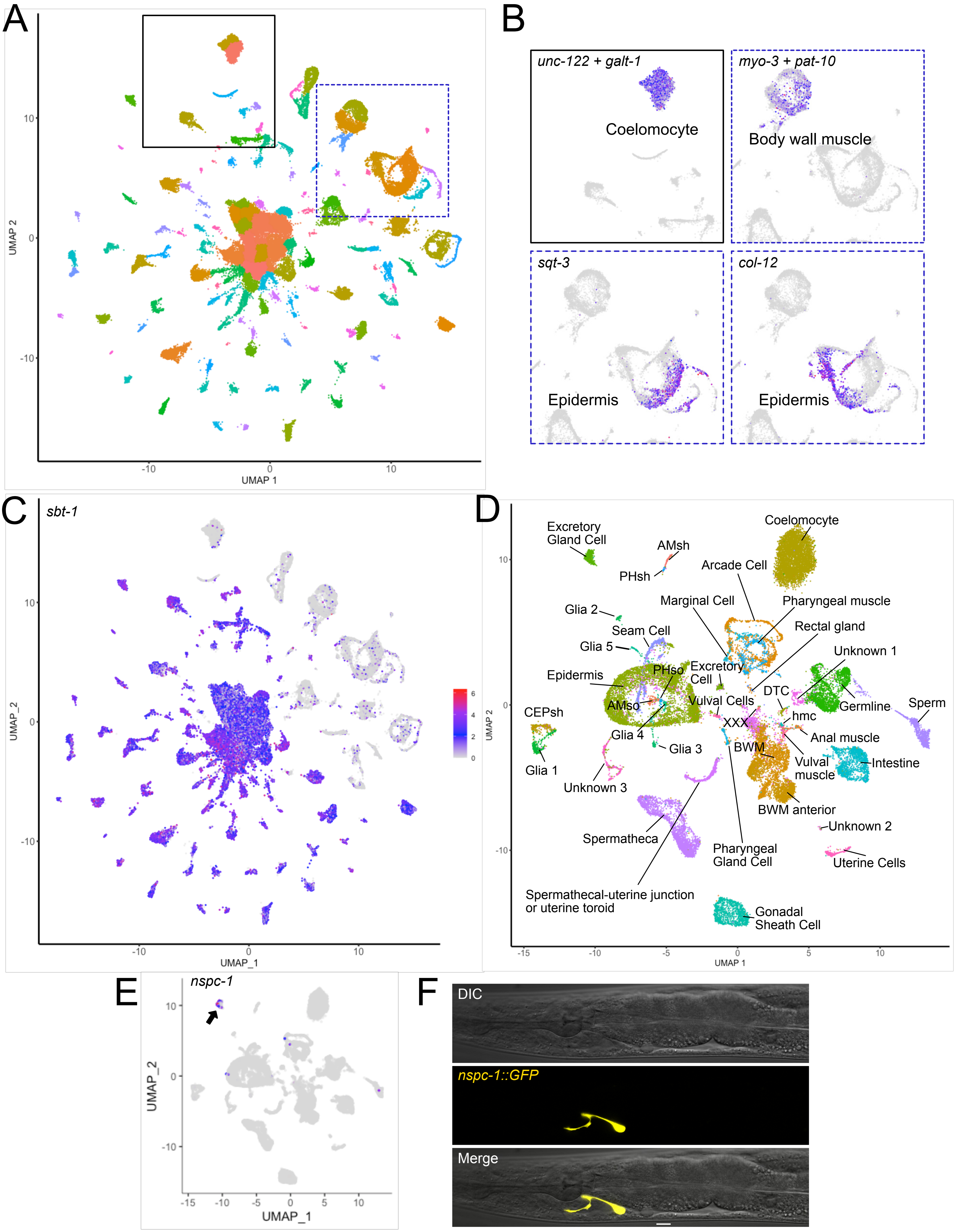
A
UMAP 2
UMAP 1
B
unc-122 + galt-1
Coelomocyte
myo-3 + pat-10
Body wall muscle
sqt-3
Epidermis
col-12
Epidermis
C
sbt-1
UMAP_2
UMAP_1
D
Excretory Gland Cell
AMsh
PHsh
Coelomocyte
Arcade Cell
Marginal Cell
Pharyngeal muscle
Glia 2
Seam Cell
Rectal gland
Glia 5
Unknown 1
Epidermis
PHso
Excretory Cell
DTC
Germline
Sperm
CEPsh
AMso
Vulval Cells
XXX
hmc
Anal muscle
Glia 4
Glia 3
BWM
Vulval muscle
Intestine
Glia 1
Unknown 3
BWM anterior
Unknown 2
Spermatheca
Pharyngeal Gland Cell
Uterine Cells
Spermathecal-uterine junction or uterine toroid
Gonadal Sheath Cell
UMAP 2
UMAP 1
E
nspc-1
UMAP_2
UMAP_1
F
DIC
nspc-1::GFP
Merge

**Figure S2. Distinguishing non-neuronal vs neuronal cells.** A) UMAP projection of 100,955 single cell profiles, colored by cluster. B) Known markers identify clusters of non-neuronal cells (*unc-122* + *galt-1*, coelomocytes; *myo-3* + *pat-10*, body wall muscles; *sqt-3* and *col-12*, epidermis). Boxes correspond to sub-regions in A. C) The neuropeptide processing gene *sbt-1* is expressed in all neuronal clusters (blue-magenta) and largely absent from non-neuronal clusters (gray). D) Sub-UMAP of all non-neuronal cells, labeled by cell type. E) *nspc-1*, a member of the nematode-specific peptide c (*nspc*) gene family, is restricted to a single non-neuronal cluster (arrow). F) The transcriptional reporter *nspc-1::GFP* is exclusively expressed in the excretory gland cell (yellow). DIC (Differential Interference Contrast). Scale bar = 10 μm. Related to Figure 1.

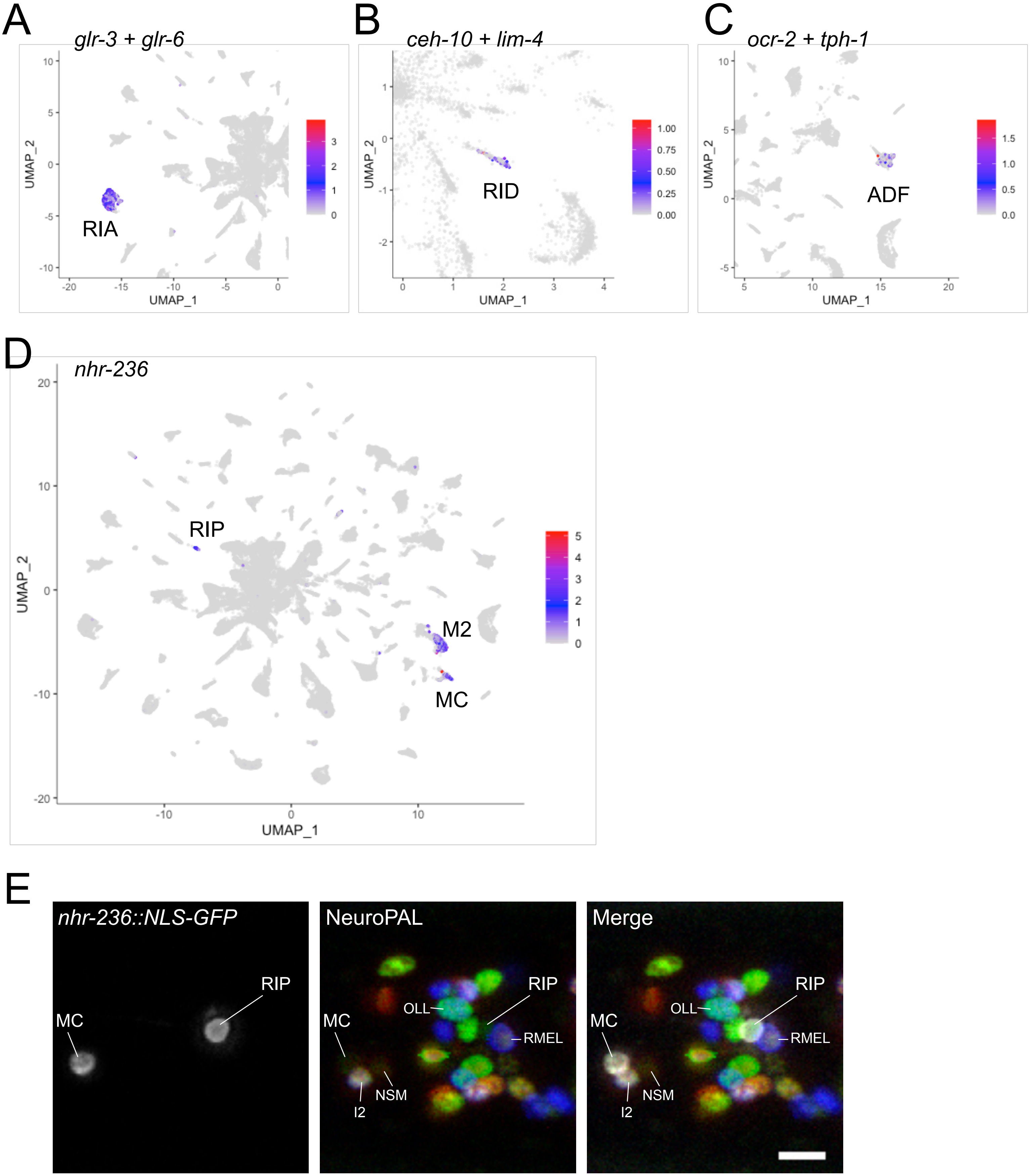

A
glr-3 + glr-6
RIA
UMAP_1
UMAP_2
B
ceh-10 + lim-4
RID
UMAP_1
UMAP_2
C
ocr-2 + tph-1
ADF
UMAP_1
UMAP_2
D
nhr-236
RIP
M2
MC
UMAP_1
UMAP_2
E
nhr-236::NLS-GFP
NeuroPAL
Merge
MC
RIP
OLL
RMEL
NSM
I2

**Figure S3. Annotation of neuronal clusters.** A) Co-expression of the glutamate receptor genes *glr-3* and *glr-6* demarcate the RIA cluster. B) Co-expression of the homeodomain transcription factors *ceh-10* and *lim-4* label the RID cluster. C) Co-expression of the transient receptor potential channel (trp) gene *ocr-2* and tryptophan hydroxylase *tph-1* label the sensory neuron ADF. D) The nuclear hormone receptor *nhr-236* is primarily detected in three clusters, corresponding to RIP, M2, and MC. E) Z-projection of confocal stack showing *nhr-236::NLS-GFP* expression in RIP and MC in a NeuroPAL strain. M2 (not shown) also consistently expressed GFP. Scale bar = 5 µm. Related to Figure 2.

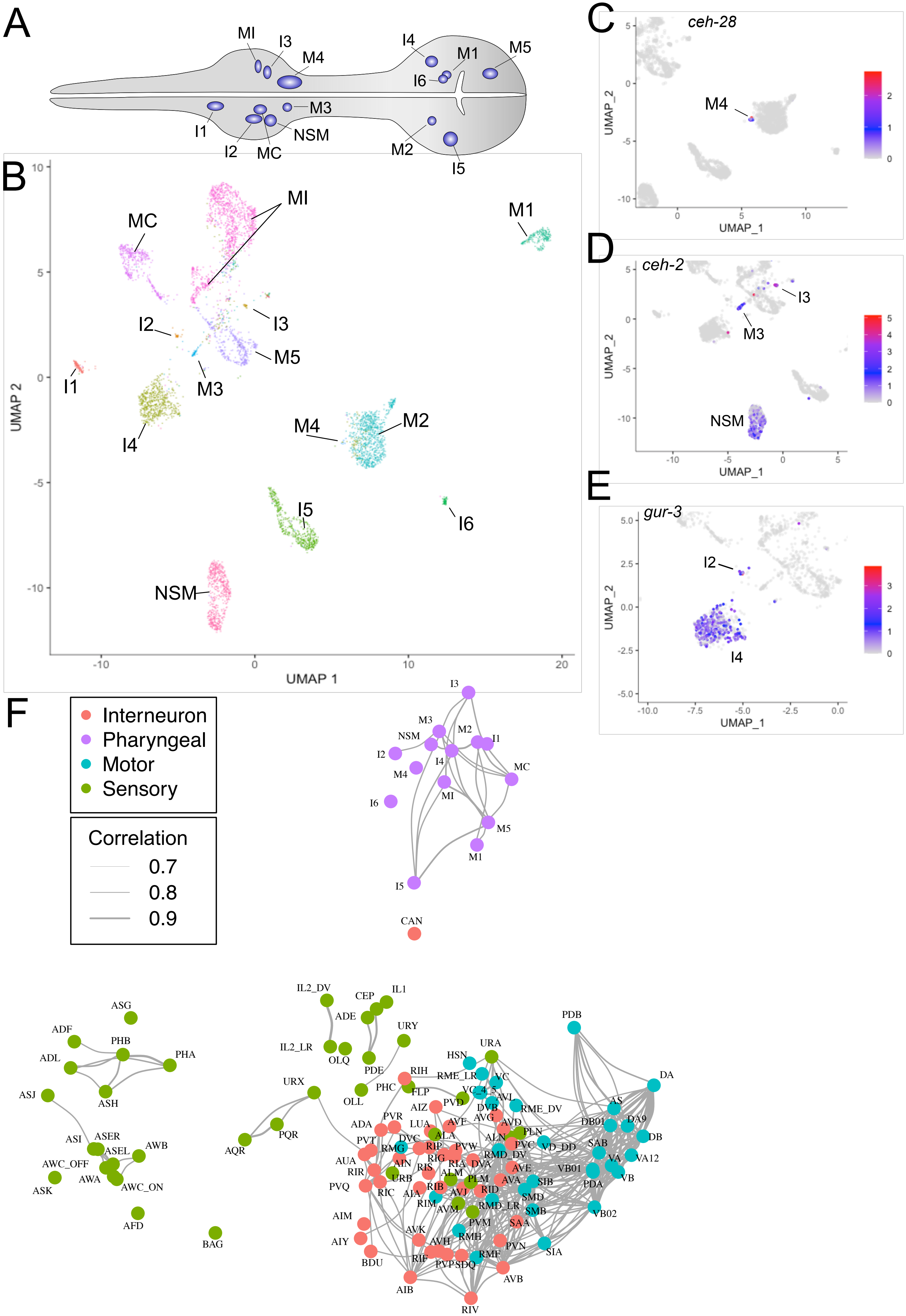
A
MI
I3
M4
I4
M1
M5
I6
M3
I1
I2
MC
NSM
M2
I5
B
MI
MC
M1
I2
I3
M5
I1
M3
M4
M2
I4
I5
I6
NSM
UMAP 2
UMAP 1
C
ceh-28
M4
UMAP_2
UMAP_1
D
ceh-2
I3
M3
NSM
UMAP_2
UMAP_1
E
gur-3
I2
I4
UMAP_2
UMAP_1
F
Interneuron
Pharyngeal
Motor
Sensory
Correlation
0.7
0.8
0.9
I3
M3
NSM
M2
I1
I2
I4
MC
M4
MI
I6
M5
M1
I5
CAN
ASG
ADF
PHB
ADL
PHA
ASJ
ASH
ASI
ASER
ASEL
AWB
AWC_OFF
AWA
AWC_ON
ASK
AFD
BAG
IL2_DV
CEP
IL1
ADE
URY
IL2_LR
OLQ
PDE
URX
PQR
AQR
OLL
PHC
PDB
DA
URA
HSN
RIH
RME_LR
VC
FLP
VC_4_5
AVL
PVD
DVB
RME_DV
AIZ
AVG
AVD
AS
DA9
DB01
DB
PLN
LUA
AVF
ALA
ADA
PVR
PVT
DVC
ALN
PVC
VD_DD
SAB
VA12
RMG
RIP
PVW
RMD_DV
VA
AUA
RIG
AIN
RIS
RIA
DVA
AVE
VB01
VB
RIR
URB
ALM
PLM
AVA
SIB
PDA
PVQ
RIC
AIA
RIB
AVJ
RID
SMD
RIM
AVM
RMD_LR
SMB
VB02
AIM
PVM
SAA
AVK
RMH
AIY
AVH
PVN
SIA
RIF
PVP
SDQ
RMF
BDU
AVB
AIB
RIV

**Figure S4. Identifying pharyngeal neuron types and neuron clustering.** A) Diagram denoting neurons (blue) in pharynx (gray). B) Sub-UMAP of neurons expressing the pharyngeal neuron marker *ceh-34* revealed independent clusters for each known pharyngeal neuron type. C-E) Expression of known specific marker genes identifies individual pharyngeal neuron types. C) The M4-specific homeodomain transcription factor, *ceh-28* is exclusively detected in a small but distinct group of 12 cells. D) *ceh-2*, a marker for I3, M3 and NSM pharyngeal neurons, is restricted to 3 clusters. E) Restricted expression of *gur-3*, a marker for I2 and I4 pharyngeal neurons. F) Network analysis of all transcriptionally distinct neuron types and subtypes. *C. elegans* neuron types organized in a force-directed network according to transcriptomic similarities. Colors denote distinct neuron modalities, and widths of edges show the strengths of the transcriptome similarity between each pair of neuron types. Edges with Pearson correlation coefficients > 0.7 are shown. Related to Figure 2.

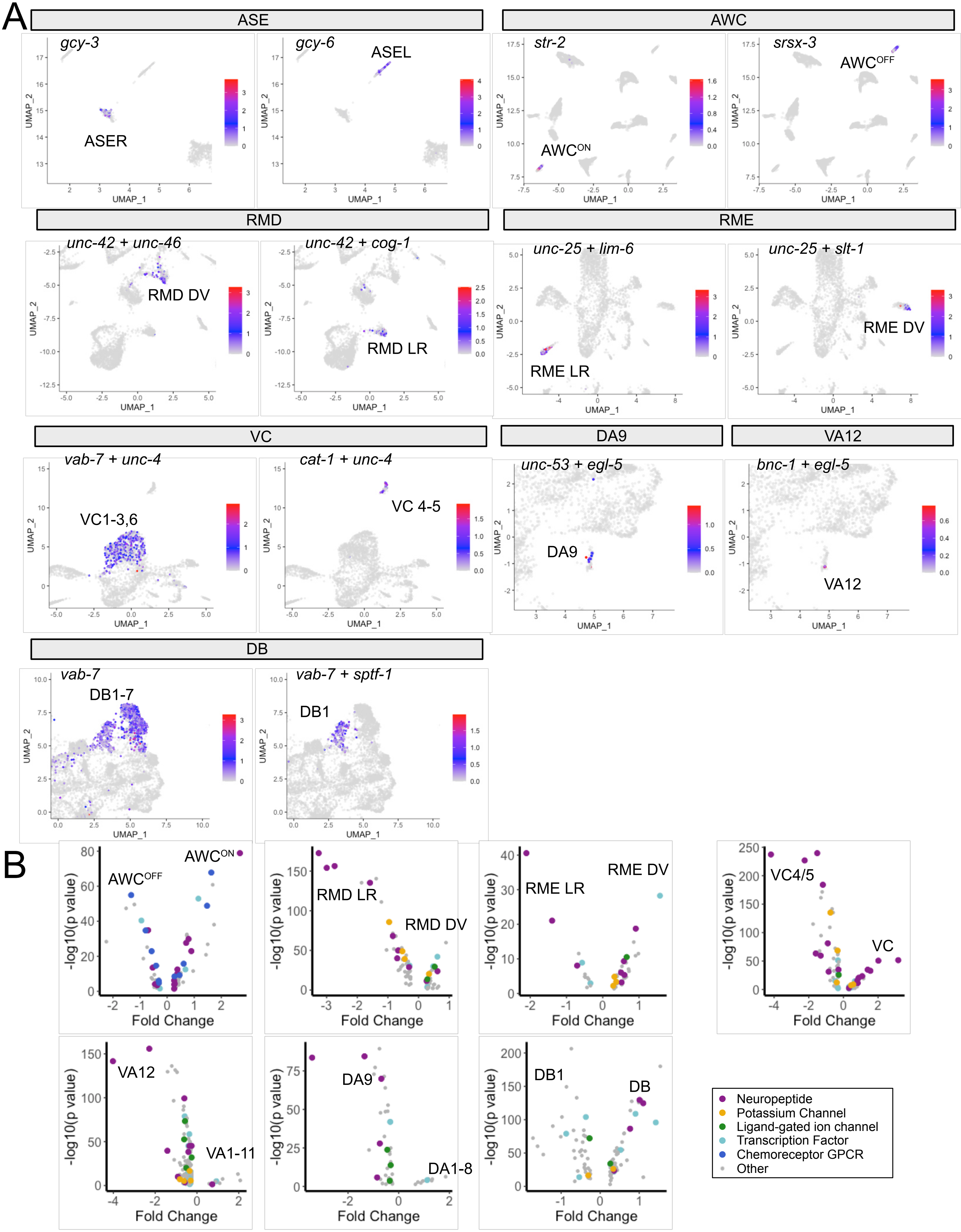

A
ASE
gcy-3
ASER
gcy-6
ASEL
AWC
str-2
AWC^ON
srsx-3
AWC^OFF
RMD
unc-42 + unc-46
RMD DV
unc-42 + cog-1
RMD LR
RME
unc-25 + lim-6
RME LR
unc-25 + slt-1
RME DV
VC
vab-7 + unc-4
VC1-3,6
cat-1 + unc-4
VC 4-5
DA9
unc-53 + egl-5
DA9
VA12
bnc-1 + egl-5
VA12
DB
vab-7
DB1-7
vab-7 + sptf-1
DB1
UMAP_1
UMAP_2
B
AWC^ON
AWC^OFF
RMD LR
RMD DV
RME LR
RME DV
VC4/5
VC
VA12
VA1-11
DA9
DA1-8
DB1
DB
-log10(p value)
Fold Change
Neuropeptide
Potassium Channel
Ligand-gated ion channel
Transcription Factor
Chemoreceptor GPCR
Other

**Figure S5. Identification of neuron sub-types.** A) Sub-UMAPs showing expression of known markers for neuron sub-types for ASE (*gcy-3*, ASER and *gcy-6*, ASEL), AWC (*str-2*, AWC$^{ON}$ and *srsx-3*, AWC$^{OFF}$), VC (*vab-7* + *unc-4*, VC1-3,6 and *cat-1* + *unc-4*, VC4-5), RMD (*unc-42* + *unc-46*, RMD DV and *unc-42* + *cog-1*, RMD LR), RME (*unc-25* + *lim-6*, RME LR and *unc-25* + *slt-1*, RME DV), DA9 (*unc-53* + *egl-5*), VA12 (*bnc-1* + *egl-5*), DB (*vab-7*, DB1-7 and *vab-7* + *sptf-1*, DB1). B) Volcano plots of genes that are differentially expressed between neuron subtypes. Inset depicts color coding for selected gene families. Related to Figure 3.

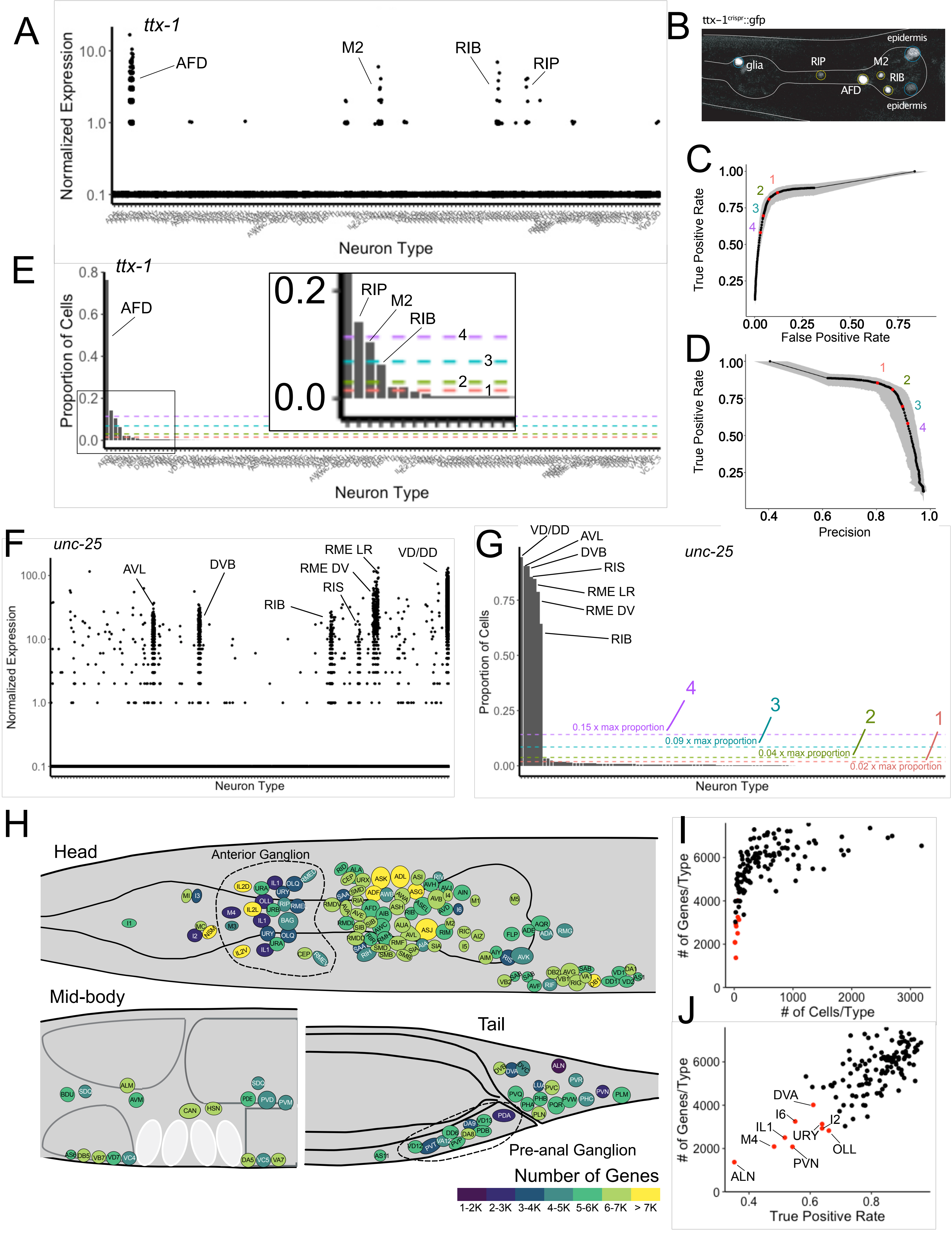
A
ttx-1
Normalized Expression
10.0
1.0
0.1
AFD
M2
RIB
RIP
Neuron Type
B
ttx-1crispr::gfp
epidermis
glia
RIP
M2
RIB
AFD
epidermis
C
True Positive Rate
False Positive Rate
1
2
3
4
D
True Positive Rate
Precision
E
ttx-1
Proportion of Cells
AFD
RIP
M2
RIB
4
3
2
1
Neuron Type
F
unc-25
Normalized Expression
AVL
DVB
RME LR
RME DV
RIS
RIB
VD/DD
Neuron Type
G
unc-25
Proportion of Cells
VD/DD
AVL
DVB
RIS
RME LR
RME DV
RIB
0.15 x max proportion
0.09 x max proportion
0.04 x max proportion
0.02 x max proportion
Neuron Type
H
Head
Anterior Ganglion
Mid-body
Tail
Pre-anal Ganglion
Number of Genes
1-2K
2-3K
3-4K
4-5K
5-6K
6-7K
> 7K
I
### of Genes/Type
### of Cells/Type
J
### of Genes/Type
True Positive Rate
DVA
I6
IL1
I2
M4
URY
OLL
PVN
ALN

**Figure S6. Establishing expression thresholds.** A) Jitter plot of normalized *ttx-1* expression (Y-axis) in all neuronal clusters (X-axis) shows strongest expression in AFD, M2, RIB and RIP. B) Confocal image of *ttx-1$^{crispr}$::GFP* shows expression in AFD, M2, RIB and RIP neurons and in glia and epidermal cells. C) Receiver-Operator Characteristic (ROC) curve of True Positive Rate (TPR) vs False Positive Rate (FPR) for a range of thresholds (1-4) (red dots) compared to ground truth expression data (see Methods). Increased stringency diminishes both the TPR and FPR. Grey shading represents 95% confidence intervals. D) Thresholds 1-4 (red dots) plotted on Precision-Recall (PR) Curve of Recall (TPR) vs Precision [1 – False Discovery Rate (FDR)]. Grey shading represents 95% confidence intervals. E) The proportion of cells in each neuron-specific cluster expressing *ttx-1*. Inset shows expanded view of boxed region. Thresholds (1-4) are set to different proportions of *ttx-1*-expressing cells in each cluster (see Methods). Note that neurons RIB and M2, that show expression of native *ttx-1$^{crispr}$::GFP,* are excluded by the thresholds 3 and 4. Numbers (1-4) correspond to thresholds in C and D. F) Jitter plot shows strong expression of *unc-25*/GAD in seven known GABAergic neuron types (AVL, DVB, RIB, RIS, RME, DD, VD) but scattered *unc-25/GAD* expression is also detected in other cell types. G) Bar graph plotting the proportion of cells in each neuron type that express *unc-25/GAD*. Thresholds of increasing stringency (1-4) are set to different proportions of cells in a given cluster that express *unc-25/GAD*. Note that threshold 2 distinguishes known *unc-25/GAD*-positive neurons from other neuron types with lower detected levels of *unc-25/GAD* (see Methods). H) Cell soma for each neuron type in the head, mid-body and tail regions are colored according to the number of genes detected using threshold 2. Neuron types in the anterior ganglion (dashed line) are among those with the fewest cells and also lower numbers of detected genes. I) Number of genes detected with threshold 2 for each neuron type plotted against the number of cells in each neuron-type cluster. Spearman's rank correlation

= 0.783, p < 2.2e-16). J) Number of genes detected with threshold 2 plotted against the True Positive Rate (TPR) for each neuron type. Spearman's rank correlation = 0.678, p < 2.2e-16. Neurons with fewest cells, low TPR and number of genes/type are denoted (red). Related to Figure 4, 5, S9-S12.

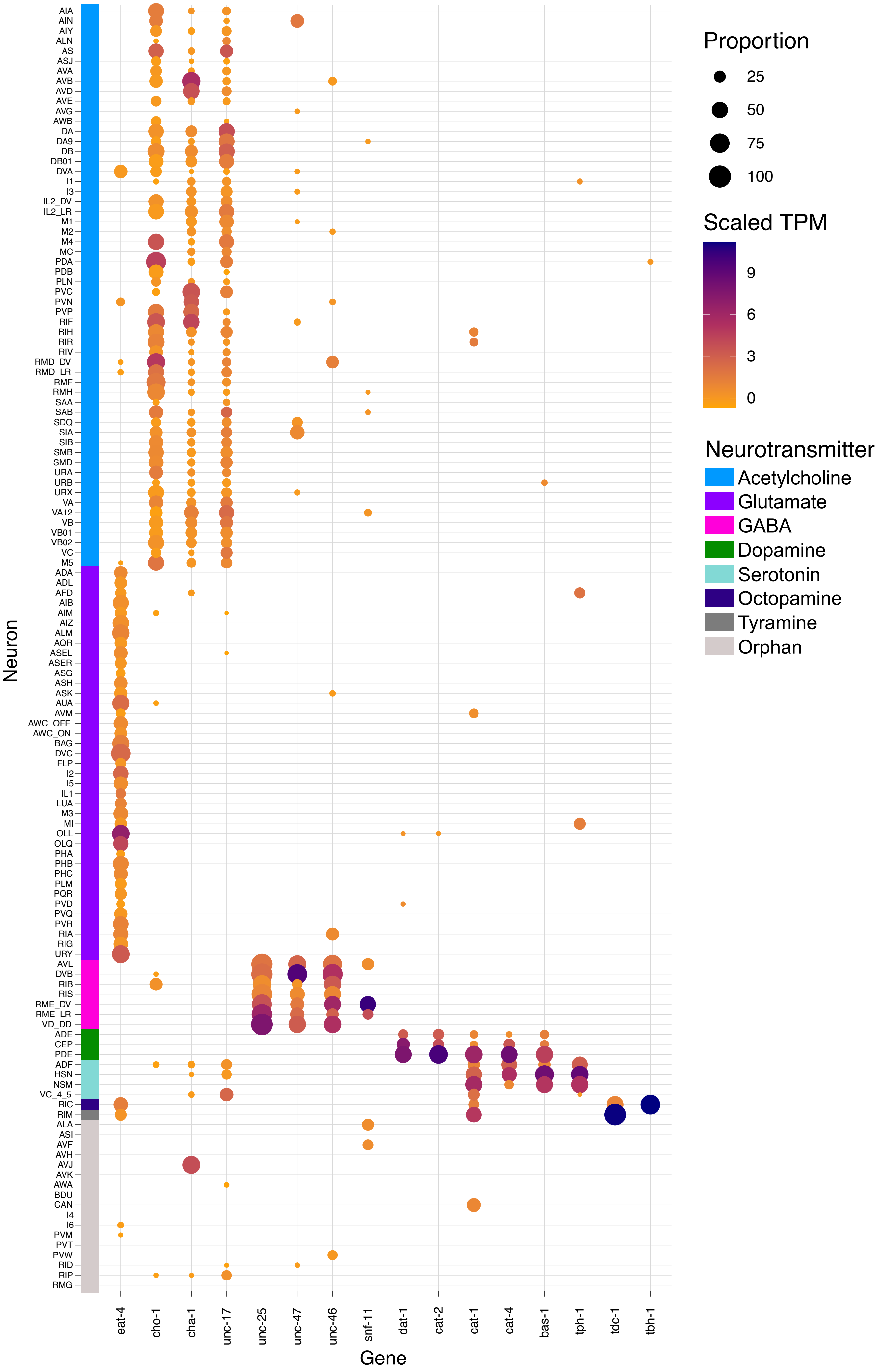

Proportion
25
50
75
100
Scaled TPM
9
6
3
0
Neurotransmitter
Acetylcholine
Glutamate
GABA
Dopamine
Serotonin
Octopamine
Tyramine
Orphan
Neuron
Gene
eat-4
cho-1
cha-1
unc-17
unc-25
unc-47
unc-46
snf-11
dat-1
cat-2
cat-1
cat-4
bas-1
tph-1
tdc-1
tbh-1
AIA
AIN
AIY
ALN
AS
ASJ
AVA
AVB
AVD
AVE
AVG
AWB
DA
DA9
DB
DB01
DVA
I1
I3
IL2_DV
IL2_LR
M1
M2
M4
MC
PDA
PDB
PLN
PVC
PVN
PVP
RIF
RIH
RIR
RIV
RMD_DV
RMD_LR
RMF
RMH
SAA
SAB
SDQ
SIA
SIB
SMB
SMD
URA
URB
URX
VA
VA12
VB
VB01
VB02
VC
M5
ADA
ADL
AFD
AIB
AIM
AIZ
ALM
AQR
ASEL
ASER
ASG
ASH
ASK
AUA
AVM
AWC_OFF
AWC_ON
BAG
DVC
FLP
I2
I5
IL1
LUA
M3
MI
OLL
OLQ
PHA
PHB
PHC
PLM
PQR
PVD
PVQ
PVR
RIA
RIG
URY
AVL
DVB
RIB
RIS
RME_DV
RME_LR
VD_DD
ADE
CEP
PDE
ADF
HSN
NSM
VC_4_5
RIC
RIM
ALA
ASI
AVF
AVH
AVJ
AVK
AWA
BDU
CAN
I4
I6
PVM
PVT
PVW
RID
RIP
RMG

**Figure S7. Expression of neurotransmitter-related genes.** Heatmap showing scaled expression within each neuron type for genes specifically required for mediating synthesis and release of different classes of chemical neurotransmitters.

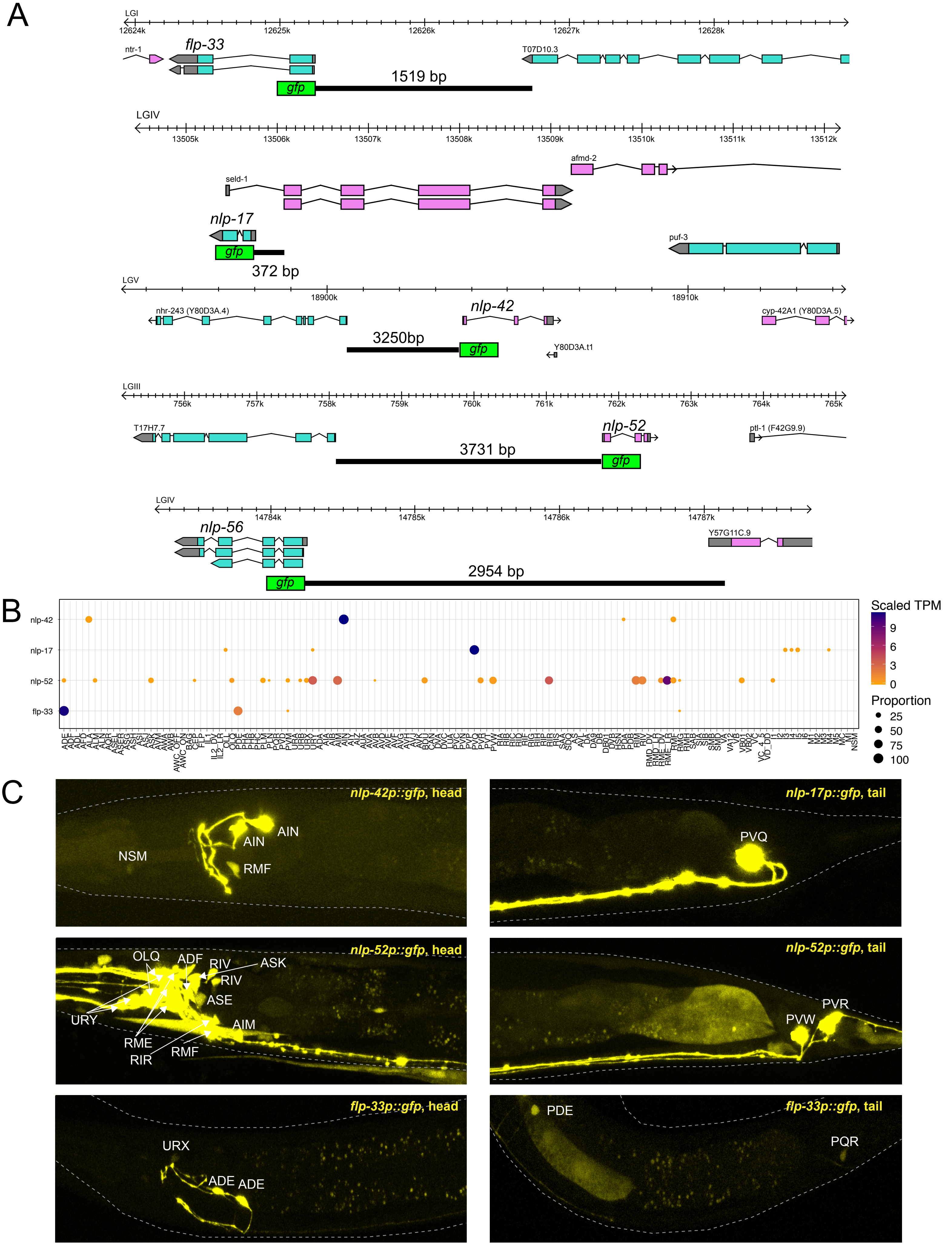

A
LGI
12624k
12625k
12626k
12627k
12628k
ntr-1
flp-33
T07D10.3
gfp
1519 bp
LGIV
13505k
13506k
13507k
13508k
13509k
13510k
13511k
13512k
afmd-2
seld-1
nlp-17
puf-3
gfp
372 bp
LGV
18900k
18910k
nhr-243 (Y80D3A.4)
nlp-42
cyp-42A1 (Y80D3A.5)
3250bp
gfp
Y80D3A.t1
LGIII
756k
757k
758k
759k
760k
761k
762k
763k
764k
765k
T17H7.7
nlp-52
ptl-1 (F42G9.9)
3731 bp
gfp
LGIV
14784k
14785k
14786k
14787k
nlp-56
Y57G11C.9
2954 bp
gfp
B
nlp-42
nlp-17
nlp-52
flp-33
Scaled TPM
9
6
3
0
Proportion
25
50
75
100
C
nlp-42p::gfp, head
NSM
AIN
AIN
RMF
nlp-17p::gfp, tail
PVQ
nlp-52p::gfp, head
OLQ
ADF
RIV
ASK
RIV
ASE
URY
AIM
RME
RIR
RMF
nlp-52p::gfp, tail
PVR
PVW
flp-33p::gfp, head
URX
ADE
ADE
flp-33p::gfp, tail
PDE
PQR

**Figure S8. Neuropeptide gene reporter strains validate single cell RNA-Seq data**. A) Diagrams depicting upstream regions (black lines) in transcriptional GFP reporter genes. B) Heatmap showing single cell RNA-Seq expression of four selected neuropeptides (*nlp-42*, *nlp-17*, *nlp-52*, *flp-33*) and (C) corresponding confocal micrographs. Neuron types expressing each neuropeptide reporter were determined by co-localization with NeuroPAL markers (not shown). *nlp-42p::gfp* is robustly expressed in AIN and RMF with faint expression in NSM and PHB. *nlp-17* expression at threshold 2 was 480x stronger in PVQ than for six additional neuron types and the *nlp-17p::gfp* reporter was selectively detected in PVQ. GFP reporters for *nlp-52* and *flp-33* (C) are largely congruent with single-cell RNA-Seq results (B). Related to Figure 4.

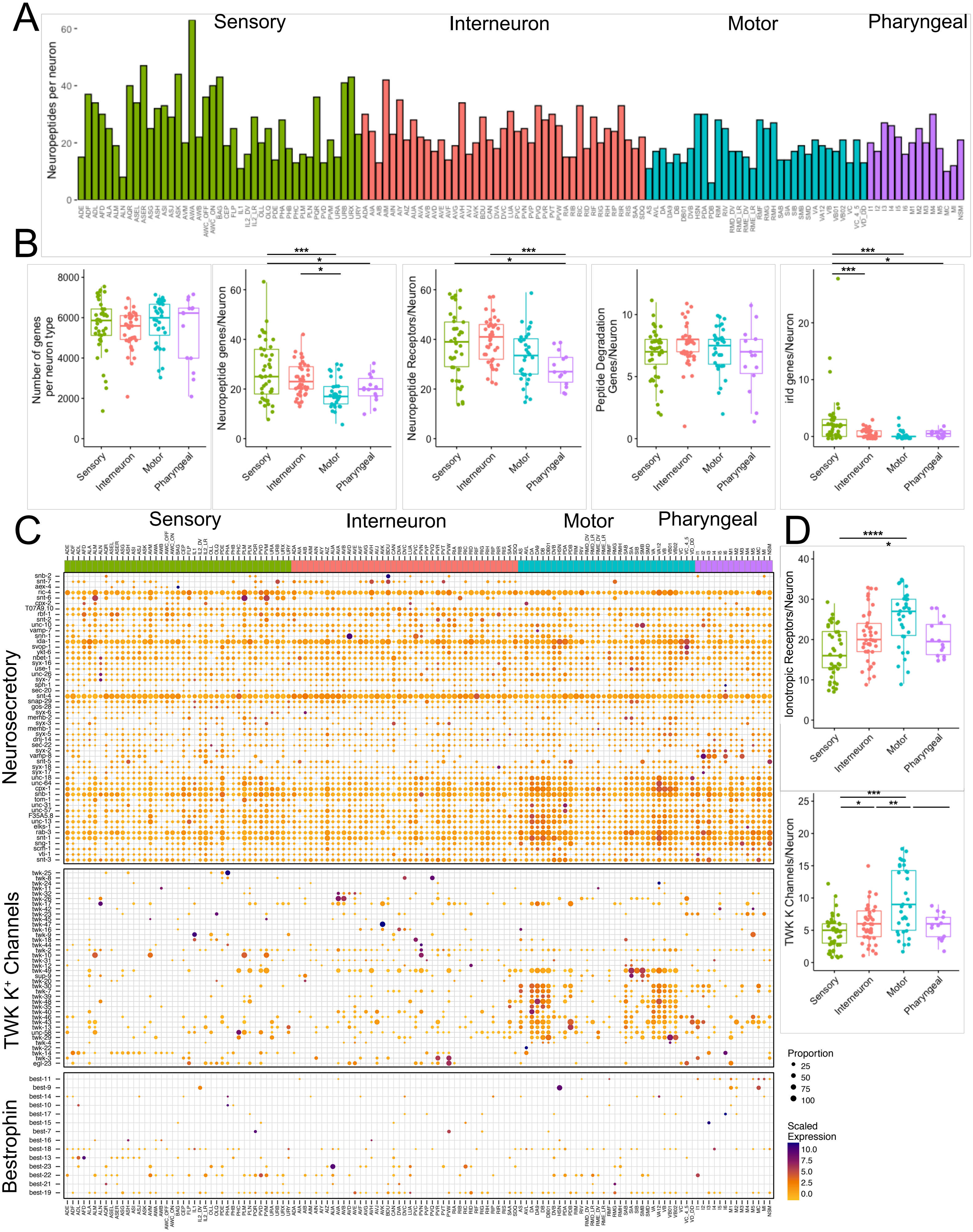

A
Sensory
Interneuron
Motor
Pharyngeal
Neuropeptides per neuron
B
Number of genes per neuron type
Neuropeptide genes/Neuron
Neuropeptide Receptors/Neuron
Peptide Degradation Genes/Neuron
irld genes/Neuron
Sensory
Interneuron
Motor
Pharyngeal
C
Sensory
Interneuron
Motor
Pharyngeal
Neurosecretory
TWK K⁺ Channels
Bestrophin
D
Ionotropic Receptors/Neuron
TWK K Channels/Neuron
Proportion
25
50
75
100
Scaled Expression
10.0
7.5
5.0
2.5
0.0

**Figure S9. Expression of selected gene families for neuronal signaling and ion channel function in different neuron classes.** A) Bar graph showing expression of neuropeptide encoding genes at threshold 2 for neurons grouped by modality (Sensory, Interneuron, Motor, Pharyngeal). B) Number of genes (threshold 2) expressed in each neuron type grouped by modality for all genes and neuropeptide signaling genes (neuropeptide-encoding genes, neuropeptide receptors, peptide degradation genes, *irld* genes). Boxes are interquartile ranges. Statistical tests: ANOVA, with Tukey post-hoc comparisons for neuropeptide receptors, Kruskal-Wallis test for other genes. $^*p < 0.05$, $^{**}p < 0.01$, $^{***}p < 0.001$. C) Heatmaps showing expression of genes for neurosecretory components, TWK $K^+$ channels, and Bestrophin anion channels across 128 neuron types (columns) grouped by neuron modality. Protein subfamilies are clustered separately. D) Number of genes (threshold 2) expressed in each neuron type grouped by modality Ligand-Gated Ion Channels (LGICs) and TWK $K^+$ Channels. Boxes are interquartile ranges. Statistical tests: Kruskal-Wallis. $^*p < 0.05$, $^{**}p < 0.01$, $^{***}p < 0.001$, $^{****}p < 0.0001$. Related to Figure 4.

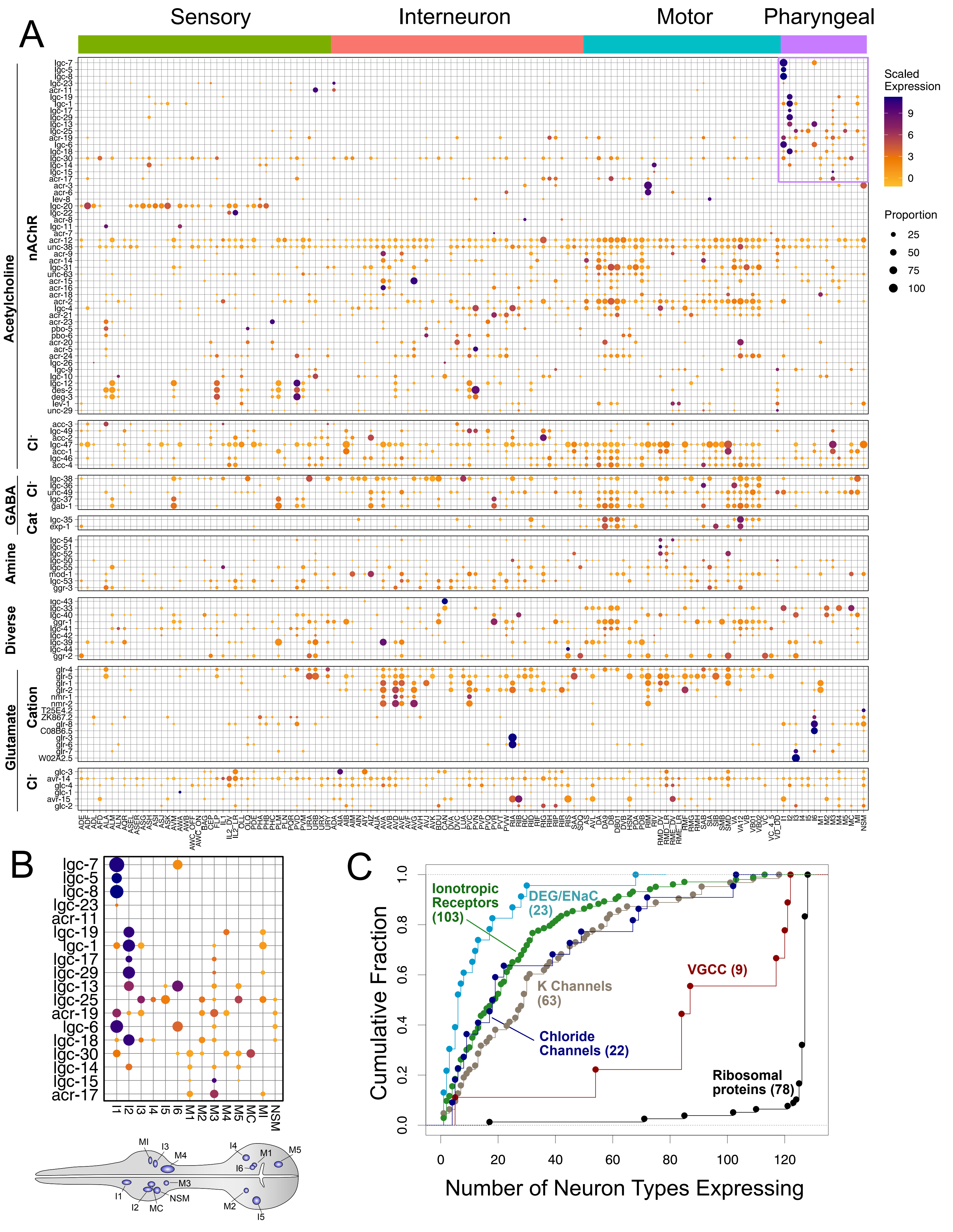
A
Sensory
Interneuron
Motor
Pharyngeal
Acetylcholine
nAChR
lgc-7
lgc-5
lgc-8
lgc-23
acr-11
lgc-19
lgc-1
lgc-17
lgc-29
lgc-13
lgc-25
acr-19
lgc-6
lgc-18
lgc-30
lgc-14
lgc-15
acr-17
acr-3
acr-6
lev-8
lgc-20
lgc-22
acr-8
lgc-11
acr-7
acr-12
unc-38
acr-9
acr-14
lgc-31
unc-63
acr-15
acr-16
acr-18
acr-2
lgc-4
acr-21
acr-23
pbo-5
pbo-6
acr-20
acr-5
acr-24
lgc-26
lgc-9
lgc-10
lgc-12
des-2
deg-3
lev-1
unc-29
Cl-
acc-3
lgc-49
acc-2
lgc-47
acc-1
lgc-46
acc-4
GABA
Cl-
lgc-38
lgc-36
unc-49
lgc-37
gab-1
Cat
lgc-35
exp-1
Amine
lgc-54
lgc-51
lgc-52
lgc-50
lgc-55
mod-1
lgc-53
ggr-3
Diverse
lgc-43
lgc-33
lgc-40
ggr-1
lgc-41
lgc-42
lgc-39
lgc-44
ggr-2
Glutamate
Cation
glr-4
glr-5
glr-1
glr-2
nmr-1
nmr-2
T25E4.2
ZK867.2
glr-8
C08B6.5
glr-3
glr-6
glr-7
W02A2.5
Cl-
glc-3
avr-14
glc-4
glc-1
avr-15
glc-2
ADE ADF ADL AFD ALA ALM ALN AQR ASEL ASER ASG ASH ASI ASJ ASK AVM AWA AWB AWC_OFF AWC_ON BAG CEP FLP IL1 IL2_DV IL2_LR OLL OLQ PDE PHA PHB PHC PLM PLN PQR PVD PVM URA URB URX URY ADA AIA AIB AIM AIN AIY AIZ AUA AVA AVB AVD AVE AVF AVG AVH AVJ AVK BDU CAN DVA DVC LUA PVC PVN PVP PVQ PVR PVT PVW RIA RIB RIC RID RIF RIG RIH RIP RIR RIS SAA SDQ AS AVL DA DA9 DB DB01 DVB HSN PDA PDB RIM RIV RMD_DV RMD_LR RME_DV RME_LR RMF RMG RMH SAB SIA SIB SMB SMD VA VA12 VB VB01 VB02 VC VC_4_5 VD_DD I1 I2 I3 I4 I5 I6 M1 M2 M3 M4 M5 MC MI NSM
Scaled Expression
9
6
3
0
Proportion
25
50
75
100
B
lgc-7
lgc-5
lgc-8
lgc-23
acr-11
lgc-19
lgc-1
lgc-17
lgc-29
lgc-13
lgc-25
acr-19
lgc-6
lgc-18
lgc-30
lgc-14
lgc-15
acr-17
I1 I2 I3 I4 I5 I6 M1 M2 M3 M4 M5 MC MI NSM
MI
I3
M4
I4
M1
M5
I6
I1
M3
NSM
I2
MC
M2
I5
C
Cumulative Fraction
0.0
0.2
0.4
0.6
0.8
1.0
Ionotropic Receptors (103)
DEG/ENaC (23)
K Channels (63)
Chloride Channels (22)
VGCC (9)
Ribosomal proteins (78)
0
20
40
60
80
100
120
Number of Neuron Types Expressing

**Figure S10. Expression of ligand-gated ion channels.** A) Heatmaps of ionotropic receptors including 89 Cys-loop ligand-gated ion channel (LGIC) genes grouped by neurotransmitter (Acetylcholine, GABA, Amine, Glutamate) and by ion selectivity ($Cl^-$ vs cation) and 14 glutamate-gated cation channels across 128 neuron types (columns). B) (Top) Inset of selected region of heatmap in A showing 18 Cys-loop LGICs enriched in pharyngeal neurons (bottom). C) Cumulative distribution plot of number of neuron types expressing genes encoding ionotropic receptors (Cys-loop LGICs and glutamate-gated cation channels), DEG/ENaC channels, potassium (K) channels, chloride channels, VGCC (Voltage-Gated Calcium Channels) and ribosomal proteins (see also Figure 4A). Related to Figure 4.

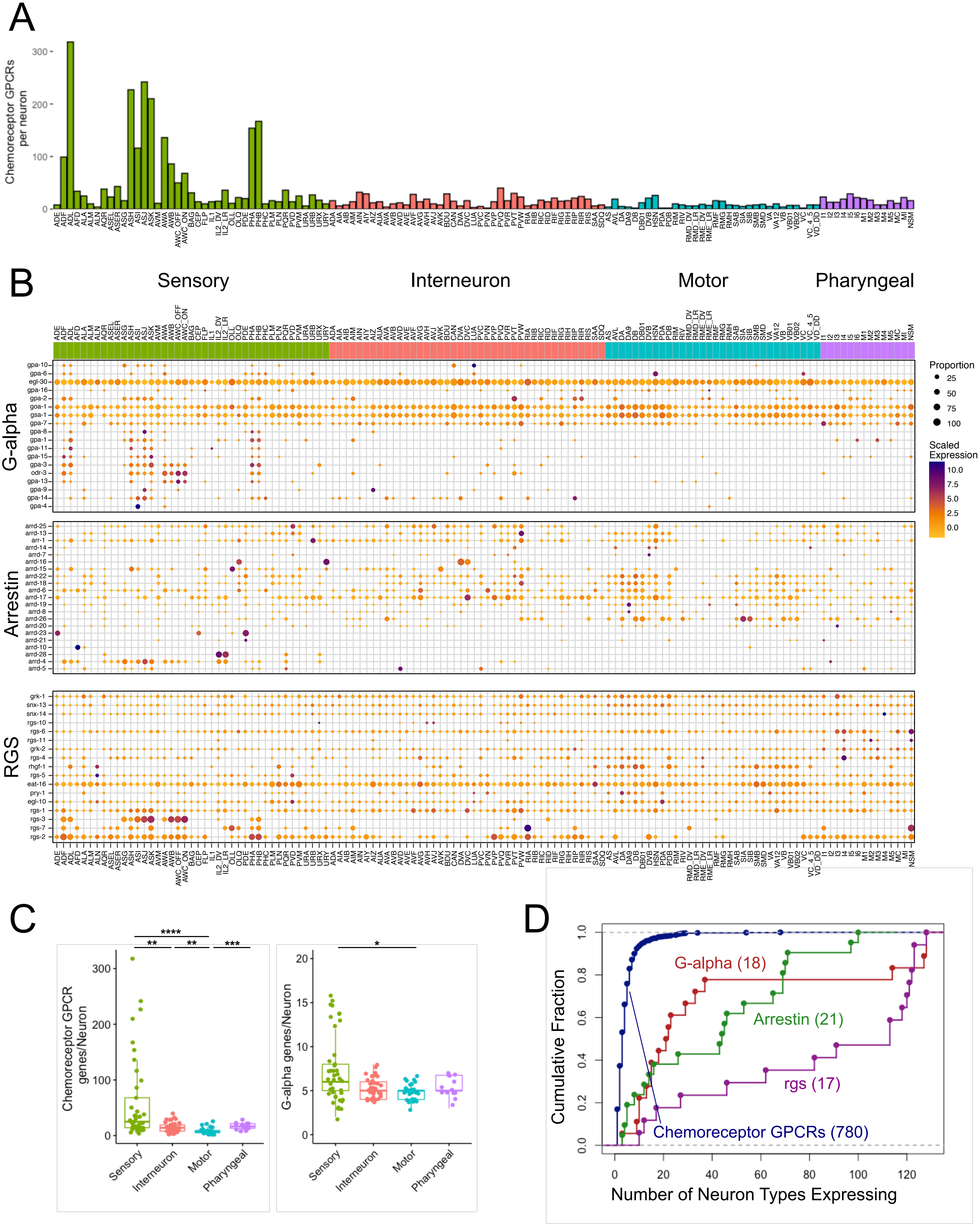

A
Chemoreceptor GPCRs per neuron
B
Sensory
Interneuron
Motor
Pharyngeal
G-alpha
Arrestin
RGS
Proportion
25
50
75
100
Scaled Expression
10.0
7.5
5.0
2.5
0.0
gpa-10
gpa-6
egl-30
gpa-16
gpa-2
goa-1
gsa-1
gpa-7
gpa-8
gpa-1
gpa-11
gpa-15
gpa-3
odr-3
gpa-13
gpa-9
gpa-14
gpa-4
arrd-25
arrd-13
arr-1
arrd-14
arrd-7
arrd-16
arrd-15
arrd-22
arrd-18
arrd-6
arrd-17
arrd-19
arrd-8
arrd-26
arrd-20
arrd-23
arrd-21
arrd-10
arrd-28
arrd-4
arrd-5
grk-1
snx-13
snx-14
rgs-10
rgs-6
rgs-11
grk-2
rgs-4
rhgf-1
rgs-5
eat-16
pry-1
egl-10
rgs-1
rgs-3
rgs-7
rgs-2
C
Chemoreceptor GPCR genes/Neuron
G-alpha genes/Neuron
Sensory
Interneuron
Motor
Pharyngeal
D
Cumulative Fraction
Number of Neuron Types Expressing
G-alpha (18)
Arrestin (21)
rgs (17)
Chemoreceptor GPCRs (780)

**Figure S11. Expression of chemoreceptor GPCRs and downstream signaling pathways.** A) Bar graph showing expression of chemoreceptor GPCRs across all neuron types. All neurons express at least two chemoreceptor GPCRs. Note enrichment in a subset of ciliated amphid/phasmid sensory neurons. B) Heatmaps showing expression of genes for G-alpha, arrestin and RGS families. Note that neurons expressing the highest numbers of chemoreceptors (Panel A, ADL, ASH, ASJ, ASK, AWA, PHA, PHB) also abundantly express G-alpha and RGS proteins. C) The number of chemoreceptor GPCRs (left) and G-alpha genes (right) grouped by neuron modality (Sensory, Interneuron, Motor, Pharyngeal). Boxes are interquartile ranges. Kruskal-wallis test, *$p < 0.05$, **$p < 0.01$, ***$p < 0.001$, ****$p < 0.0001$. D) Cumulative distribution plot of chemoreceptor GPCRs, G-alpha, arrestin and RGS families.

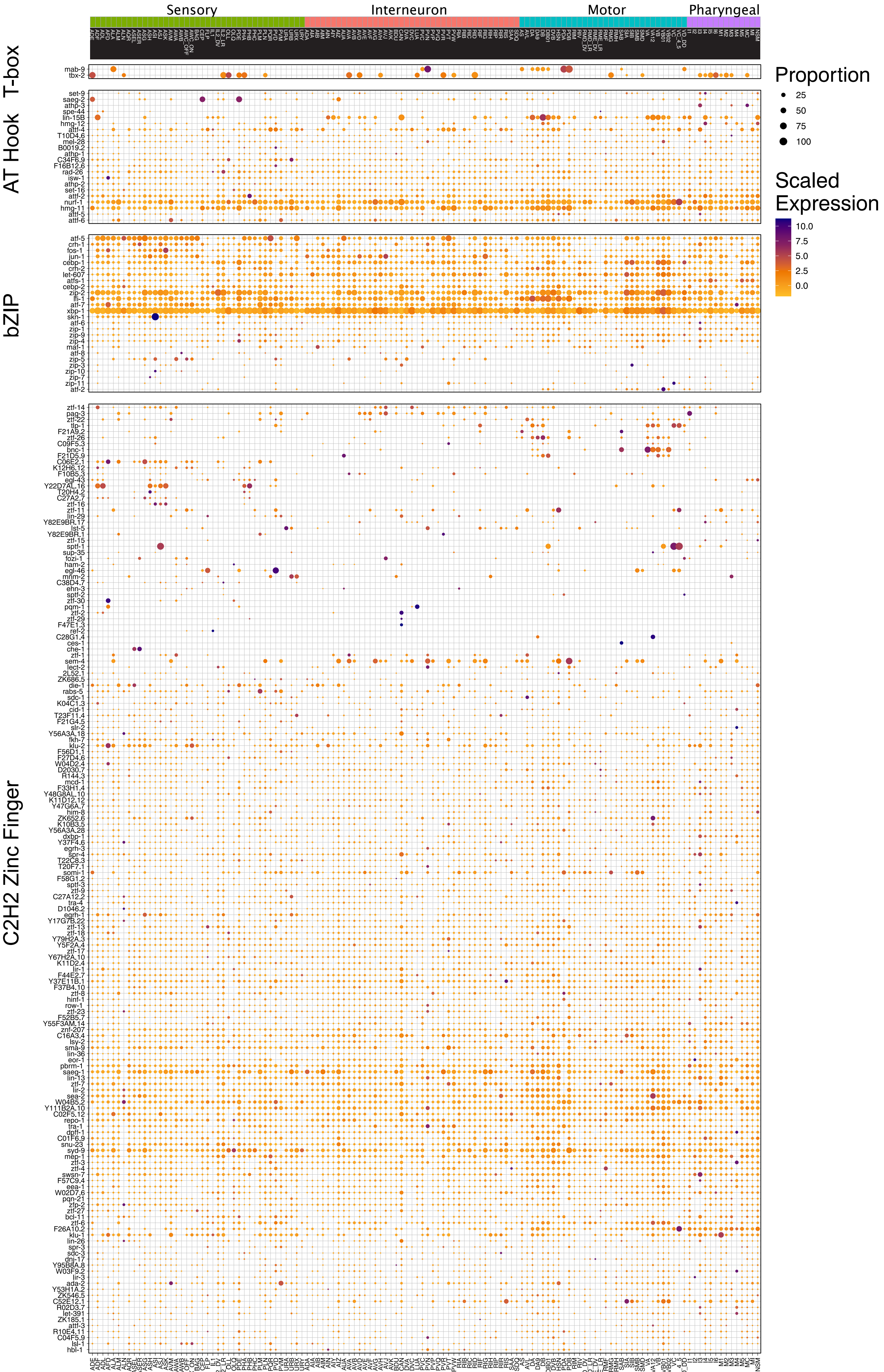
Sensory
Interneuron
Motor
Pharyngeal
T-box
AT Hook
bZIP
C2H2 Zinc Finger
Proportion
25
50
75
100
Scaled Expression
10.0
7.5
5.0
2.5
0.0

**Figure S12. Expression of transcription factor subfamilies.** Heatmaps showing expression of the T-box, AT hook, bZIP and C2H2 zinc-finger (ZF) transcription factor families. Neurons are grouped according to modality (Sensory, Interneuron, Motor, Pharyngeal).and genes are clustered within each family. Related to Figure 5.

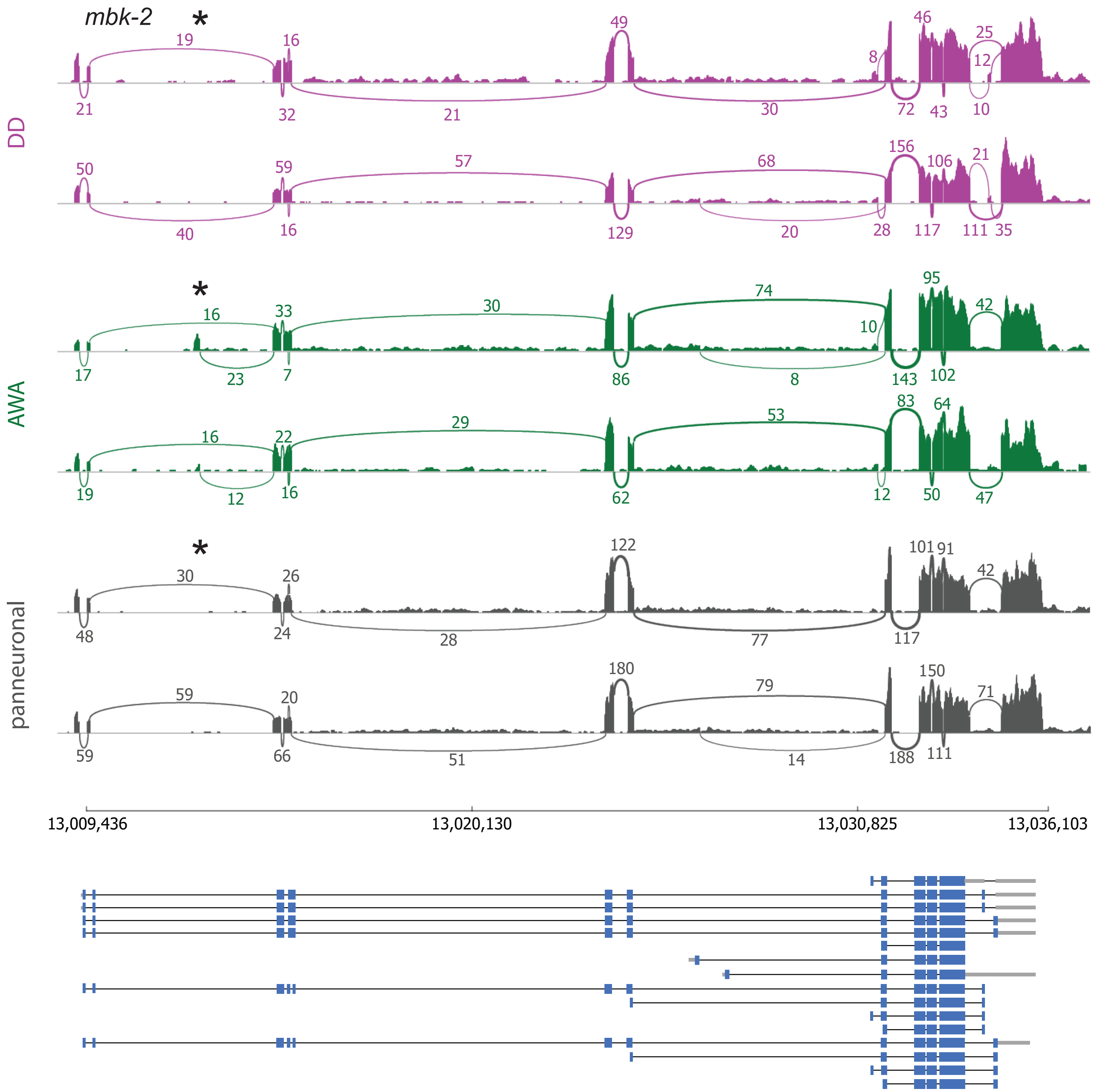

mbk-2
*
DD
19
16
49
46
25
12
8
21
32
21
30
72
43
10
50
59
57
68
156
106
21
40
16
129
20
28
117
111
35
AWA
*
16
33
30
74
95
42
10
17
23
7
86
8
143
102
16
22
29
53
83
64
19
12
16
62
12
50
47
panneuronal
*
30
26
122
101
91
42
48
24
28
77
117
59
20
180
79
150
71
59
66
51
14
188
111
13,009,436
13,020,130
13,030,825
13,036,103

**Figure S13. Detection of neuron-specific splicing of a novel exon by bulk RNA-Seq.** Sashimi plot representing the splicing pattern of the gene *mbk-2* in neurons DD, AWA and pan-neuronal samples. Two representative samples are represented for each condition. The (*) indicates an alternative 5' exon used exclusively in AWA and undetectable in pan-neuronal samples. Related to Figure 6.

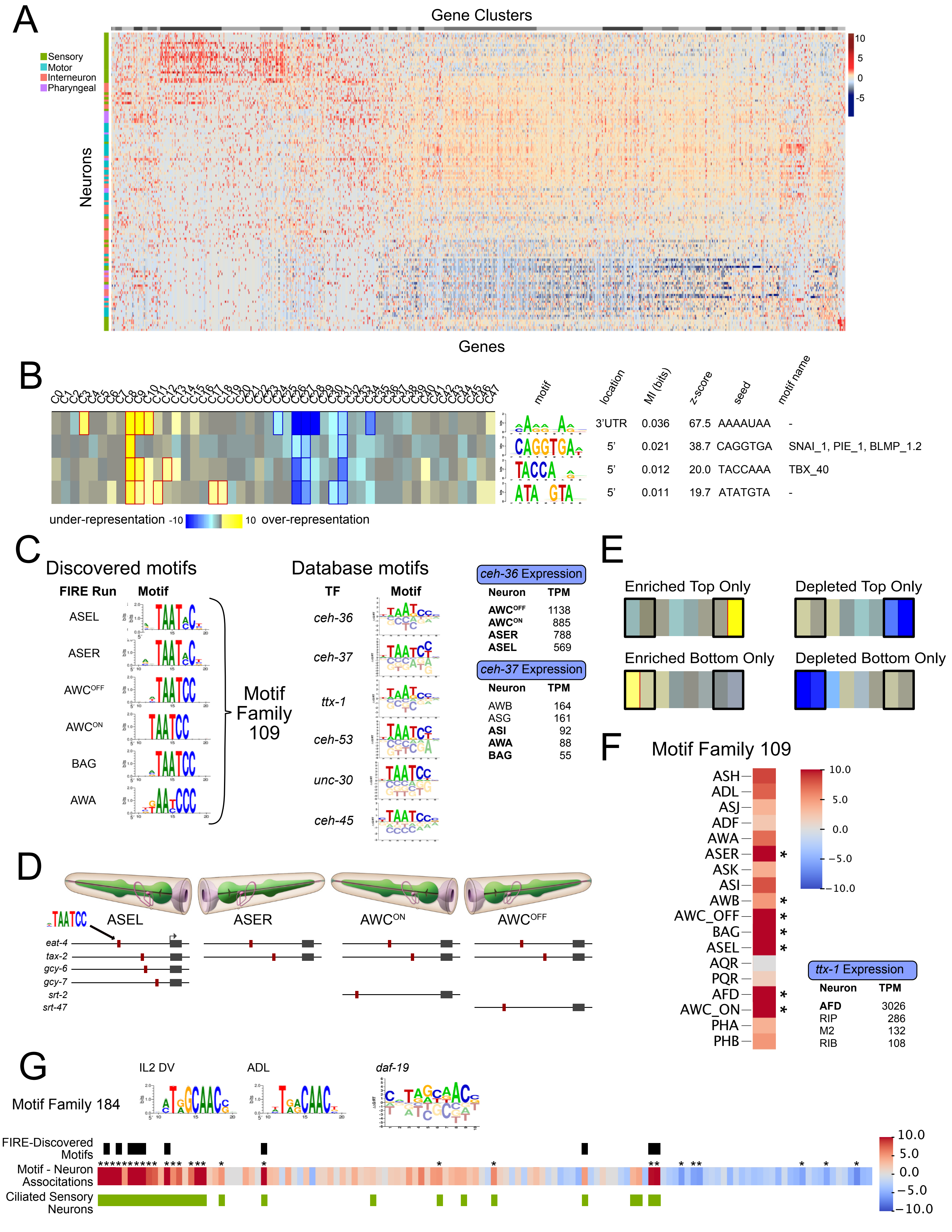
A
Gene Clusters
Sensory
Motor
Interneuron
Pharyngeal
Neurons
Genes
10
5
0
-5
B
motif
location
MI (bits)
z-score
seed
motif name
3'UTR 0.036 67.5 AAAAUAA -
5' 0.021 38.7 CAGGTGA SNAI_1, PIE_1, BLMP_1.2
5' 0.012 20.0 TACCAAA TBX_40
5' 0.011 19.7 ATATGTA -
under-representation -10
10 over-representation
C
Discovered motifs
FIRE Run
Motif
ASEL
ASER
AWCOFF
AWCON
BAG
AWA
Motif Family 109
Database motifs
TF
Motif
ceh-36
ceh-37
ttx-1
ceh-53
unc-30
ceh-45
ceh-36 Expression
Neuron TPM
AWCOFF 1138
AWCON 885
ASER 788
ASEL 569
ceh-37 Expression
Neuron TPM
AWB 164
ASG 161
ASI 92
AWA 88
BAG 55
D
ASEL
ASER
AWCON
AWCOFF
eat-4
tax-2
gcy-6
gcy-7
srt-2
srt-47
E
Enriched Top Only
Depleted Top Only
Enriched Bottom Only
Depleted Bottom Only
F
Motif Family 109
ASH
ADL
ASJ
ADF
AWA
ASER
ASK
ASI
AWB
AWC_OFF
BAG
ASEL
AQR
PQR
AFD
AWC_ON
PHA
PHB
10.0
5.0
0.0
−5.0
−10.0
ttx-1 Expression
Neuron TPM
AFD 3026
RIP 286
M2 132
RIB 108
G
IL2 DV
ADL
daf-19
Motif Family 184
FIRE-Discovered Motifs
Motif - Neuron Associations
Ciliated Sensory Neurons
10.0
5.0
0.0
−5.0
−10.0

**Figure S14. FIRE discovered known regulatory motifs in individual neurons.** A) Heatmap of scaled gene expression (z-scores for each gene in each neuron). Both genes (columns) and neurons (rows) are ordered by similarity. X-axis color bar (top) denotes 48 gene clusters. Row color bar (left) indicates neuron modality (Sensory, Motor, Interneuron, Pharyngeal). B) Selected results from FIRE analysis of 48 gene clusters. C) Left: Motif family 109 includes highly similar motifs discovered in independent FIRE analyses from individual neurons. Center: Motifs from the CIS-BP database for six different K50 homeodomain transcription factors (CEH-36, CEH-37, TTX-1, CEH-53, UNC-30, CEH-45) match motif family 109. Right) Expression (TPM or transcripts per million) of two transcription factors (*ceh-36, ceh-37*) in our scRNA-Seq data corresponds to the neurons in which the cognate motif was discovered. D) The TAATCC motif (red boxes) in the 5' regions of genes enriched in ASEL, ASER, $AWC^{ON}$ or $AWC^{OFF}$, including genes expressed in all four neurons (top rows) as well as genes expressed in subsets of these neurons. Expression of *eat-4*, *tax-2*, *gcy-6*, *gcy-7* and *srt-2* is *ceh-36*-dependent in these neurons. E) Schematic showing the z-score bins tested for relative enrichment for motif-neuron associations (See Methods). F) Motif-neuron associations (from FIRE Main Figure B for motif family 109 in a subset of neurons) showing correspondence to expression of K50 homeodomain proteins (black arrows). Colors indicate -log10(p-value) for positive associations and log10(p-value) for negative associations. Asterisks denote significant associations (pval < 1e-5, log fold change > 0.5). G) Top: Two similar motifs discovered in FIRE runs of individual neurons IL2_DV and ADL are members of motif family 184 that matches the DAF-19 motif from CIS-BP. Other members of the DAF-19 motif family were discovered in FIRE analysis of 8 additional individual neurons (black boxes). Bottom: Motif family 184 showed significant positive associations (asterisks) with 22 ciliated sensory neuron types (green boxes) and significant negative associations with some motor neurons (blue boxes with asterisks). Related to Figure 8.

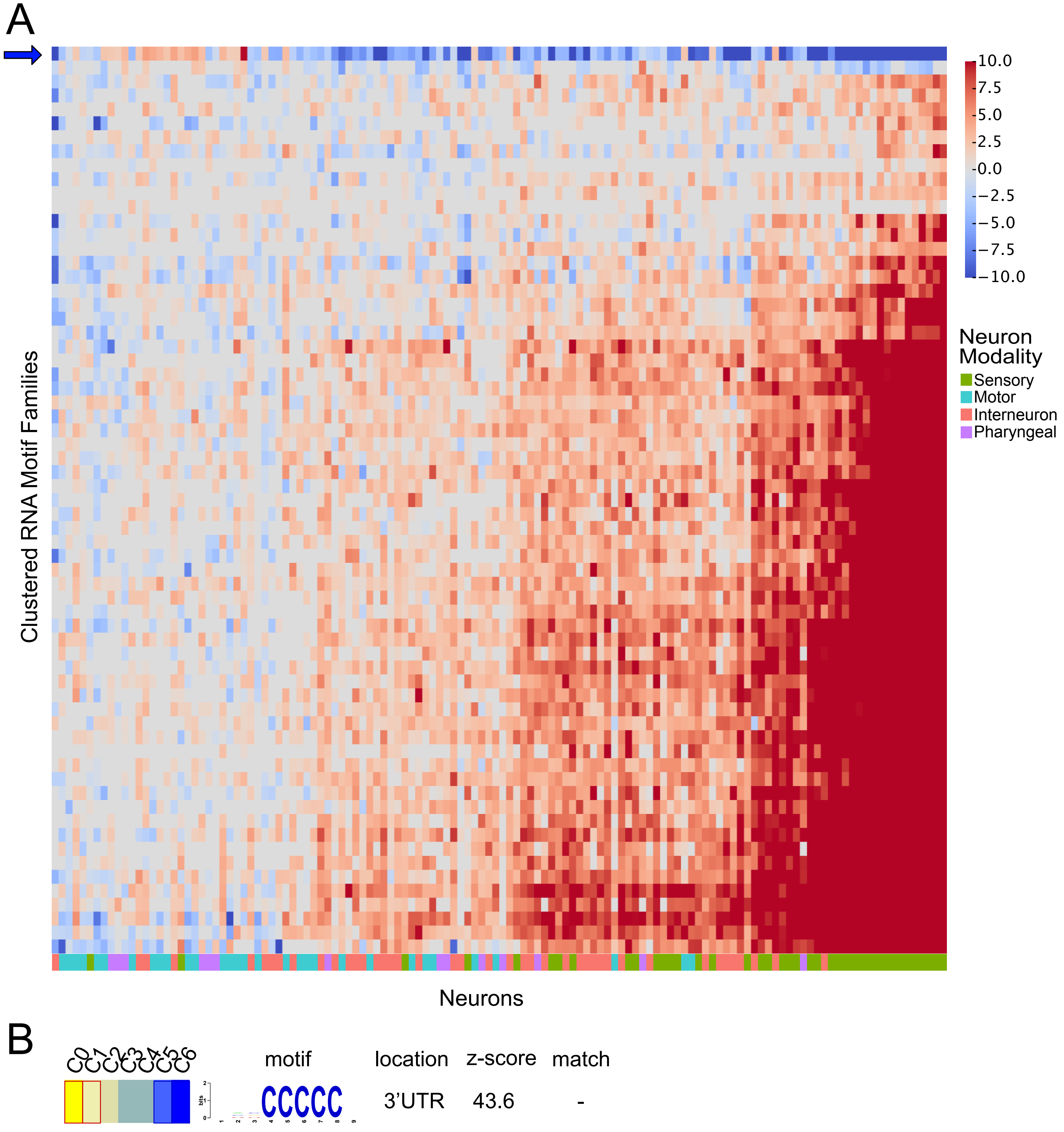
A
Clustered RNA Motif Families
Neurons
10.0
7.5
5.0
2.5
0.0
−2.5
−5.0
−7.5
−10.0
Neuron Modality
Sensory
Motor
Interneuron
Pharyngeal
B
C0
C1
C2
C3
C4
C5
C6
motif
location
z-score
match
CCCCC
3'UTR
43.6
-

**Figure S15. RNA motif families are positively associated with a majority of neuron types.** Heatmap of log-transformed p-values showing motif-neuron associations for RNA motif families (columns) across all neuron types (rows). Note the strong positive signal in sensory neurons, and the one motif family with largely negative associations with neurons (blue arrow). B) Representative RNA motif (from ADL FIRE run) of family 23 showing a poly-C RNA sequence in the 3' UTR. Related to Figure 8.

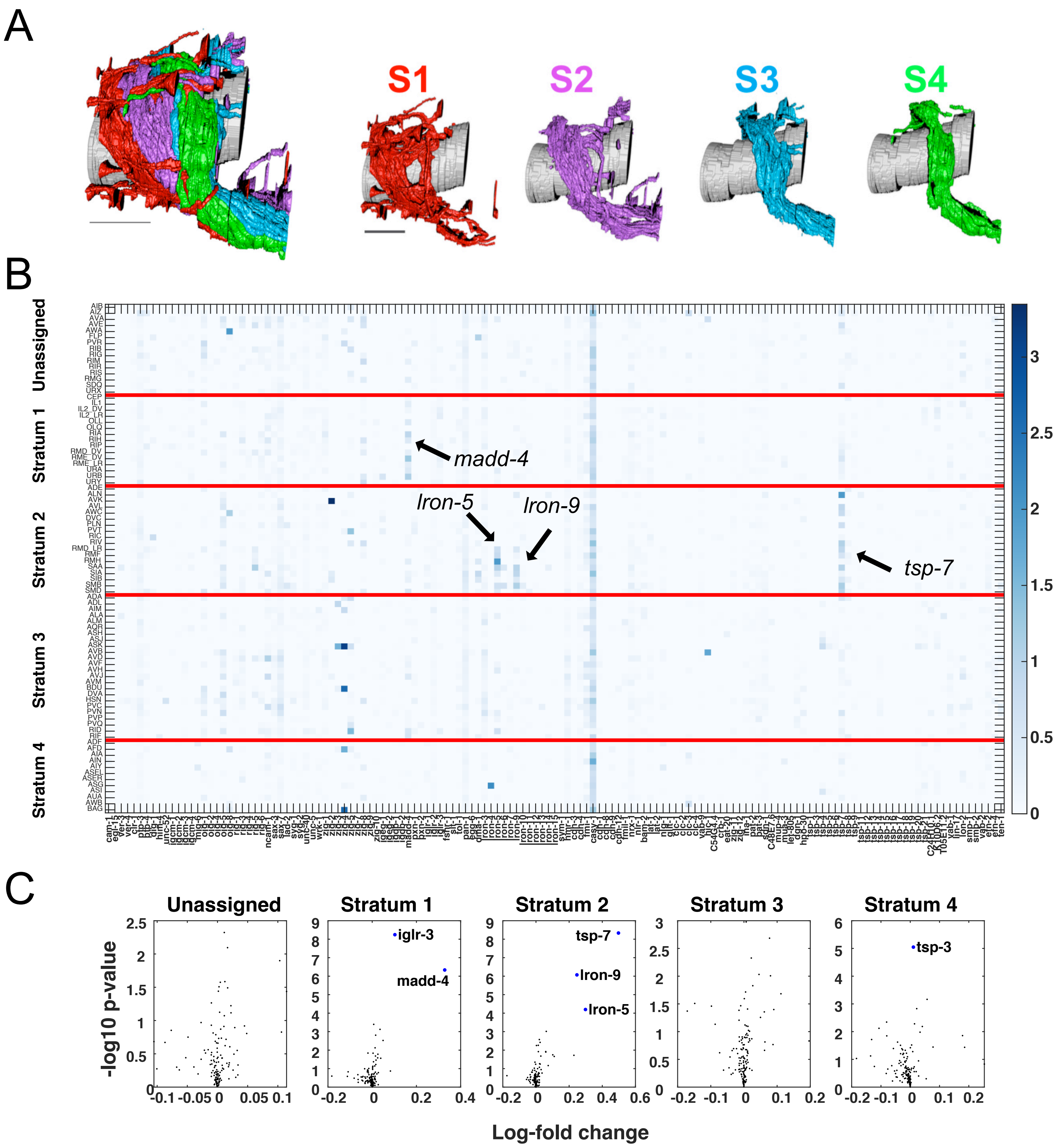
A
S1
S2
S3
S4
B
Unassigned
Stratum 1
Stratum 2
Stratum 3
Stratum 4
madd-4
lron-5
lron-9
tsp-7
3
2.5
2
1.5
1
0.5
0
C
Unassigned
Stratum 1
Stratum 2
Stratum 3
Stratum 4
iglr-3
madd-4
tsp-7
lron-9
lron-5
tsp-3
-log10 p-value
Log-fold change

**Figure S16. Differential expression of CAMs between strata.** A) Cartoon representation of nerve ring strata (Moyle et al., 2020). Figure courtesy of Daniel Colón-Ramos. B) Heatmap of CAM expression (columns) in neurons grouped by strata (rows). Black arrows indicate four significantly enriched CAMs with the highest log fold changes between strata. Colors indicate log-transformed expression values. Red lines separate strata. C) Volcano plots showing log-fold change (x-axis) by -log10 p-value (y-axis) for each CAM in the neurons in each stratum compared to all other neurons. The six labeled genes were significantly enriched in the strata shown. Related to Figure 9.

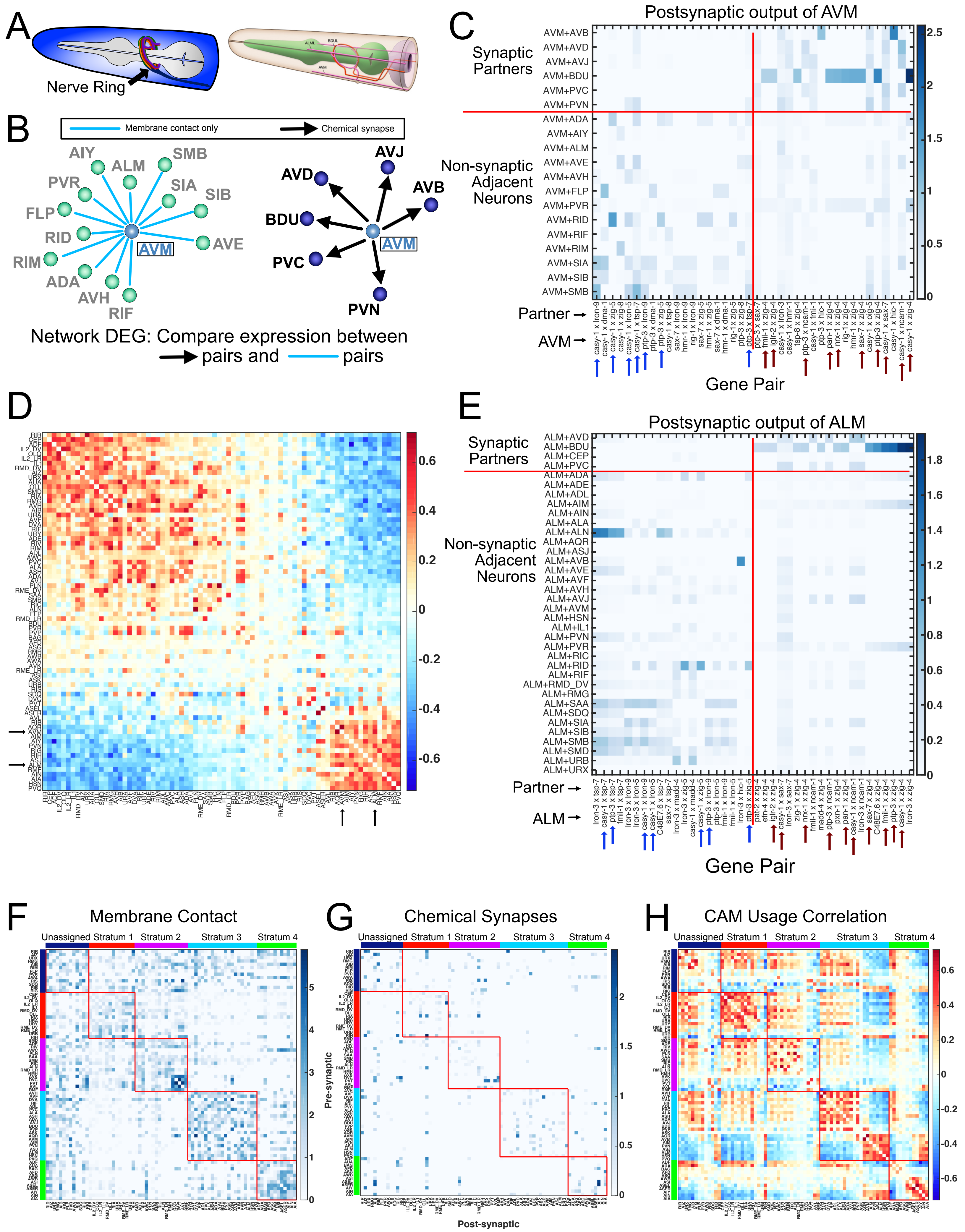
A
Nerve Ring
B
Membrane contact only
Chemical synapse
AIY
ALM
SMB
PVR
SIA
SIB
FLP
RID
AVE
RIM
ADA
AVH
RIF
AVM
AVD
AVJ
AVB
BDU
PVC
PVN
Network DEG: Compare expression between
pairs and pairs
C
Postsynaptic output of AVM
Synaptic Partners
Non-synaptic Adjacent Neurons
AVM+AVB
AVM+AVD
AVM+AVJ
AVM+BDU
AVM+PVC
AVM+PVN
AVM+ADA
AVM+AIY
AVM+ALM
AVM+AVE
AVM+AVH
AVM+FLP
AVM+PVR
AVM+RID
AVM+RIF
AVM+RIM
AVM+SIA
AVM+SIB
AVM+SMB
Partner →
AVM →
Gene Pair
D
E
Postsynaptic output of ALM
Synaptic Partners
Non-synaptic Adjacent Neurons
ALM+AVD
ALM+BDU
ALM+CEP
ALM+PVC
ALM+ADA
ALM+ADE
ALM+ADL
ALM+AIM
ALM+AIN
ALM+ALA
ALM+ALN
ALM+AQR
ALM+ASJ
ALM+AVB
ALM+AVE
ALM+AVF
ALM+AVH
ALM+AVJ
ALM+AVM
ALM+HSN
ALM+IL1
ALM+PVN
ALM+PVR
ALM+RIC
ALM+RID
ALM+RIF
ALM+RMD_DV
ALM+RMG
ALM+SAA
ALM+SDQ
ALM+SIA
ALM+SIB
ALM+SMB
ALM+SMD
ALM+URB
ALM+URX
Partner →
ALM →
Gene Pair
F
Membrane Contact
G
Chemical Synapses
H
CAM Usage Correlation
Unassigned
Stratum 1
Stratum 2
Stratum 3
Stratum 4
Pre-synaptic
Post-synaptic

**Figure S17. Differential expression of cell adhesion molecules among neurons and their postsynaptic partners.** A) Left: Cartoon representation of the *C. elegans* nerve ring. Right: Subset of nerve ring neurites from the touch neurons AVM, ALM and their synaptic partner BDU. Courtesy of WormAtlas. B) Diagram of neurons receiving chemical postsynaptic output from AVM (right) and neurons with membrane contact but no synapses with AVM (left). Differential cell adhesion molecule (CAM) gene expression was determined between AVM + synaptic partners and AVM + non-synaptic adjacent neurons. C) Heatmap of the 20 gene pairs with highest log fold change in AVM and its postsynaptic outputs compared to AVM and adjacent neurons (right of vertical red line). The 20 gene pairs with highest log fold change in AVM and adjacent neurons compared to AVM and synaptic partners are shown to the left of vertical red line. Blue and dark red arrows indicate gene pairs common among analyses of AVM and ALM (panel D). The first gene listed is expressed in AVM, the second gene is expressed in the synaptic or adjacent partner. D) Correlation matrix showing the relationship of CAM usage across all neurons in the nerve ring (84 neuron types). Note the similarities in CAM usage among the neurons in the lower right-hand corner. The black arrows indicate AVM and ALM (correlation 0.607). E) Heatmap as in C, for ALM. 10 of the top 20 gene pairs (dark red arrows) higher in ALM and its synaptic partners compared to ALM and adjacent neurons are common with the AVM analysis in B. On the left, 7 of the top 20 genes pairs higher in ALM/adjacent are common with AVM (blue arrows). F) The membrane adjacency matrix was grouped according to nerve ring strata as defined in Moyle, et al. Within each stratum, neurons were ordered according to their CAM usage correlations (see panel H). Membrane contact is denser within strata than between strata, but not for the unassigned neurons which generally contact neurons in other strata. Colored bars indicate neuron strata assignments. Each stratum is outlined by a thin red box. G) The same ordering as in F was imposed upon the chemical connectome revealing that most synapses are detected between neurons within the same stratum. H) The CAM usage correlation matrix (as in D) was grouped by strata, then sorted by similarity (using multidimensional scaling) within each stratum. Neurons within stratum 1 are similar to each other. Strata 2, 3, and 4 all show two distinct groups of neurons based on CAM usage at postsynaptic outputs. Related to Figure 9.

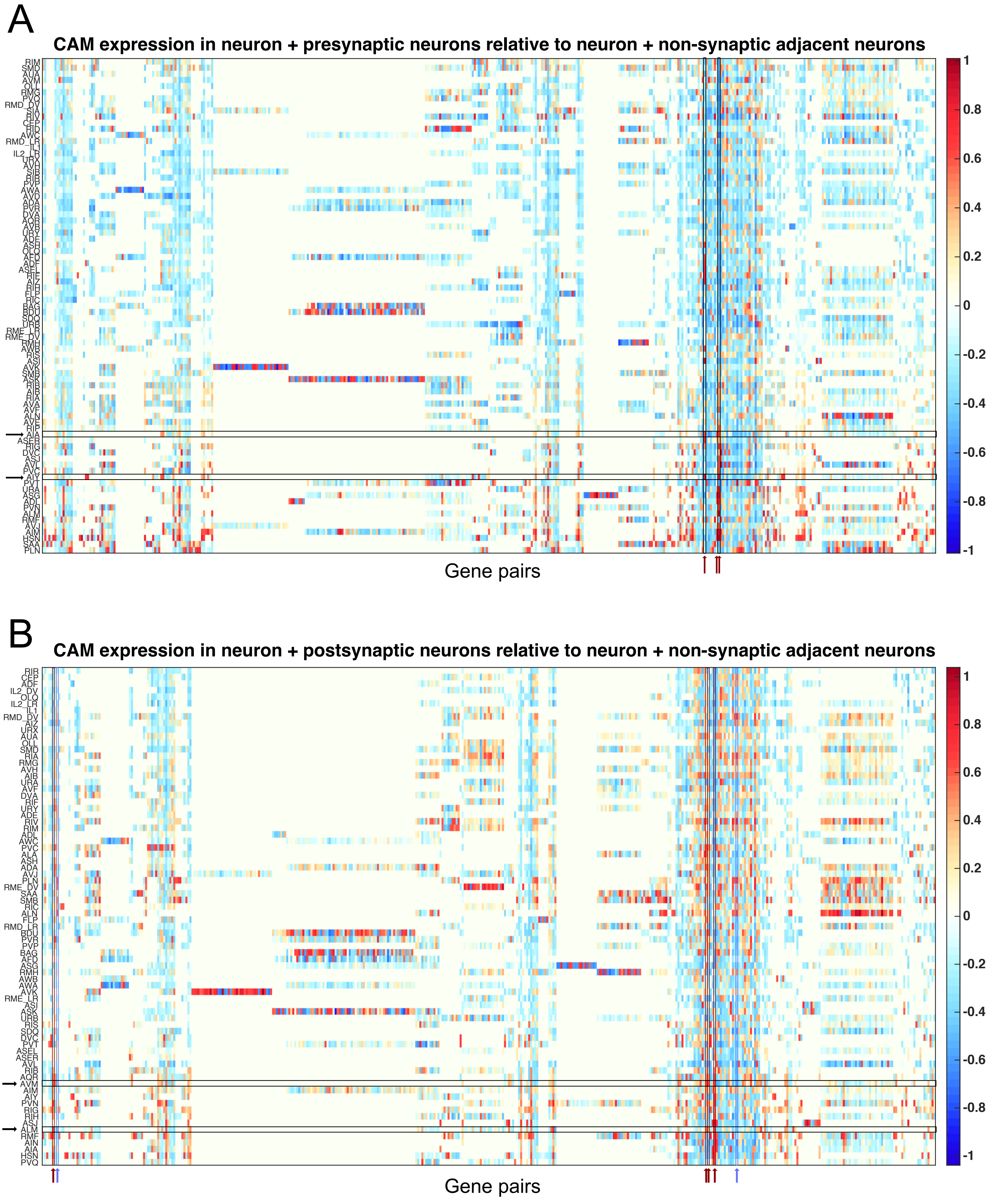
A
CAM expression in neuron + presynaptic neurons relative to neuron + non-synaptic adjacent neurons
Gene pairs
1
0.8
0.6
0.4
0.2
0
-0.2
-0.4
-0.6
-0.8
-1
B
CAM expression in neuron + postsynaptic neurons relative to neuron + non-synaptic adjacent neurons
Gene pairs
1
0.8
0.6
0.4
0.2
0
-0.2
-0.4
-0.6
-0.8
-1

**Figure S18. CAM usage across nerve ring neurons.** A) Heatmap of CAM usage in presynaptic connections for each neuron in the nerve ring. Neurons are rows, gene pairs are columns. Colors indicate the log of the t-statistic comparing gene pair expression in each neuron + presynaptic partners vs each neuron + non-synaptic adjacent neurons. Positive values (orange and red) indicate higher expression in synaptically-connected neurons relative to adjacent neurons. Negative values (blue) indicate higher expression in adjacent neurons relative to synaptically-connected neurons. Only gene pairs with a log fold change in expression > 0.2 in either direction are displayed. Black arrows indicate AIA and AIY, and red arrows indicate two gene pairs with the high log fold changes in synaptic neurons for AIA and AIY. B) Heatmap as in A, but for CAM usage in postsynaptic connections for each neuron in the nerve ring. Black arrows indicate AVM and ALM, red arrows indicate gene pairs with high log fold change in synaptic connections relative to adjacent neurons for both AVM and ALM, and blue arrows indicate gene pairs with high log fold change in adjacent neurons compared to synaptically connected neurons for both AVM and ALM. Related to Figure 9.

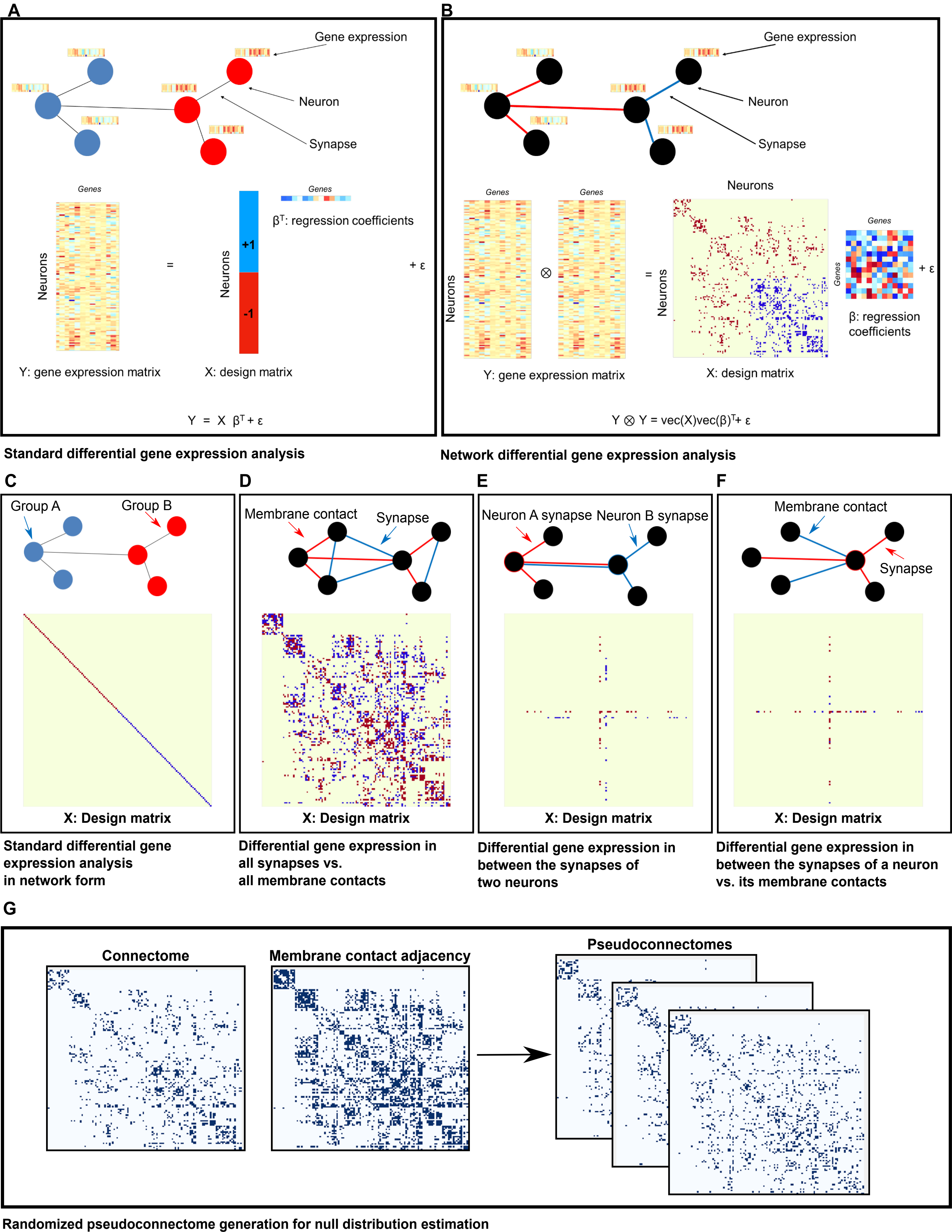
A
Gene expression
Neuron
Synapse
Genes
Neurons
=
Neurons
+1
-1
Genes
β^T: regression coefficients
+ ε
Y: gene expression matrix
X: design matrix
Y = X β^T + ε
Standard differential gene expression analysis
B
Gene expression
Neuron
Synapse
Genes
Genes
Neurons
Neurons
⊗
=
Neurons
Genes
Genes
β: regression coefficients
+ ε
Y: gene expression matrix
X: design matrix
Y ⊗ Y = vec(X)vec(β)^T+ ε
Network differential gene expression analysis
C
Group A
Group B
X: Design matrix
Standard differential gene expression analysis in network form
D
Membrane contact
Synapse
X: Design matrix
Differential gene expression in all synapses vs. all membrane contacts
E
Neuron A synapse
Neuron B synapse
X: Design matrix
Differential gene expression in between the synapses of two neurons
F
Membrane contact
Synapse
X: Design matrix
Differential gene expression in between the synapses of a neuron vs. its membrane contacts
G
Connectome
Membrane contact adjacency
Pseudoconnectomes
Randomized pseudoconnectome generation for null distribution estimation

**Figure S19. Network differential expression methods.** A) Overview of standard differential gene expression analysis (DGE): DGE can be represented as a regression model with the response variables as the neuron-wise gene expression data and the design matrix as 1s and -1s denoting the group memberships of neurons. B) Schematic of network differential gene expression analysis (nDGE): nDGE is a generalization of DGE that represents the pairwise co-expression of all genes in all pairs of neurons as the response variable. The design matrix is a square matrix of all pairs of neurons with 1s and -1s placed in locations where pairs of edges of the network are contrasted. C) Design matrix for nDGE that yields identical results as standard DGE. D) Design matrix for nDGE that explores the global genetic differences between synapses and non-synaptic membrane contact. E) Design matrix for nDGE that explores the differential gene co-expression differences between the synapses of two different neurons. F) Design matrix for nDGE that aims to demonstrate gene co-expression enrichment in synapses of a particular neuron compared to non-synaptic membrane contacts. G) The statistical significance of nDGE test statistic is obtained by computing the mean and variance of the null distribution of the test statistic resulting from pseudoconnectomes that respect the network topology of the original connectome**.** Related to Figure 9.
